# Plasticity of the primary metabolome in 241 cold grown *Arabidopsis thaliana* accessions and its relation to natural habitat temperature

**DOI:** 10.1101/2020.09.24.311092

**Authors:** Jakob Weiszmann, Pieter Clauw, Joanna Jagoda, Ilka Reichardt-Gomez, Stefanie Koemeda, Jakub Jez, Magnus Nordborg, Dirk Walther, Thomas Nägele, Wolfram Weckwerth

## Abstract

In the present study, 241 natural accessions of *Arabidopsis thaliana* were grown under two different temperature regimes, 16 °C and 6 °C, and growth parameters were recorded together with metabolite profiles to investigate the natural variation in metabolic responses and growth rates. Primary metabolism and growth rates of accessions significantly differed between accessions and both growth conditions. Relative growth rates showed high correlations to specific metabolite pools. Metabolic distances based on whole metabolite profiles were built from principal component centroids between both growth setups. Genomic prediction using ridge-regression best linear unbiased prediction (rrBLUP) revealed a significant prediction accuracy of metabolite profiles in both conditions and metabolic distances, which suggests a tight relationship between genome and primary metabolome. GWAS analysis revealed significantly associated SNPs for a number of metabolites, especially for fumarate metabolism at low temperature. A highly significant correlation was observed between metabolic distances and maximum temperature in the original growth habitat between January and March. Inverse data-driven modelling revealed that metabolic pathway regulation and metabolic reaction elasticities distinguish accessions originating from warm and cold growth habitats.

## Introduction

Acclimation and adaptation of metabolism to a changing environment are key processes for plant survival and reproductive success. The multitude of different abiotic and biotic stressors requires plant metabolism to be highly flexible, as the mode of reprograming of metabolism depends on the type and strength of stress that plants are exposed to ^1,2^. The metabolic response to changing environmental factors differs significantly between plant species ^3^, as well as among ecotypes or cultivars of the same species ^4–6^. Temperature affects plant development and has been shown to be an important determinant for the geographical distribution range of many temperate plant species, e.g., *A. thaliana* ^7^. Considering that only 5 % of the land mass worldwide is free of freezing events ^8^ and low temperature damage leads to significant losses in agricultural yield ^9,10^, the investigation of plant cold response bears a large potential in establishing a sustainable supply of food for a growing world population ^11^.

Exposure to low temperature immediately affects plant metabolism by reducing enzymatic reaction rates, which has a significant effect on biosynthesis, degradation and transport processes (see, e.g., ^12^). Within a process termed cold acclimation, metabolism is adjusted to low temperature, which, in many temperate plant species, results in increased freezing tolerance ^13^. Cold acclimation is a multigenic process that affects hundreds of genes, numerous signalling cascades and metabolic pathways to stabilize photosynthetic capacity and plant performance ^14^. Cold exposure typically results in a rapid increase of the C-repeat binding factor (CBF) transcription factors (TFs) that regulate more than 100 genes, the so-called CBF regulon, that plays a dominant role in cold acclimation ^15^. Comparing Italian and Swedish *A. thaliana* accessions revealed lower induction of the CBF regulon in the Italian accessions, which contributed to lower freezing tolerance compared to the Swedish accessions ^16^. Although CBF TFs rapidly increase after cold exposure, comparison of time-resolved cold response of *A. thaliana* revealed a faster metabolic response when compared to transcriptional response ^2^. This finding indicates a complex mode of regulation, which, in addition to transcription, also includes translational, post-translational and metabolic regulation ^17^

*A. thaliana* inhabits a large latitudinal range ^18^, and is therefore confronted with a wide range of climatic conditions. This wide distribution and the predominantly selfing reproduction type have led to the development of a large number of genetically distinct (homozygous) inbred lines called accessions, which are well adapted to the prevailing microclimate ^19–21^. The accessions feature large variances in cold and freezing resistance, acquired after cold acclimation and naïve, without cold acclimation. These adaptions were shown to be connected to the mean minimum temperature of origin, indicating selective pressure by the ability to adapt to low temperatures ^22,23^. The variance in freezing tolerance along geographical clines of origin, were correlated to several differences in the accumulation of sugars and the expression of a number of CBF-regulated genes, after an acclimation phase at low, non-freezing temperatures ^24^. A further example of the adaptation of *A. thaliana* to local climates was recently given, by showing a strong connection of climate of origin and the life-history strategy, i.e. the prevalence of winter or summer annuality ^25^.

It has been reported earlier that habitat temperature of natural *A. thaliana* accessions determines the response of physiological parameters like photosynthesis and transpiration to growth temperature ^26^. Although it is known that photosynthesis needs to be tightly linked to carbohydrates and primary metabolism in order to sustain growth and development, it remains unclear how natural variation of primary metabolism relates to growth rates. In this study, natural variation of growth rates of *A. thaliana* was monitored together with dynamics of primary metabolites under moderate (16 °C) and low (6 °C) temperature. In total, 241 natural accessions were analysed growing for three weeks under each condition.

## Results

### Natural variation allows for genomic prediction of metabolome plasticity and metabolic distance between 6 °C and 16 °C growth conditions

Absolute metabolite amount was quantified from leaf material of *A. thaliana* accessions, comprising 37 primary metabolites of which 18 changed at least two-fold and significantly in their amount (ANOVA, p<0.05) between the two different growth temperature regimes, i.e., 16 °C and 6 °C (Figure 1). Metabolite profiles differed in an accession specific manner. Most of significantly changed metabolites (15) accumulated in the plants grown at 6 °C, and only spermidine, ornithine, and glycine accumulated to higher amounts in plants grown at 16 °C. Strongest accumulation in the cold growth condition with fold changes > 45 was observed for raffinose and galactinol. Principal component analysis (PCA) separated the two growth conditions, and the two first principal components (PCs) together covered 48.52 % of total variance (Figure 2). Highest factor loadings separating 6 °C from 16 °C were found for carbohydrates and alcohols, e.g., raffinose, galactinol, sucrose, trehalose, and myo – inositol (Supplemental Table 2). Genomic prediction was performed applying the Best Linear Unbiased Predication (BLUP) methodology ^27^ and a strong predictability of metabolite profiles could be shown (Figure 3a). 25,826 unique SNPs were used to predict the 37 metabolites in both growth conditions, as well as a cross condition approach in which the metabolite profiles of the 16 °C condition were utilized to predict the metabolite profiles of the 6 °C condition and *vice versa.* For within condition prediction, Kernel density functions of predictability, scored by Pearson correlation of observed versus predicted values, peaked at a correlation coefficient of ~0.5 for the 16 °C condition and ~0.4 for the 6 °C condition. Cross-condition prediction accuracy was slightly lower (Figure 3a). Metabolic distances were calculated for all accessions to investigate natural variation of metabolic responses to the cold growth conditions. Metabolic distance values represented Euclidian distances in the PCA space covering the divergence of metabolism between 6 °C and 16 °C. Distances comprised information about all quantified metabolites and, therefore, allowed insight into the amplitude of changes on a large part of plant primary metabolism between different conditions. rrBLUP was used for prediction of metabolic distance. As shown in figure 3a metabolic distance was predicted with a slightly better correlation coefficient than the average of individual metabolites in the metabolite profile, indicating that the amplitude of change to environmental perturbation is closely related to genome variation. To investigate the genetic background of differences metabolism, GWAS was conducted using the metabolite levels in both conditions (Supplemental Table 5). The strongest, significant correlation was found for SNPs in the promotor region of the FUM2 gene (AT5g50950) in the 6 °C condition, which highlights the influence of genetic variation in the regulation of fumarate metabolism under the applied growth condition (Figure 3 b and c).

**Figure 1.**
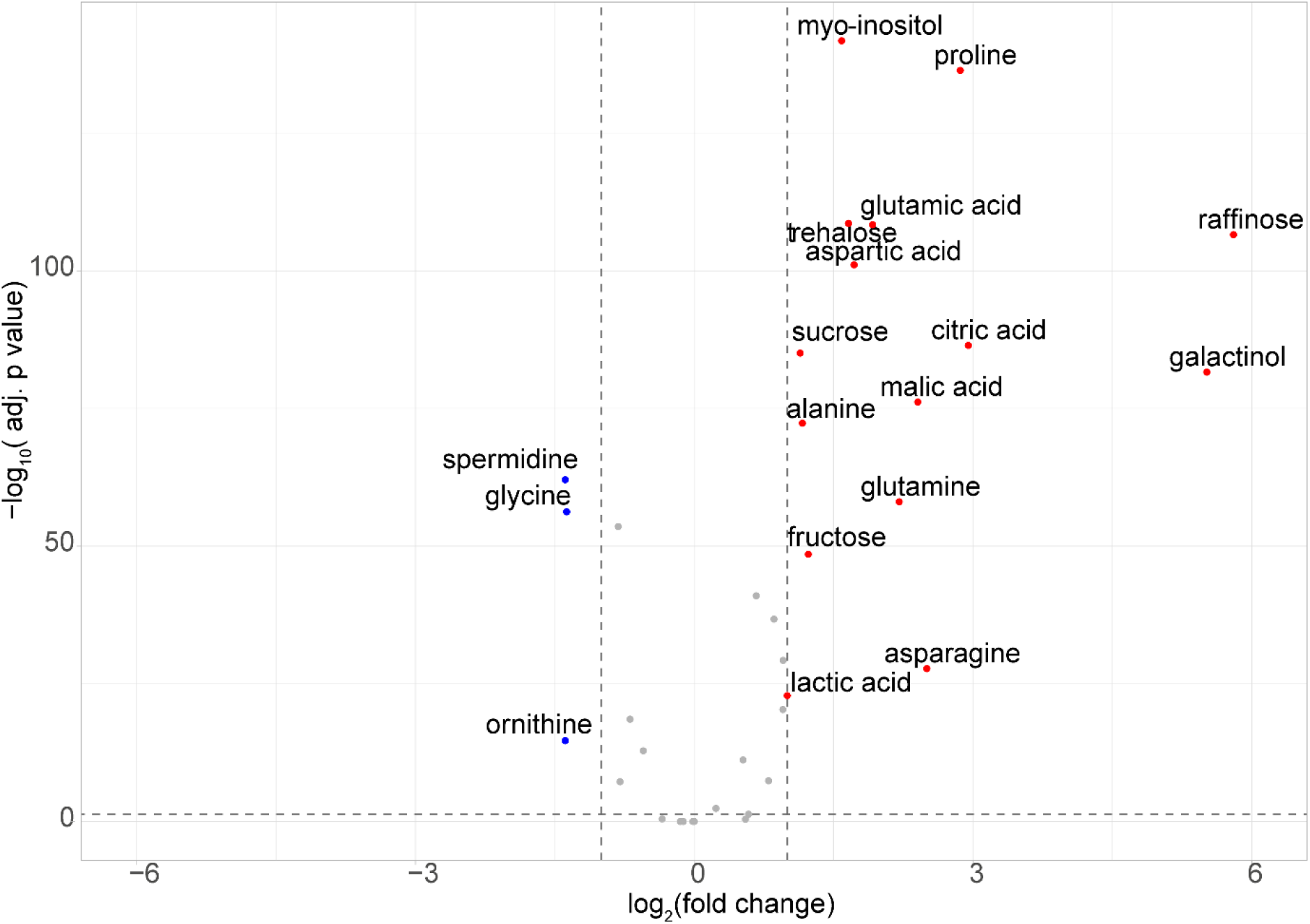
Volcano plot of targeted GC-MS data, depicting fold changes and significance of difference (p-values calculated by ANOVA, adjusted with Bonferroni correction) of metabolites between the 16 °C and the 6 °C growth condition (ratio c(6 °C)/c(16 °C)). Red dots depict metabolites with fold change ≥ 2 (≥ 1 on log_2_ scale) and p-value ≤ 0.05 (≥ 1.3 on negative log_10_ scale). Blue dots depict metabolites with fold change ≤ 0.5 (≤ −1 on log_2_ scale) and p-value ≤ 0.05 (≥ 1.3 on negative log_10_ scale).

**Figure 2.**
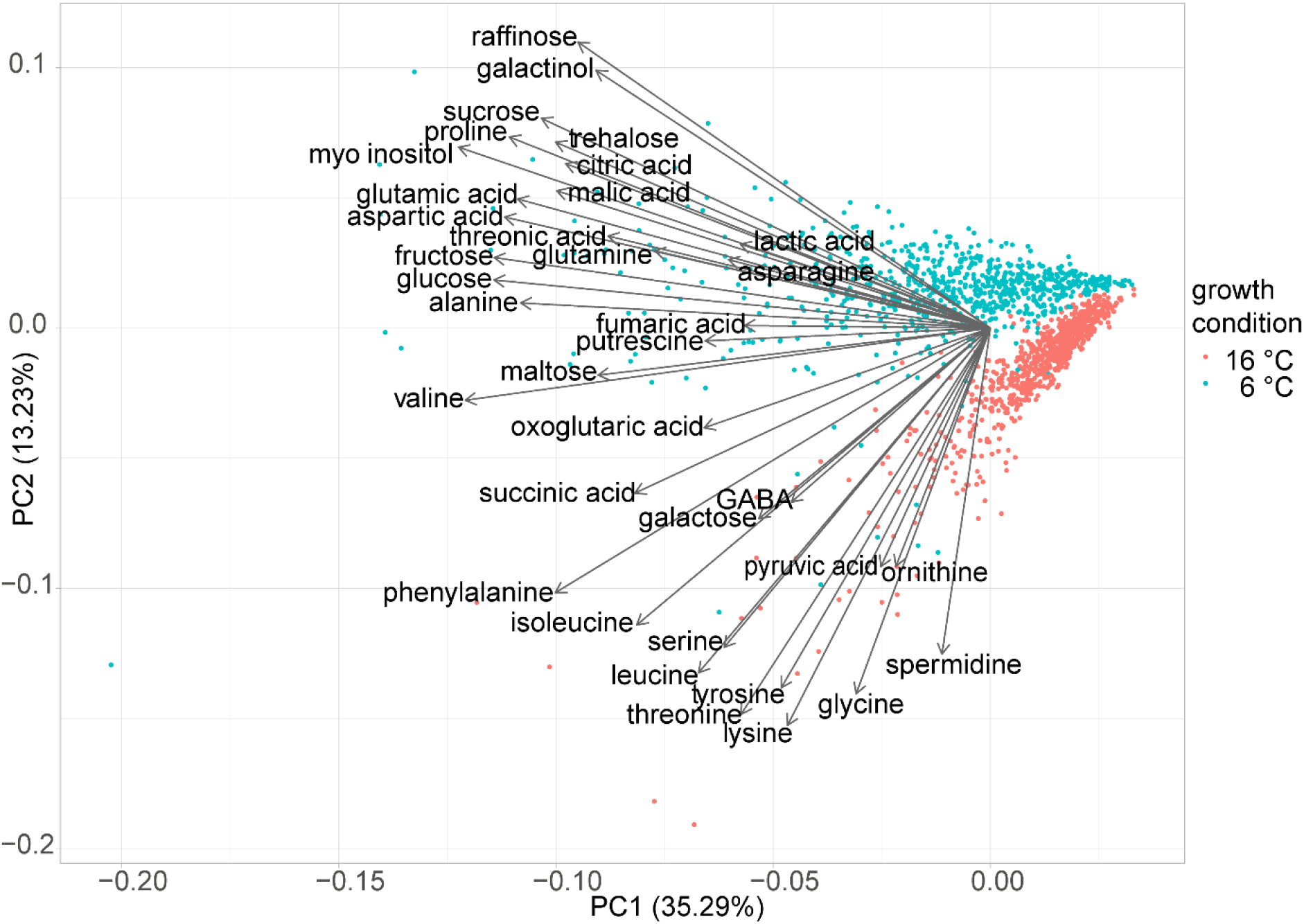
Principal Component Analysis (biplot) of targeted GC-MS data. Red dots represent samples grown at 16 °C; blue dots represent samples grown at 6 °C, arrows denote loadings of metabolites on PC1 and PC2, respectively.

**Figure 3.**
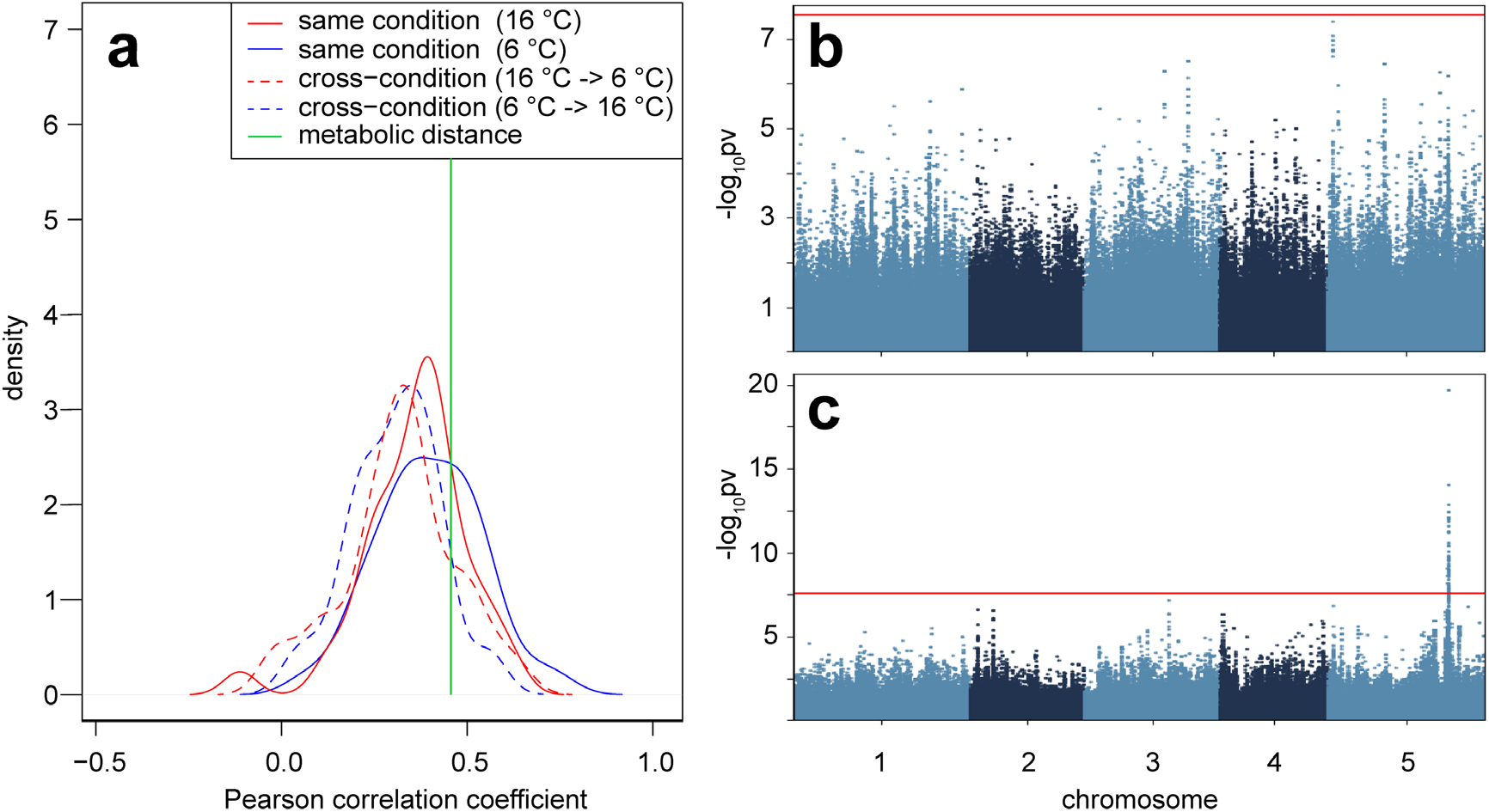
a- Prediction accuracy of genomic prediction by BLUP shown as kernel density functions of Pearson correlation coefficients of predicted versus measured concentrations of 37 metabolites. Solid lines show accuracy of predictions based on a subset of the same condition (red – 16 °C, blue – 6 °C) and dashed lines show predictions based on a subset of the other condition (red – subset of the 16 °C metabolite profiles predicting 6 °C profiles; blue – a subset of 6 °C profiles predicting 16 °C profiles). The green line shows the prediction accuracy for the metabolic distance which is the overall change from 16 °C to 6 °C growth conditions (for more details see text). b- mGWAS of fumarate concentration in the 16 °C condition, red line indicates significance threshold after Bonferroni correction. c- mGWAS of fumarate concentration in the 6 °C condition, red line indicates significance threshold after Bonferroni correction, the peak in chromosome 5 corresponds to the FUM2 gene.

### Q1 temperature at the natural origin of *Arabidopsis* accessions is linked to metabolic distances between cold and warm growth conditions

Accessions showed large diversity in metabolic adjustments to the cold growth conditions, reflected by a large range of metabolic distances (Figure 4). For southern accessions, a relatively small metabolic distance was observed, while northern accessions showed relatively large metabolic distances. Each of the accessions was assigned to a genetic admixture group ^28^ and a one-way ANOVA revealed significant differences in metabolic distance between the groups (one-way ANOVA p-value: 9.54E-13). A trend along a gradient of latitude of origin of the admixture groups was revealed (Figure 4b & Figure S 1) indicating a directed influence of genetic and geographic origin on the metabolic response to cold growth conditions. As the metabolic distance roughly correlated to a north – south gradient, a dataset containing climatic variables was used to find correlations between the climate of origin for the analysed accessions and their metabolic response. To investigate the relationship of metabolic response to cold and climate of origin, Spearman correlation coefficients between the metabolic distance and environmental variables, comprising temperature, solar radiation, water vapour pressure, precipitation and wind speed were calculated. Highest correlation coefficients for metabolic distance were observed for temperature variables between January and March, i.e., the first quarter of the year (Q1). Correlation coefficients between metabolic distance and values of climate parameters for each month revealed that temperature was the most influential parameter throughout the year, but the correlation strength decreased in the warmer part of the year, i.e., between May and August (Figure 5, Table S 1). Precipitation had a neglible correlation with metabolic distance, indicating a low impact of this factor on regulation of metabolic reactions of plants to the applied cold growth conditions. To investigate if a combination of climate variables could yield a better explanatory model for metabolic distance, backwards stepwise linear regression, selecting for the lowest RMSE (root mean square error) was employed. Using a dataset containing summary variables of climate parameters for each quarter of the year, the model with the lowest RMSE contained only the variable describing the average maximum temperature of Q1. Additional statistical analyses using different methods for variable selection, i.e., ridge regression, lasso regression and partial least square discriminant analysis (PLS-DA) confirmed that Q1 temperature was consistently the strongest predictor for metabolic distance. Additionally, stepwise backwards linear regression on a dataset containing climate variables for each month, as well as a dataset of bioclimatic variables, yielded models containing temperatures in Q1 as the most influential independent variable. Evaluation of a linear model of metabolic distance and maximum temperature in January to March showed a significantly negative correlation (R^2^ = 0.2687, p= <2.2E-16; Figure 6). This correlation was stronger than the correlation of metabolic distance with geographic latitude (R^2^ = 0.1861, p= 1.6E-12). The correlation of temperature at the geographical origin and the observed metabolic distance stayed significant after including a correction for population structure via a partial Mantel test using a genetic relatedness matrix as control variable (Table S 1).

**Figure 4.**
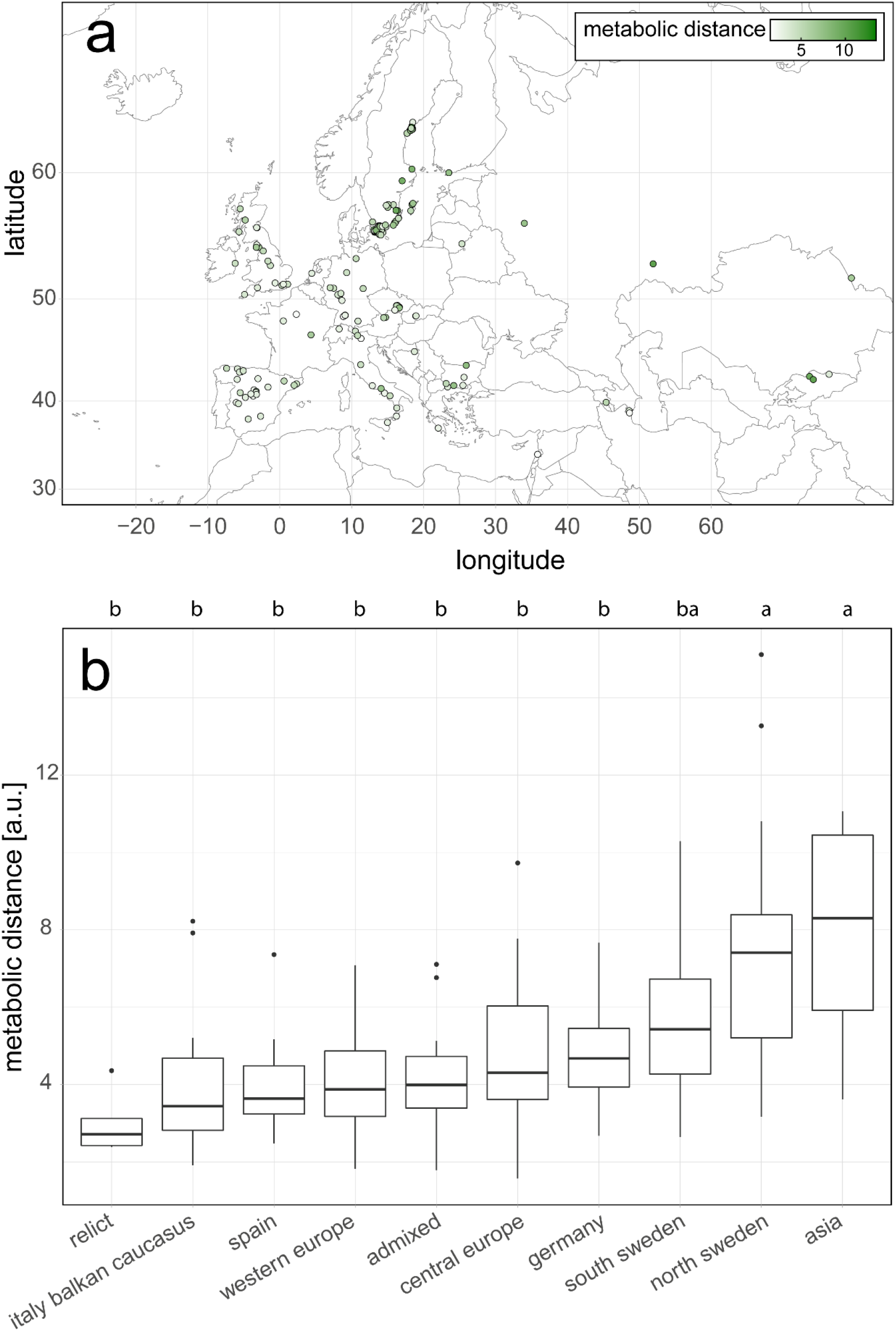
a – Map of included natural Arabidopsis accessions, colour corresponds to metabolic distance between 16 °C and 6 °C growth condition (4 Asian accessions not shown). b - Metabolic distances of Arabidopsis natural accessions, grouped by genetic admixture group. Letters denote significance groups according to ANOVA with a Tukey- HSD post-hoc test.

**Figure 5.**
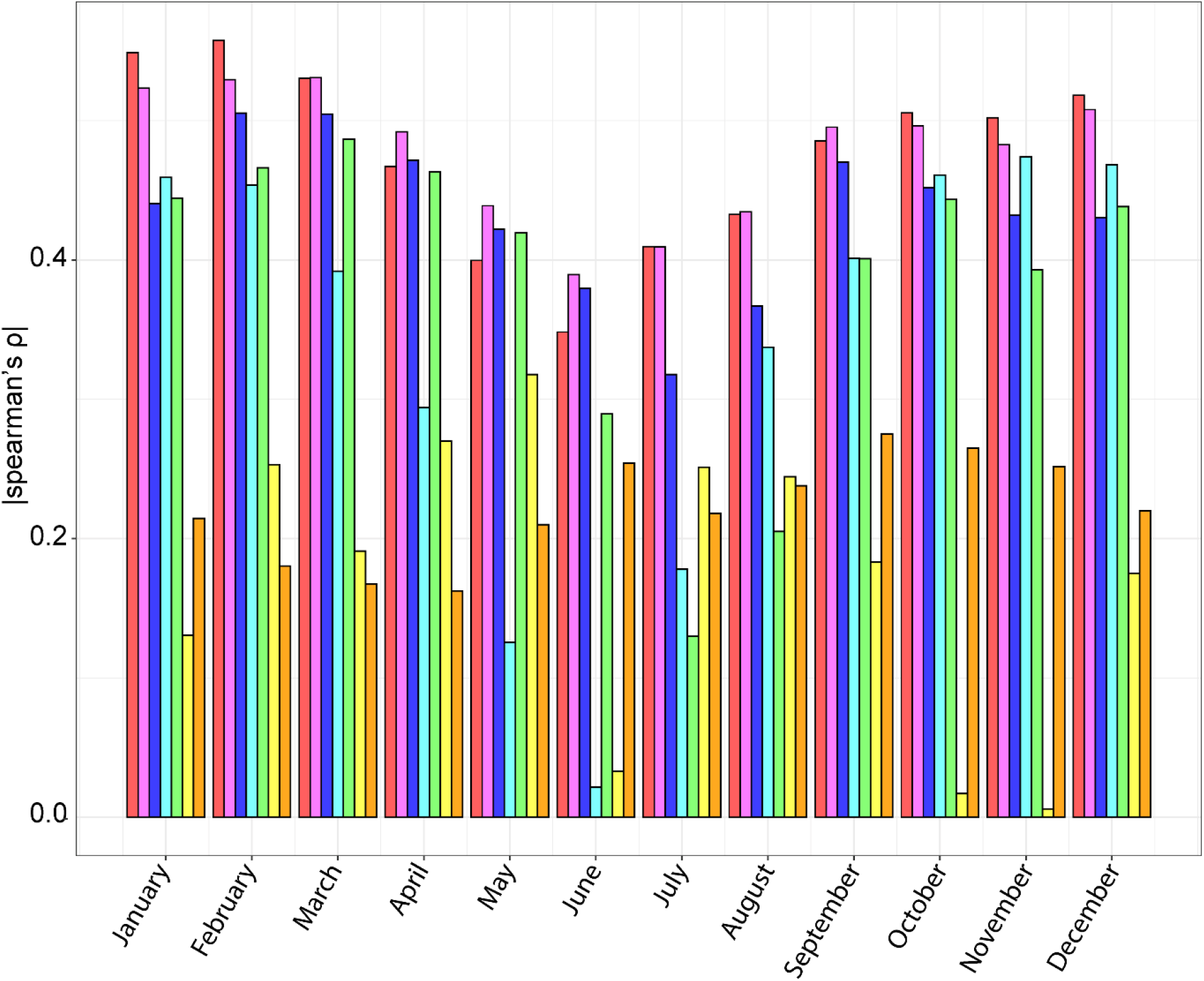
Spearman’s ρ (absolute values) describing the relation between metabolic distance and climate of origin for each month (Jan-Dec). red – temperature maximum, pink – temperature average, blue – temperature minimum, teal – solar radiation, green – water vapour pressure, yellow – precipitation, orange – wind speed

**Figure 6.**
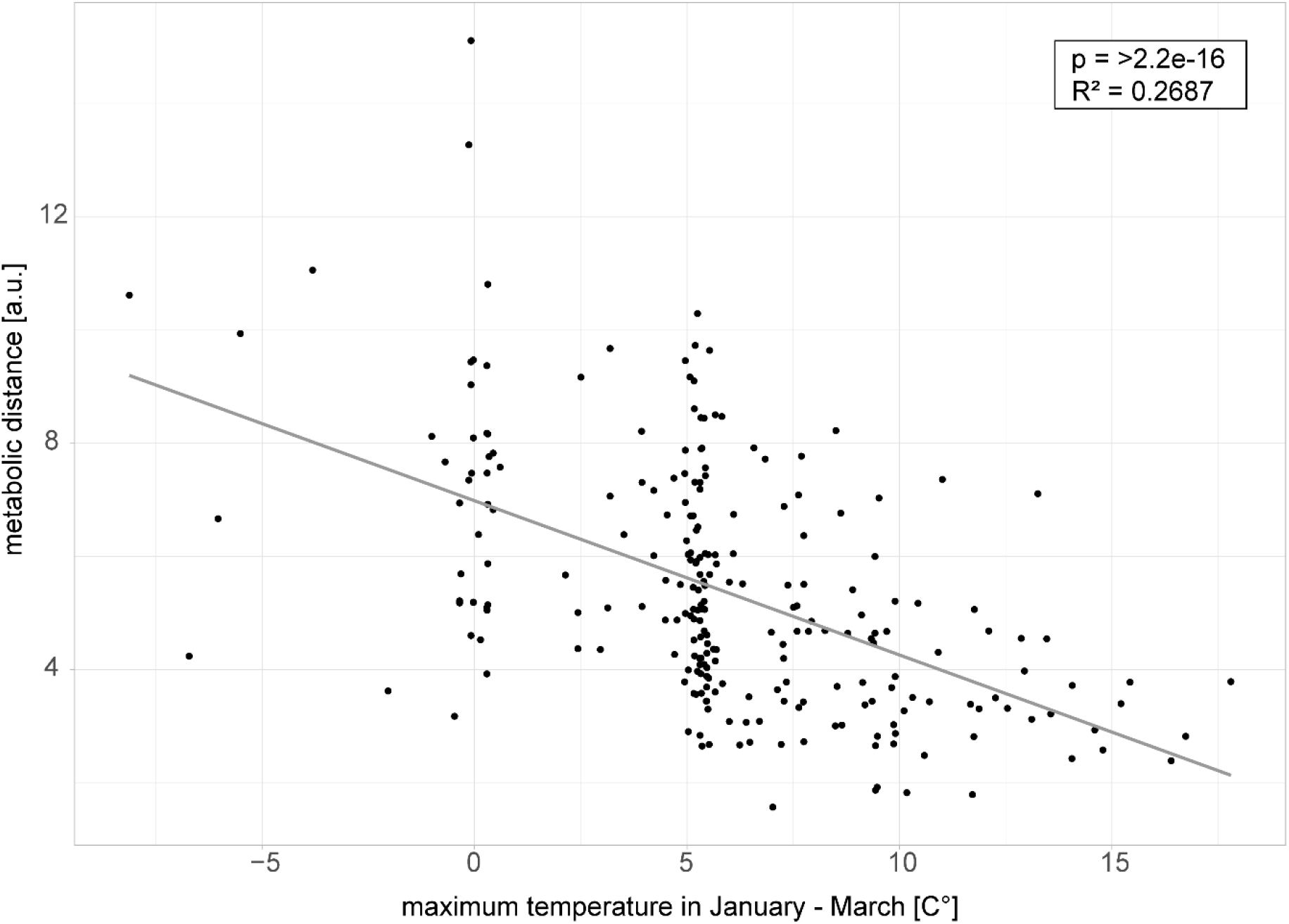
Relationship of maximum Q1 temperature (January – March) and metabolic distance (n=241). R^2^ = 0.2687, p= <2.2e- 16, Spearman’s ρ = −0.55.

### Accessions from cold and warm climate have specific metabolite profiles under cold growth conditions

Climate of origin, especially maximum Q1 temperature, was significantly correlated to metabolic distance. Therefore, two subsets of the dataset, representing accessions originating from colder or warmer climate, defined by the upper and lower quartiles (25 % and 75 %; 4.77 °C and 7.86 °C respectively) of maximum Q1 temperature (Figure S 4), were selected to investigate the differences in metabolic response to the cold growth conditions. In accessions originating from colder climate (< 4.77 °C maximum Q1 temperature), most investigated metabolites were present in higher concentration at 6 °C compared to accessions originating from warmer climate (> 7.86 °C maximum Q1 temperature). Of these metabolites, 10 were present in significantly higher concentration (one-Way ANOVA p-Value <0.05, fold change >2; Figure 7 A) and 20 were present in slightly, but significantly higher concentration (one-Way ANOVA p-Value <0.05, fold change >1 & <2). Only glutamic acid and glutamine were found at significantly higher concentrations in accessions originating from warmer climate in the 6 °C condition (one-Way ANOVA p-Value <0.05, fold change <1).

**Figure 7.**
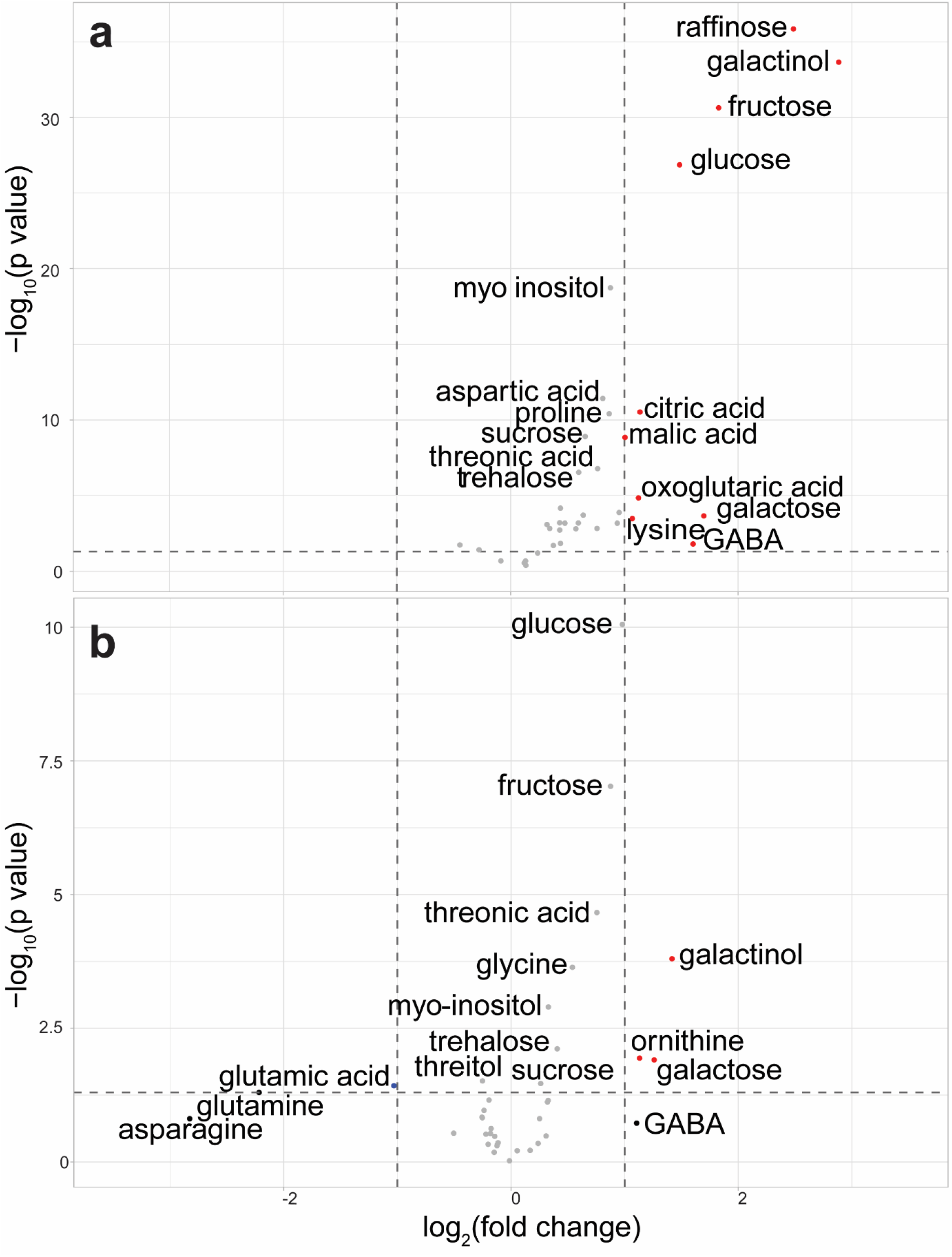
Volcano plots of differences between accessions originating from cold climate (< 4.71 °C maximum Q1 temperature) and accessions from warm climates (> 7.93 °C maximum Q1 temperature). A – Differences in absolute metabolite concentration in the cold growth condition; B- Differences in accumulation (Metabolite concentration 6 °C/ Metabolite concentration 16 °C).

Accessions originating from colder climates contained higher concentrations of sugars, and particularly of raffinose, than accessions from warm climates at 6 °C. However, raffinose was also found in higher concentration in the 16 °C growth condition in those accessions. Therefore, the proportional increase between the 6 °C and the 16 °C condition, representing the raffinose accumulation caused by the low temperature, was not significantly different between the two groups of colder or warmer origin. Glucose and fructose, on the other hand, both accumulated to higher absolute amount and in higher proportion in the accession group originating from colder climate at 6 °C (fold change ~2). In general, the accumulation of glucose and fructose was negatively correlated with maximum Q1 temperature (glc: spearman’s ρ = −0.45, p-value = 1.85E-13; frc: spearman’s ρ = −0.35, p-value = 2.19E-8).

The amount of galactinol, which is a substrate for raffinose biosynthesis, was significantly higher at 6 °C and the accumulation caused by the 6 °C condition, compared to the 16 °C condition was stronger in accessions from colder climate. Also, the amount of ornithine was higher at 6 °C and it accumulated stronger in the plants from cold climates (Figure 7).

With decreasing Q1 temperature of origin, both the average in metabolic distance and the deviation from this average increased, resulting in a higher absolute variance in metabolic distance in accessions originating from colder regions (Figure 6, Figure S 2). Therefore, some accessions from colder habitats feature a similarly small metabolic response to 6 °C as accessions coming from warmer regions, while others show metabolic distances, which are more than three times as big. The increase of variance reflected in the metabolic distance of accessions originating from colder climates was based on strong increases in the absolute variance of levels of galactinol, raffinose, threitol, fructose, glucose, citric acid, threonic acid, and proline in 6 °C (Figure 6, Figure S 2). The variance of galactinol and raffinose, described by the Full Width at Half Maximum (FWHM) of kernel density functions, was higher by a factor of ~4.5 and ~6.5 respectively, in the accessions originating from colder climates.

To test to what extent genetic variance was explaining the observed variation in metabolite levels, broad-sense heritabilities were calculated for each metabolite in both temperatures. These heritabilities ranged from close to zero to 0.52 and 0.43 in respectively 16 °C and 6 °C. Four metabolites of those that lead to increased variance in metabolic distance had the highest heritabilities in 6 °C: galactinol, raffinose, fructose, glucose (Figure S 3). The highest heritabilities in both temperatures were found for galactinol and raffinose. This shows that there is a genetic basis for the observed variance in metabolites. With a mixed-effect model we tested for significance of the genotype specific temperature response (genotype by environment interaction; GxE) of each metabolite. Eleven metabolites showed a significant GxE effect (citric acid, fructose, galactinol, glucose, malic acid, myo inositol, oxoglutaric acid, proline, raffinose, serine, and trehalose; fdr < 0.05). Together, this shows that there is a genetic basis for both the variation of certain metabolites within specific temperatures, as for the temperature response itself.

### Inverse data driven modelling indicates different regulation of amino and organic acid metabolism as well as sucrose cycling between accessions originating from warm and cold climates

To investigate regulation of metabolism, reaction elasticities were calculated based on a method for inverse data driven modelling, which connects metabolite variance information with a metabolic network ^29,30^. Metabolite data from the 6 °C condition of two subsets representing cold and warm Q1 climate were used for this approach. Calculations resulted in the biochemical Jacobian matrix, representing rate elasticities for both groups. Inverse approximation was performed in five independent replicates using different threshold values for the definition of cold and warm origin accessions (Figure S 4). To find the strongest differences in Jacobian entries between accessions originating from warm and cold regions, reaction elasticities, i.e., entries of the Jacobian matrix, were statistically analysed in a PCA (Figure 8). This analysis showed a clear separation between the two groups of origin on PC1 (>50 %) indicating strong differences in the biochemical regulation in response to the cold growth condition. Absolute values of loading scores for the individual Jacobian matrix entries for PC1 were listed according to their influence on separation (Figure 9). Strongest changes in reaction elasticities were observed in fumarate metabolism, amino acid biosynthesis, sugar cleavage, and branched chain amino acid (BCAA) metabolism.

**Figure 8.**
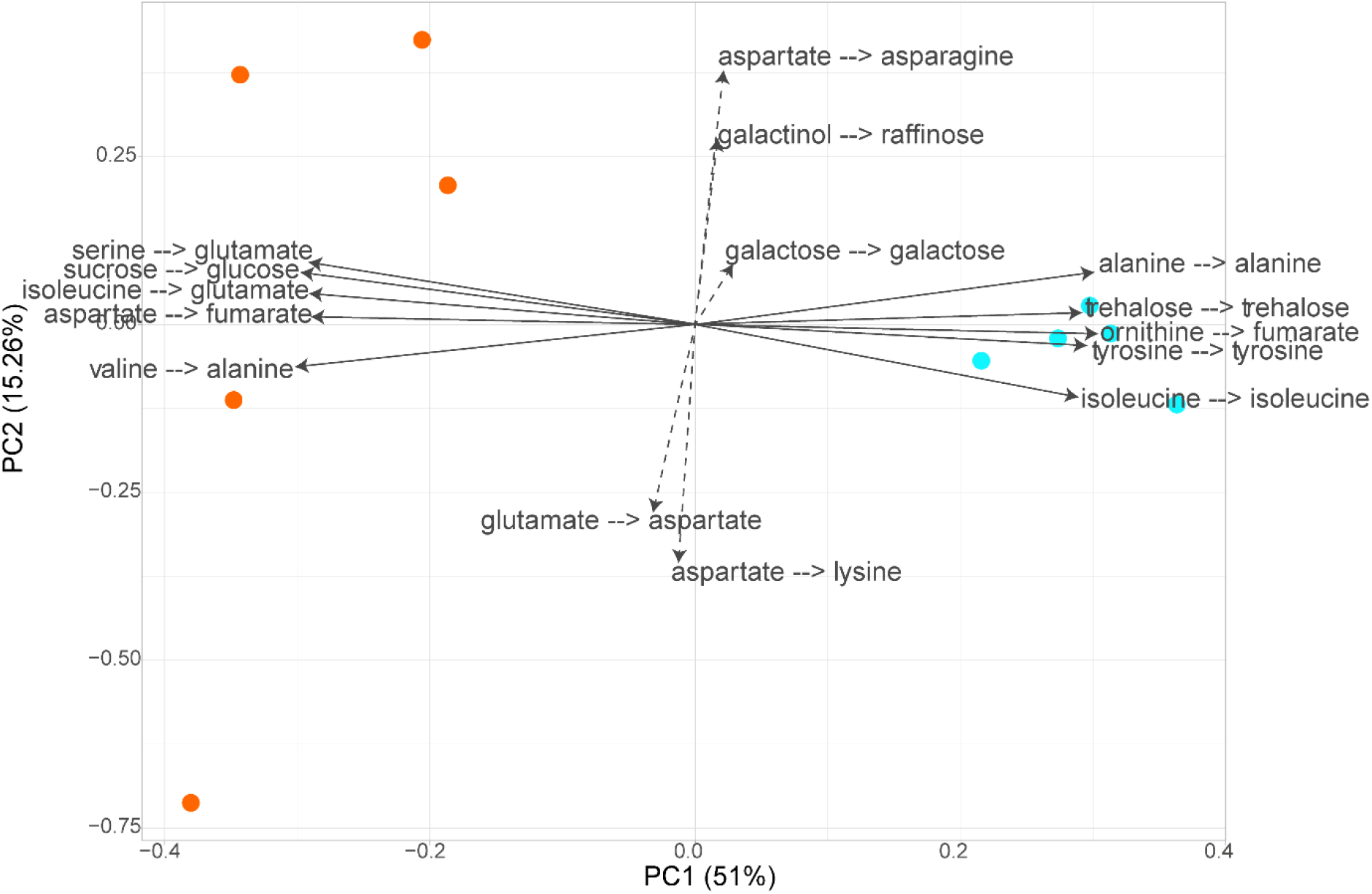
Principal component analysis (PCA) of Jacobian matrix entries of accessions from cold (teal) and warm (orange) origins. Five Variations of quantile threshold in definition of cold and warm are depicted ((20 %, 80 %; 23 %, 77 %; 24 %, 76 %; 25 %, 75 %; 26 %, 74 %). Jacobian matrices were calculated from covariance matrices based on metabolite data from plants grown in the 6 °C condition. Depicted are the 10 strongest loadings (lines) and the 5 weakest loadings (dashed lines) for PC, X --> Y = Metabolic function Y depending on metabolite X 1. Orange – warmer origin, teal – colder origin.

**Figure 9:**
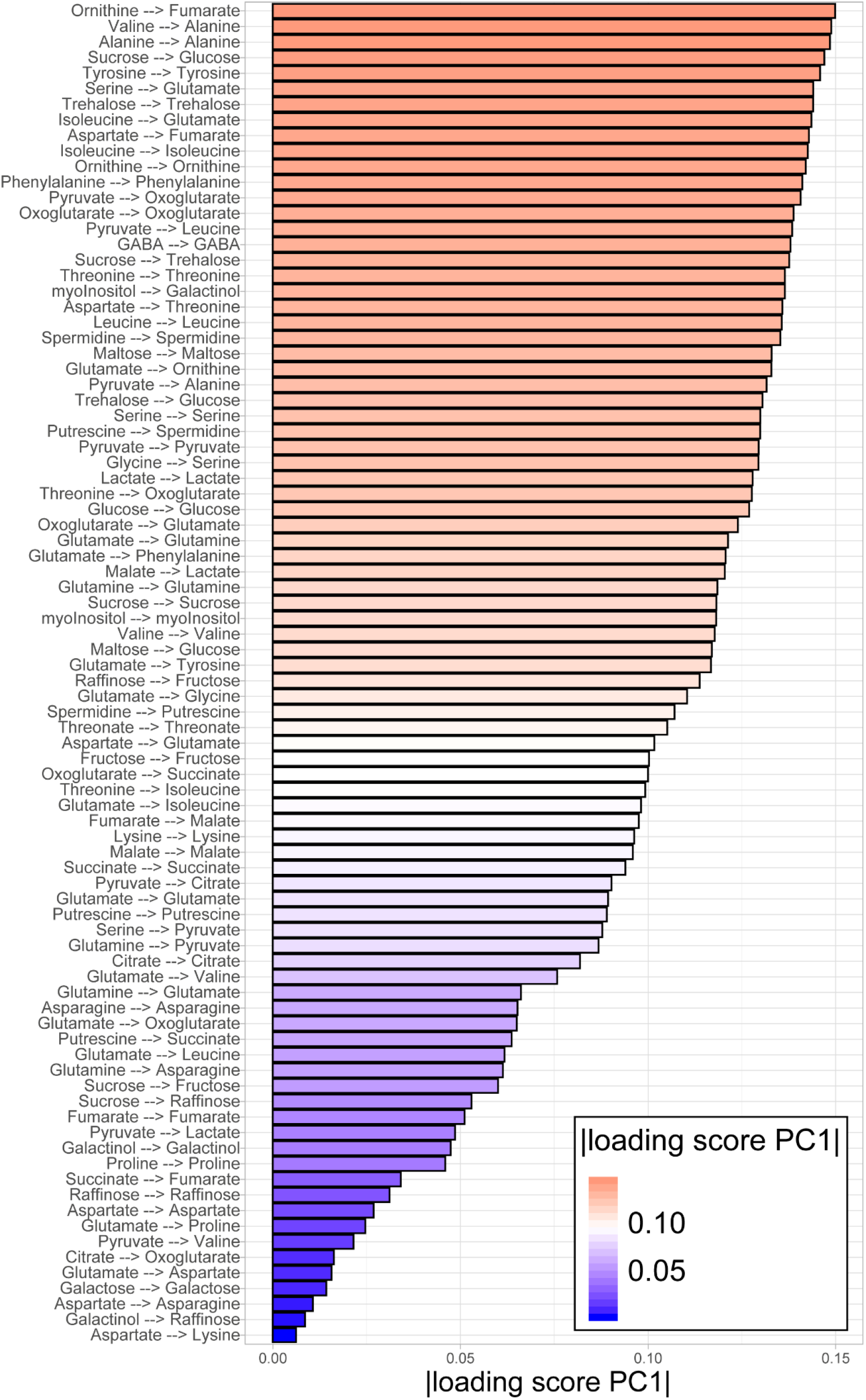
Absolute loading scores of Jacobian matrix entries, representing the contribution to separating the two origin groups on PC1; X --> Y = Metabolic function Y depending on metabolite X

Jacobian entries related to fumarate metabolism had a strong influence on the separation of the two origin groups, pointing to a differential regulation of fumarate metabolism under low temperature, which is further highlighted by the strong correlation of SNPs in the promotor region of FUM2 with fumarate levels in the 6 °C condition (Figure 3c).

Reaction elasticities for glucose, which were related to sucrose cleavage, were different in the 6 °C condition between the two origin groups. This, together with the link of hexose accumulation to Q1 climate of origin, pointed to different strategies in the central sucrose metabolism. Furthermore, SNPs in 6 invertase genes (AT1G35580, AT3G13784, AT1G12240, AT3G13790, AT1G62660, AT3G52600, AT4G09510) showed strong correlation with temperature of origin in the first months of the year in an analysis performed with the online tool GenoCLIM ^31^.

Reaction elasticities in raffinose metabolism, represented by the dependency of raffinose on galactinol levels (galactinol --> raffinose) were observed to be very similar between the two origin groups, indicated by a low loadings-score in the principal component analysis (Figure 9). This finding reflected similar raffinose accumulation rates in both groups (Figure 7b). Even though the total concentrations of raffinose were significantly different (Figure 7a), the sensitivity of the respective metabolic system seemed to be very similar in the prevalent scenario.

Reaction elasticities related to alanine metabolism showed strong differences between the two origin groups (Figure 9), despite no apparent differences in accumulation or absolute concentration (Figure 7). Likewise, high loading scores for Jacobian entries of valine and isoleucine, i.e., branched chain amino acids (BCAAs), indicated its importance in cold acclimation strategies. Like for alanine, BCAAs did not differ significantly in their absolute concentration and accumulation rate between the origin groups (Figure 7). Generally, valine, isoleucine and alanine concentration did not contribute strongly in separating the growth conditions (6 °C and 16 °C) in the PCA based on the metabolite measurements (Figure 2), but rather added to the variance within the conditions, pointing to a high variance in these metabolites between the individual accessions, as well as to differences in the variance between accessions from colder and accessions from warmer climates.

### Relative growth rate was connected to different metabolite pools under cold growth conditions

To investigate correlations between metabolite levels and overall growth rates in both growth conditions, stepwise linear modelling was used to find the strongest connection between metabolism and growth. In the 6 °C growth condition, the resulting model contained phenylalanine, raffinose, serine, spermidine, and trehalose (R^2^ = 0.26, p = 2.64e-15). When these correlations were investigated one by one, phenylalanine, spermidine, and trehalose correlated positively and raffinose and serine correlated negatively with overall growth rate. In the 16 °C condition, the model with the lowest RMSE contained asparagine, glycine, pyruvic acid, serine, spermidine, and trehalose (R^2^ = 0.49, p < 2.2e-16). In this case, glycine, spermidine, and trehalose correlated positively and asparagine, pyruvic acid and serine correlated negatively with overall growth rate. Comparing the two models, three metabolites (serine, spermidine and trehalose) correlated with the growth rate in both conditions, indicating a general connection of growth with these metabolites, while phenylalanine and raffinose occurred only in the model for 6 °C and asparagine, glycine and pyruvic acid were only contained in the model for growth in the 16 °C condition.

## Discussion

### Natural habitat temperature in the first quarter of the year predicts the response of primary metabolism in cold grown plants

In previous studies, it has been described that freezing tolerance of cold acclimated plants significantly correlates with latitude of geographical origin of natural *Arabidopsis* accessions ^24,32^. In the present study is was shown that the extent of primary metabolism response to growth at 6 °C is connected to geographical latitude as well, but a correlation with climatic variables revealed a much stronger connection to the temperature between January to March. The general direction of separation between the growth conditions in a PCA was similar for all included accessions, but the metabolic response to the 6 °C condition was stronger in plants originating from colder climates. A strong connection of freezing tolerance and the temperature in January could be linked to the expression level of CBFs in four Chinese *Arabidopsis* accessions ^33^. Interestingly, no direct relation between genetic relatedness and freezing tolerance after acclimation was observed ^24^, which strongly suggests that local adaption to climate is the key driving factor for the cold response and freezing tolerance in *A. thaliana* ^34,35^. Similarly, it has recently been shown, that grapevine varieties adapted to local climate deal better with abiotic stress in a range typical for the specific climate compared to widely used commercial varieties ^36^. Climatic range boundaries of *A. thaliana* distribution were shown to be determined by a combination of temperature and precipitation ^7^, which can explain the connection of temperature in Q1, and metabolic reaction to cold. In the present study, however, no significant correlation of metabolic distance and precipitation at the original habitat was detected. This lack of correlation suggests that, even though both, temperature and precipitation, limit the distribution range, the metabolic response to these factors is uncoupled from each other. Additionally, the correlation of metabolic distance with original habitat temperature was less significant in the warmer part of the year, i.e., between May and August (Figure 5). This allows us to speculate that the extent of metabolic response to low ambient temperature underlies selective pressure only by the temperature in the cold part of the year, even though mean temperatures throughout the year also correlate to a strong extent. The increase in metabolic distance with decreasing Q1 temperature was predominantly driven by strong accumulation of raffinose, galactinol, fructose, glucose, citrate and malate (see Figure 7). Soluble sugars have been shown earlier to positively correlate with the capability of natural accessions to acclimate to low temperature and to increase freezing tolerance ^22^. However, plant development and growth under low temperature results in a different metabolic constitution than observed for plants which were shifted from ambient to low temperature in mature stage for cold acclimation ^37^. In particular, *Arabidopsis* leaves which developed at 5°C accumulated relatively high amounts of soluble sugars but, in contrast to cold shifted plants, released suppression of photosynthetic genes which the authors discussed to be essential for development of full freezing tolerance ^37^. Although photosynthetic parameters were not quantified in the present study, correlation of metabolites with growth rates at 16 °C and 6 °C revealed a significantly positive correlation with spermidine and trehalose while phenylalanine only correlated positively under 6 °C. Interestingly, spermidine and trehalose have previously been found to correlate positively with growth under 20°C while phenylalanine correlated negatively under these conditions ^38^. Following the discussion of Meyer and colleagues, who interpreted growth-correlated metabolites as positive (or negative) signals, this would indicate that also under low temperature spermidine and trehalose represent conserved growth signals. Consequentially, due to its negative correlation under ambient temperature ^38^ and a missing correlation with growth rate at 16 °C, phenylalanine would represent a cold-specific growth signal. Phenylalanine represents a central metabolic precursor for numerous secondary metabolites, e.g., flavonoids and lignin ^39,40^. Hence, in contrast to spermidine and trehalose, which were rather discussed as pure growth signals than growth substrate molecules ^38^, phenylalanine might play a more complex role and might serve as a central metabolic integrator for growth, development and stress protection under low temperature.

The metabolic response was predictable from genetic variation among the investigated genotypes (Figure 3a). Interestingly, the predictability improved gradually from the 16°C to the 6°C growth condition and showed the best predictability for metabolic distance (Figure 3 a). Metabolic distance comprises the sum of all metabolite perturbations from reference growth (16°C) to stressed condition (6°C). Accordingly, it is the most comprehensive and synergetic parameter for the description of the cold stress response in relation to the reference metabolome and, thus, correlates even stronger to genetic variation then individual metabolite concentrations.

### Cold-grown accessions originating from warmer and colder regions differ in the plasticity of primary metabolism

Accessions from colder climates showed a stronger variability in their metabolic response under 6 °C. This was reflected in higher variance of metabolic distances when compared to accessions originating from warmer regions. The applied 6 °C growth condition seemed to trigger a highly conserved and less variable metabolic response in accessions originating from warmer climates, which might be explained by a reduced amplitude of extremely low temperatures in the original habitat. In plants from colder climates the metabolic response to cold was generally stronger, but also more diverse, which hints at a larger number of employed metabolic strategies in dealing with cold, and potentially also freezing stress, among the different accessions. For plants from colder regions which are regularly confronted with freezing events, the strategy to invest in a stronger metabolic response to cool temperatures’, potentially preparing for even lower temperatures seems to be feasible, while plants from warmer regions react with smaller metabolic deviations when confronted with low temperatures, which indicates a strategy of trying to endure without investing too many resources in adaption.

As a consequence of differential metabolite covariances, the calculated reaction elasticities, described by Jacobian matrices, revealed strong differences between accessions from cold and accessions from warm climates. In general, Jacobian matrices allow the investigation of causal relations between metabolites and point to differences in metabolic regulation. Most pronounced differences in Jacobian entries between accessions originating from cold and warm habitats were found in fumarate metabolism, sucrose cleavage and BCAA metabolism. Under low temperature, fumarate serves as a carbon sink in leaf metabolism, aiding in the acclimatisation of photosynthesis to a new homeostasis ^41^. The differential accumulation of fumarate under stress in *A. thaliana* accessions with different cold acclimation potential could already be shown in an earlier study ^42^ and fumarase 2 (FUM2) was described to have a strong effect on carbon partition and growth rates in A. thaliana accessions ^43^. Remarkably, both absolute amount and accumulation rate of fumarate were not significantly correlated with the Q1 temperature of origin and therefore not different between the two origin groups in this study, which points to the importance of differences in reaction elasticity in this metabolic pathway, rather than absolute concentration differences in the cold response. Organic acids like fumarate play an important role in regulating the accumulation of solutes within the vacuole by controlling cold induced acidification ^44^, which makes them a good target for metabolic regulation, as indicated by the differences in reaction elasticities.

The score for the influence of ornithine on the metabolic function of fumarate additionally points to the plastidial conversion of ornithine to arginine via ornithine carbamoyltransferase (OTC), argininosuccinate synthase (ASSY), and argininosuccinate lyase (ASL). This transformation is an important part of nitrogen cycling and homeostasis ^45,46^. The connection of natural variation along the gradient of Q1 temperature of origin and this pathway is supported by the prevalence of a SNP in the ASL gene (At5g10920), which is highly correlated with temperature of origin in >1000 Arabidopsis accessions ^31^. This points to differences in the extent of regulation of amino acid metabolism caused by cold temperature between accessions originating from colder and accessions originating from warmer climate. It has been shown that amino acid metabolism and nitrogen usage have to be heavily adjusted under stress conditions to allow survival in plants ^47–49^.

Sucrose metabolism plays a central role in plant development, stress response and growth regulation, and its cyclic metabolism, composed of invertase-driven cleavage and cytosolic re-synthesis, represents an important buffer mechanism against environmental fluctuation ^50,51^ and the observed differences in reaction elasticities connected to sucrose metabolism strongly point to differences in regulation in this pathway between plants from warmer and colder habitats. While futile cycling of sucrose might stabilize the cellular energy status by providing electron acceptor molecules for photosynthesis and serve as an efficient mechanism to control carbon partitioning ^52^.

Entries of the Jacobian matrix connected to alanine metabolism had strong discriminatory loadings between the origin groups in the PCA (see Figure 8). Alanine plays an important part in amino group transfer reactions in amino acid and protein biosynthesis and has been described to accumulate during cold exposure in various plant species ^53^. Furthermore, alanine serves as amino group donor in photorespiration to replace nitrogen taken out of the cycle in form of serine or glycine ^54,55^.

The elasticities of reactions involved in BCAA metabolism, especially valine and isoleucine, are highly different between the investigated groups of origin at 6 °C. As for alanine there was no significant change in concentration, but the investigation of jacobian entries revealed that accessions from colder climates likely feature differences in regulation of this metabolic pathway, which has been shown to be strongly regulated in response to changes in environmental condition ^56,57^.

### Conclusions

Changes in ambient temperatures have different ecological implications in the different growth habitats of Arabidopsis. We could show that within a large group of accessions from diverse growth habitats, a significant connection of origin temperature from January to March and metabolic reaction to cold exists. Key pathways of primary metabolism were affected differently between accessions originating from cold or warm climates, not only in the amount of accumulated products, but also in the strategies of regulation. Furthermore, we could show that plants from colder regions employ a larger spectrum of metabolic responses to low ambient temperature.

While cool temperatures might be the signal for an upcoming frost period, implying long-term endurance for accessions coming from more northern, colder climates, in southern, warmer climates lower temperatures could rather indicate a short-term event requiring less adaption of metabolism. Thus, the observed differences in metabolic regulation while growing in cold conditions indicate two different strategies. Preparation for even more adverse, freezing conditions in accessions from colder climates, or trying to survive the current, mild stress situation while preparing for a fast recovery after the end of the cold phase in accessions from warmer climates. Additionally, the necessity of developing new strategies to deal with cool temperatures is of low importance for accessions from warm climates, resulting in low variance of metabolic responses, compared to accessions from climates, which are regularly exposed to sub-zero temperatures.

## Methods

### Experimental design and plant growth

Seeds of 241 natural accessions (Supplemental Table 3) of *A. thaliana* described in the 1001 genomes project^28^ were sown on sieved (6 mm) substrate (Einheitserde ED63). Pots were filled with 71.5 g ±1.5 g of soil to assure homogenous packing. The prepared pots were all covered with blue mats ^58^ to enable a robust performance of the high-throughput image analysis algorithm. Stratification was done for 4 days at 4 °C in the dark, upon which the seeds were germinated and seedlings established at 21 °C for 14 days (relative humidity: 55 %; light intensity: 160 μmol m^−2^ s^−1^; 14 h light). The temperature treatments were started by transferring the seedlings to either 6 °C or 16 °C. To simulate natural conditions temperatures fluctuated diurnally between 16-21 °C, 0.5-6 °C and 8-16 °C for respectively the 21 °C initial growth conditions and the 6 °C and 16 °C treatments (Fig. S X). Relative humidity (55 - 95 %) and light intensity (160 μmol m^−2^ s^−1^) were kept the same for all experiments. Daylength was 9h during the 16 °C and 6 °C treatments and 14h during the 21 °C initial growth conditions. Each temperature treatment was repeated three times. Five replicate plants were grown for every genotype per experiment. Plants were randomly distributed across the growth chamber with an independent randomisation pattern for each experiment. During the temperature treatments (14 DAS – 35 DAS), plants were photographed twice a day (1 hour. after/before lights switched on/off), using an RGB camera (IDS uEye UI-548xRE-C; 5MP) mounted to a robotic arm. At 35 DAS, whole rosettes were harvested, immediately frozen in liquid nitrogen and stored at −80 °C until further analysis. Rosette areas were extracted from the plant images using Lemnatec OS (LemnaTec GmbH, Aachen, Germany) software. Growth parameters were obtained through non-linear modelling. Plant sizes did not reach a plateau phase, therefore we opted for power-law function as described by Paine ^59^.

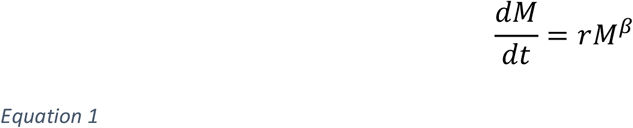

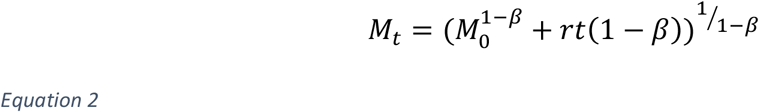

The power-law function is described with three parameters: M_0/t_ is the plant size at time point 0 or timepoint t, respectively; r is the overall growth rate; beta is scaling factor for the growth rates, letting growth rates increase or decrease with plant size. Parameter estimates for each accession were obtained in a three-step procedure. First, non-linear regression was performed for each individual plant using the nlsList function ^60^ with the power-law selfStart function (Equation 2) from Paine ^59^. In a second step, the initially obtained parameter estimates were used to define priors for Bayesian nonlinear modelling. The brm function ^61^ was used to model growth with Equation 2 for each individual plant.

priors were defined for M_0_, r and beta as follows:

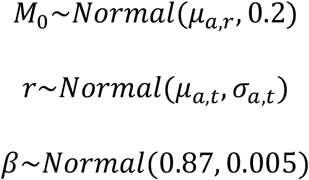

For M_0_ μ_a,r_ was defined as the median M_0_ estimate from the non-linear regression for accession a in replicate r. For the overall growth rate m_a,t_ was defined as the median r estimate from the non-linear regression for accession a in temperature t, s_a,t_ was defined as the standard deviation of the r estimates obtained from the non-linear regression for accession a in temperature t.

For every accession, estimates for each growth parameter in each temperature were obtained from a mixed model ^60^ with accession, temperature and their interaction as fixed effects and experiment as random factor. Estimates for each accession in each temperature were then calculated as estimated marginal means ^62^.

### Metabolite quantification and profiling via gas chromatography coupled to mass spectrometry (GC-MS)

Frozen leaf material was homogenized in a ball mill (Retsch GmbH, Haan, Germany). Polar metabolites were extracted and measured on gas chromatography coupled to mass spectrometry as previously described ^50^, with slight modifications. In brief, homogenized plant material was extracted with a methanol-chloroform-water mixture (MCW, 2.5/1/0.5, v/v/v) on ice for 15 min, which was split in a polar and an apolar fraction by addition of water after extraction. Subsequently, pellets were re-extracted twice using 80% ethanol at 80 °C for 30 minutes. Ethanol extracts were combined with the polar phase of the MCW extraction and dried in a vacuum concentrator (ScanVac, LaboGene, Allerød, Denmark). To compensate technical variance in the measurements, two internal standards, i.e. pentaerythritol and phenyl b-D-glucopyranoside (both Sigma-Aldrich) were spiked to the extracts before drying. Dry extracts were derivatized by methoximation for 90 min at 30°C with methoxylamine hydrochloride (Merck KGaA, Darmstadt, Germany) in pyridine and silylation for 30 min at 37°C with N-Methyl-N-(trimethylsilyl) trifluoroacetamide (MSTFA, Macherey – Nagel, Düren, Germany). Measurements were conducted on a LECO Pegasus® GCxGC-TOF mass spectrometer (LECO Corporation, St. Joseph, USA) coupled to an Agilent 6890 gas chromatograph (Agilent Technologies®, Santa Clara, USA) using an Agilent HP-5Ms column (length: 30 m, inner diameter: 0.25 mm, film: 0.25 mm). For targeted analysis, baseline correction, chromatogram deconvolution, peak finding, retention index calculation and peak area extraction were done in the software LECO Chromatof®. Retention index calculation was conducted by measuring a mixture of linear alkanes (C12-C40) with every measurement batch. All metabolites included in the targeted analysis were identified and quantified by measuring a mixture of pure standard compounds in different concentrations in every measurement batch. Areas of each metabolite were normalised to the internal standard with a minimum distance of retention time to the metabolite. Internal standard normalized areas where then normalized to the slope of peak areas of the corresponding externally measured standard row and to sample fresh weight yielding the absolute amount of metabolites [μmol gFW^−1^]. The data table containing all metabolite quantifications can be found in the supplement (Supplemental Table 4).

### Statistical analysis

Statistical analyses were conducted within the free statistical software environment R ^63^. Data manipulation, summarisation and plotting was conducted using the R package *tidyverse* ^64^.

Principal component analysis (PCA) was performed within R, after scaling and centering of metabolite data. The plot was visualised using the R package *ggfortify* ^65^. To calculate metabolic distances, multidimensional means (centroids) using the first 15 principal components (PCs) were built for each natural *Arabidopsis* accession in both conditions. Coordinates of centroids were consecutively used for the calculation of Euclidian distances between the centroid of the 6 °C and the 16 °C growth condition for each accession, representing the metabolic distances ^66^.

Spearman correlation coefficients and p-values for single correlations were calculated using the R package *Hmisc* ^67^. Climate and bioclimatic data were taken from the WorldClim Database ^68^, which was further used to calculate summary variables as three month means. The worldClim2 data was linked to each natural *Arabidopsis* accession based on longitude and latitude of their origin.

Stepwise backwards linear regression modelling and partial least square discriminant analysis (PLS-DA) were conducted within the R packages *caret* ^69^ and *leaps* ^70^, employing five times repeated 10-fold cross-validation. Model selection was based on minimizing RMSE in cross-validation (root-mean-square-error). Ridge regression, and Lasso selection were fitted with the R package *glmnet* ^71^ by splitting the dataset into training data (75 %) and test data (25 %) and selecting the penalty parameter λ (lambda) by minimising MSE (mean square error).

Population-structure-corrected correlation coefficients were calculated using the *mantel.partial* function included in the R package *vegan* ^72^ correcting the correlations with a genetic kinship matrix ^28^.

Broad sense heritabilities were calculated as the ratio between genetic variation and total phenotypic variation. Variances were estimated from a random effect model (lme function in nlme package ^60^; R), with genotype as random effect. Genetic variance was the variance allocated to the random effect ‘genotype’, total phenotypic variance was the sum of the random effect and residual variance (VarCorr function in nlme package ^60^; R).

### Data-driven inverse modelling

Calculation of Jacobian matrices was conducted as previously described ^30,73^. In brief, covariance data, calculated directly from the metabolite concentrations in the 6 °C condition for the two origin groups of *Arabidopsis* accessions, which were defined by maximum Q1 (first three months of the year) temperature of origin, under both applied growth conditions were connected with biochemical network information and used for an inverse approximation of biochemical Jacobian matrices. The calculations were repeated five times, varying the quantile threshold for the definition of the two origin groups (20 %, 80 %; 23 %, 77 %; 24 %, 76 %; 25 %, 75 %; 26 %, 74 %, upper and lower quantiles of tmax_01_02_03 temperature respectively). For each variation threshold, the inverse calculation of Jacobian matrices was conducted 1×10^4^ times and the resulting median was normalized to the inverse variance of all calculations. Each calculation was repeated 1×10^4^ times and a median was taken and normalized to the inverse variance of all calculations. The calculations were done using the numerical software environment MATLAB® (R2019b).

### Genomic prediction

SNP information was obtained from the Arabidopsis 1001 genome project information portal (SNPEFF file, version 3.1). Requiring all of the 241 accessions to have a valid (no “.” character) and homozygous allele call, and furthermore requiring the minor allele to be present in at least 10 accessions (minor allele frequency (MFA) = 4.1% of all 241 accessions) resulted in 5,613 unique SNPs. Given a genome size of approximately ~135 Mb and considering an average linkage-disequilibrium (LD) distance of 10Kb ^74,75^, SNP coverage was deemed too low (one SNP every 24Kb). Hence, we tolerated one accession to have no valid allele call yielding 25,826 unique SNPs (with MAF>4.1%, 10 accessions), corresponding to an average coverage of 5.2Kb per SNP, i.e. within the average LD) distance. In case of missing allele information, the population mean was taken as the imputed value. Alleles were encoded as −1 and 1 to reflect the two different diploid homozygous genotypes. Metabolite level data of the 37 profiled metabolites were log-transformed to render their distributions more concordant with a normal distribution.

Genomic prediction was performed applying the Best Linear Unbiased Predication (BLUP) methodology as implemented in the R-package “rrBLUP” ^27^ (Cross-validation (split of accessions into training and test population) was performed on 180 (training)/61 (test) random accession splits. As a metric of predictability, Pearson correlation coefficients of predicted vs. observed metabolite level data (log-transformed) were computed and reported over all 37 profiled metabolites.

Genome wide association analysis (GWAS) was performed to test associations between SNPs and metabolite levels for each of the metabolites, at either 16 °C or 6 °C. SNPs for all 241 accessions were obtained from the 1001 genomes project (www.1001genomes.org) and filtered to have a minor allele frequency above 5%. GWAS was done using the single trait test implemented in LIMIX ^76^. A relatedness matrix was added as covariate to the mixed effects model in order to correct for population structure.

## Acknowledgements

We would like to thank Klara Wuketich and Anneliese Auer for technical support during the phenotyping experiments. Further, we thank Matthias Nagler for assisting with sample preparation, Lena Fragner and Martin Brenner for technical assistance and Lisa Fürtauer for critical discussion. The Vienna BioCenter Core Facilities GmbH (VBCF) Plant Sciences Facility acknowledges funding from the Austrian Federal Ministry of Education, Science and Research and the City of Vienna. This work was supported by the Vienna Metabolomics Center (ViME) at the University of Vienna, and by TRR175, funded by Deutsche Forschungsgemeinschaft (DFG).

## SUPPLEMENTS

**Figure S 1.**
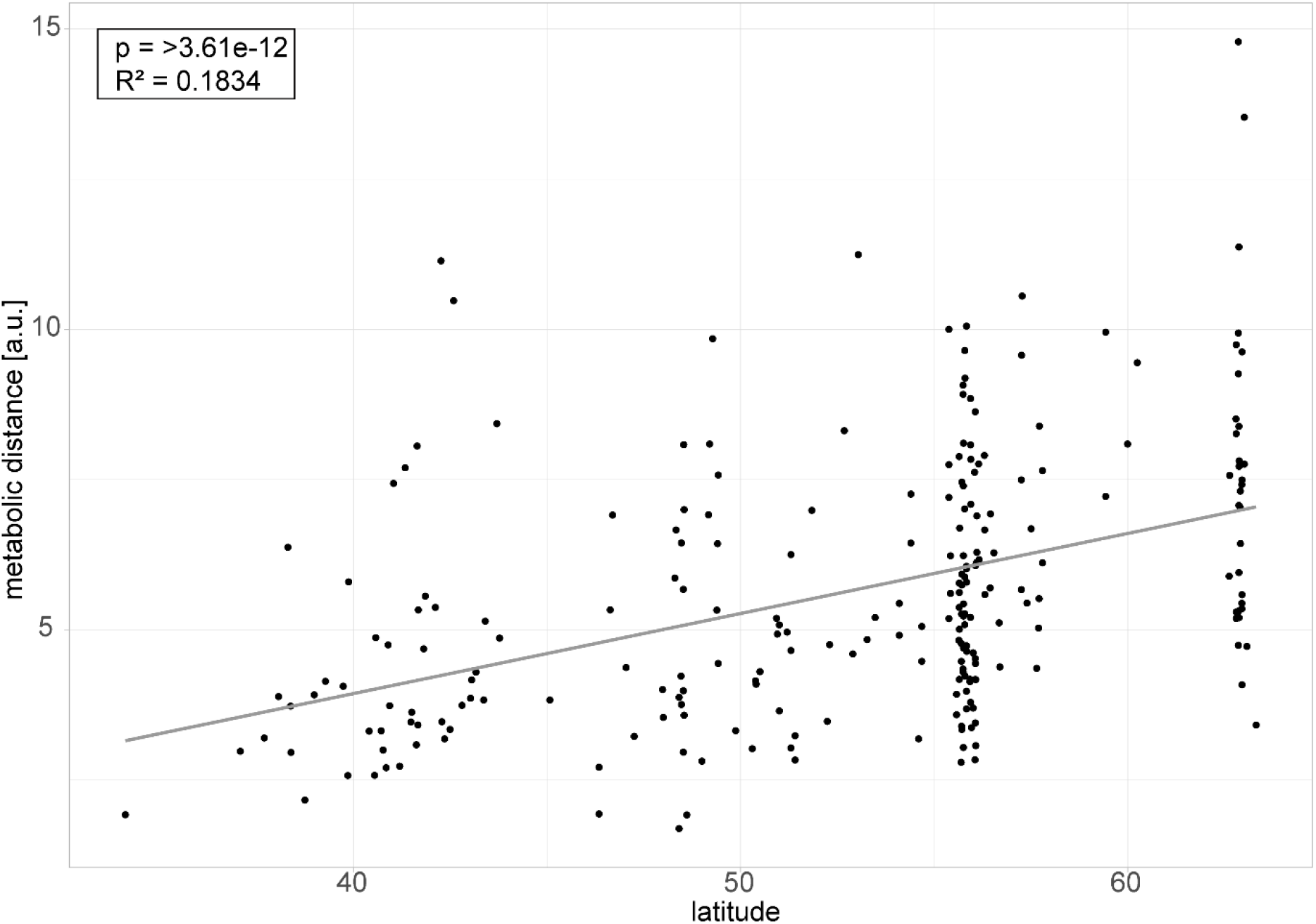
Relationship of metabolic distance and latitude of origin.

**Figure S 2.**
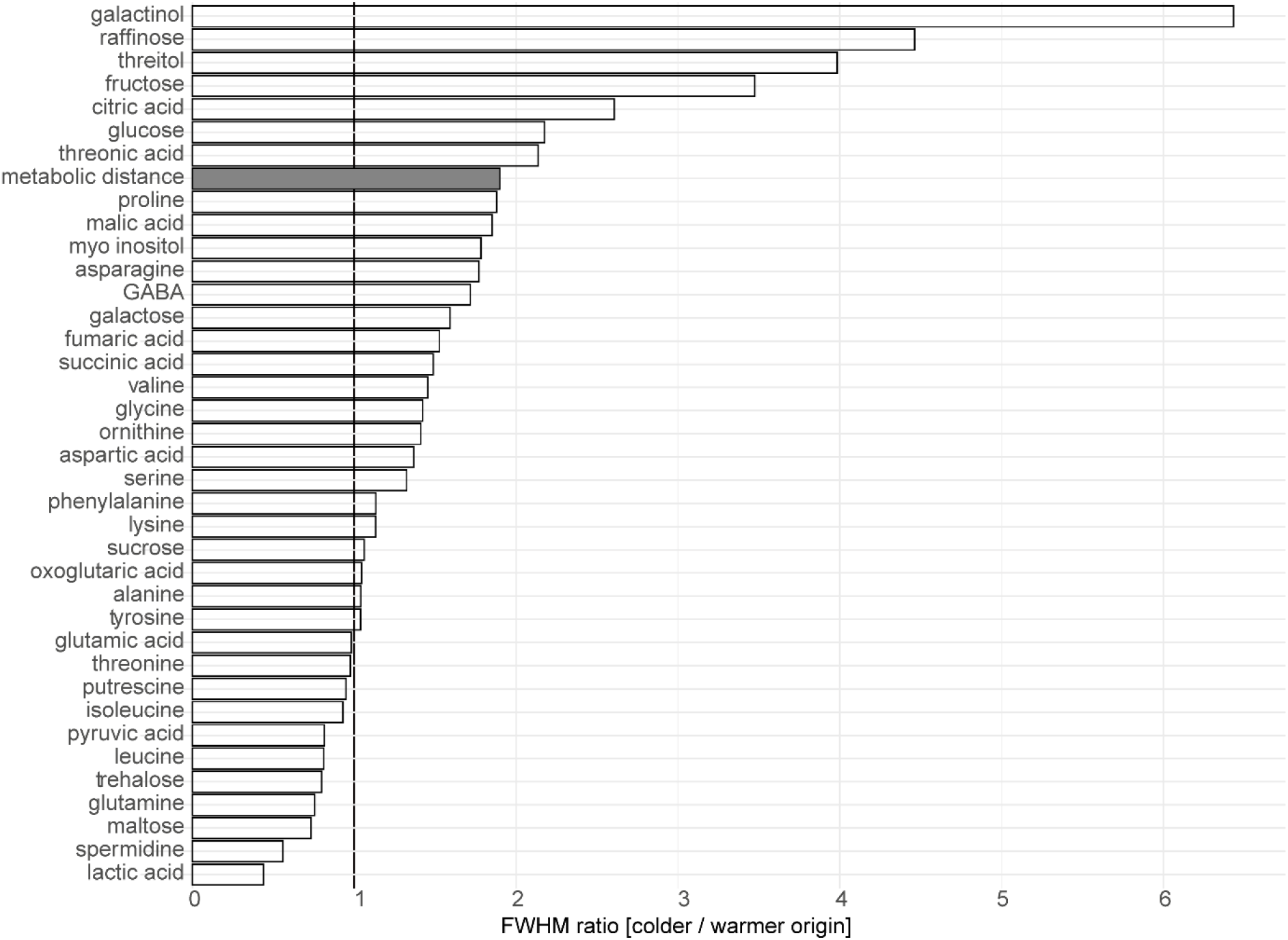
Ratios of Full Width at Half Maximum (FWHM) of kernel density functions of colder and warmer origin accessions. This ratio represents differences in variance between the groups of climatic origin. White bars – metabolites, grey bar – metabolic distance.

**Figure S 3.**
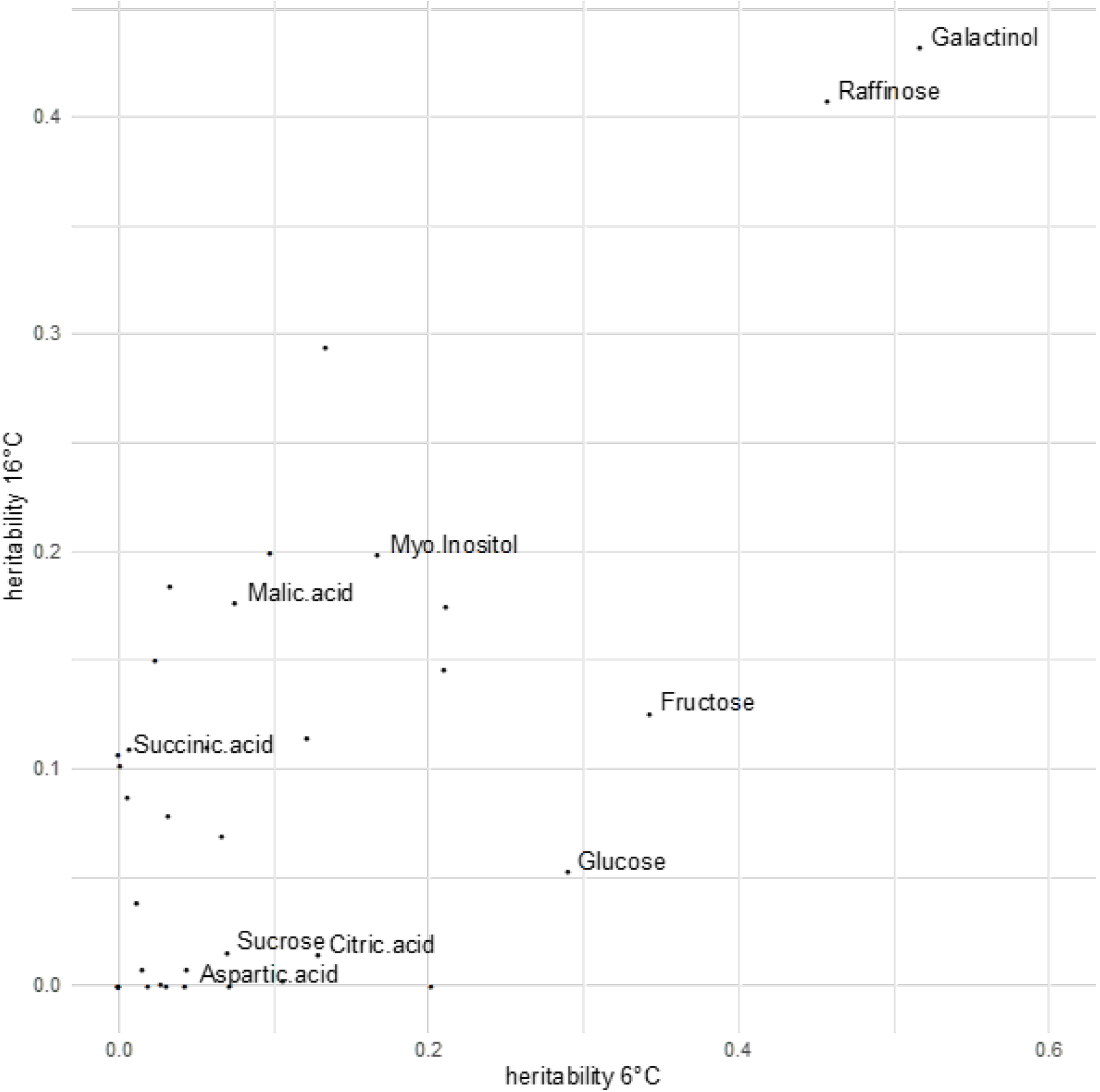
Heritabilities of individual metabolites in 16 °C and 6 °C. Broad sense heritabilities for each metabolite in 6 °C and 16 °C. Indicated are the metabolites whose increased variance were responsible for increased variance in metabolic distance.

**Figure S 4.**
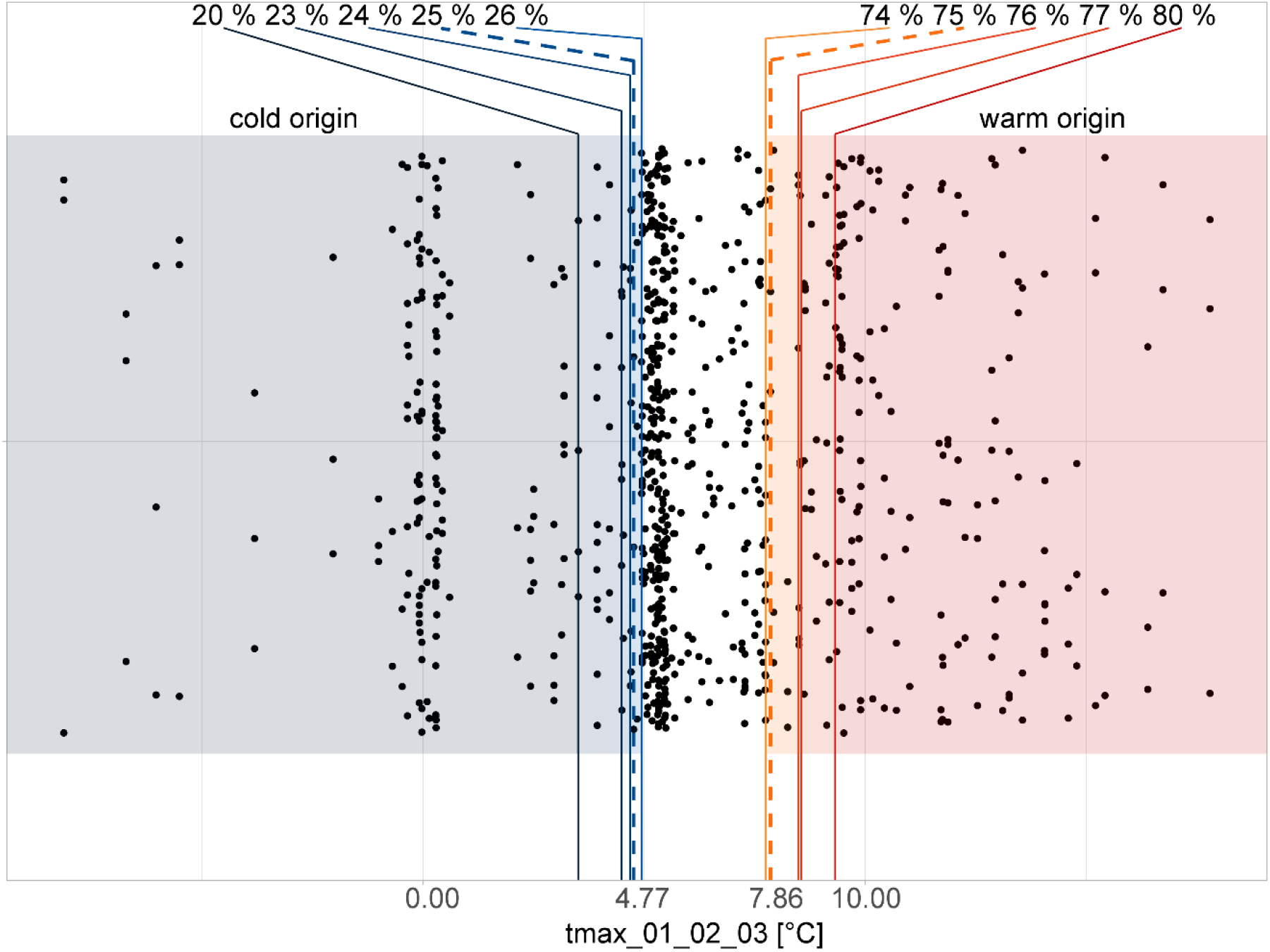
Accessions are ordered according to tmax_01_02_03 (maximum temperature in January, February and March). The lower 25 % and the upper 75 % of accessions were used as colder or warmer origin accessions respectively. For Jacobian modelling, additionally the lower 20, 23, 24 or 25 % and the upper 74, 76, 77 or 80 % of the dataset were used as variations of the threshold.

**Figure S 5.**
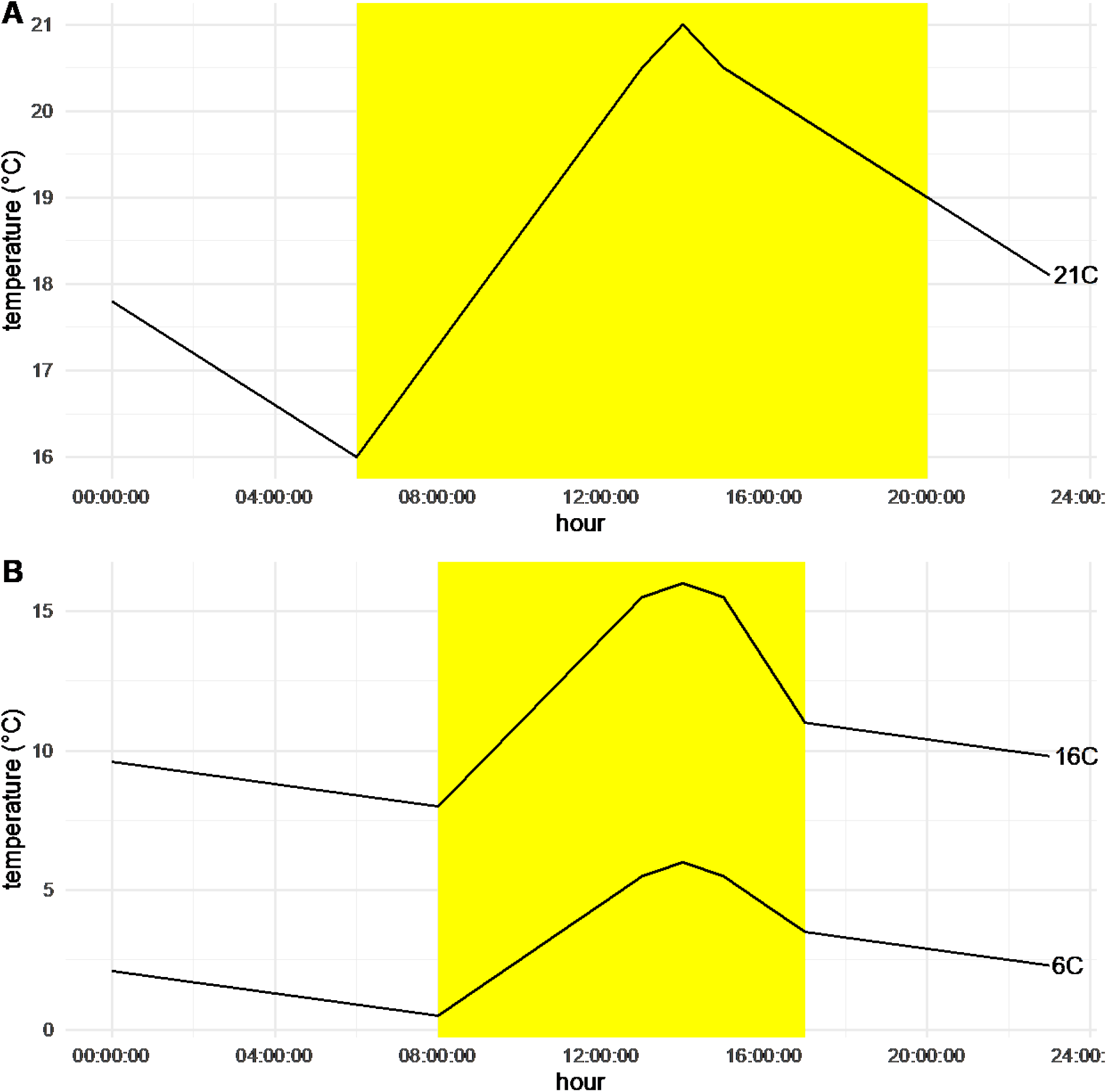
Temperature profiles of the different growth conditions. A - Temperature trajectory over 24 hours for the initial germination and growth condition with temperatures ranging from 16 to 21 °C. B – The temperature trajectory over 24 hours for the 16 °C and 6 °C growth conditions (indicated with labels 16C and 6C respectively. Yellow indicates when lights were on.

## References

1 Krasensky, J. & Jonak, C. Drought, salt, and temperature stress-induced metabolic rearrangements and regulatory networks. Journal of Experimental Botany 63, 1593–1608, doi:10.1093/jxb/err460 (2012).

2 Caldana, C. et al. High-density kinetic analysis of the metabolomic and transcriptomic response of Arabidopsis to eight environmental conditions. Plant J 67, 869–884, doi:10.1111/j.1365-313X.2011.04640.x (2011).

3 Weston, D. J. et al. Comparative physiology and transcriptional networks underlying the heat shock response in Populus trichocarpa, Arabidopsis thaliana and Glycine max. Plant, Cell & Environment 34, 1488–1506, doi:10.1111/j.1365-3040.2011.02347.x (2011).

4 Demirel, U. et al. Physiological, Biochemical, and Transcriptional Responses to Single and Combined Abiotic Stress in Stress-Tolerant and Stress-Sensitive Potato Genotypes. Frontiers in Plant Science 11, doi:10.3389/fpls.2020.00169 (2020).

5 Shankar, R., Bhattacharjee, A. & Jain, M. Transcriptome analysis in different rice cultivars provides novel insights into desiccation and salinity stress responses. Scientific Reports 6, 23719, doi:10.1038/srep23719 (2016).

6 Barah, P. et al. Genome-scale cold stress response regulatory networks in ten Arabidopsis thaliana ecotypes. BMC Genomics 14, 722, doi:10.1186/1471-2164-14-722 (2013).

7 Hoffmann, M. H. Biogeography of Arabidopsis thaliana (L.). Journal of Biogeography 29, 125–134, doi:10.1046/j.1365-2699.2002.00647.x (2002).

8 Hurry, V. Metabolic reprogramming in response to cold stress is like real estate, it’s all about location. Plant, Cell & Environment 40, 599–601, doi:10.1111/pce.12923 (2017).

9 Larcher, W. Effects of low temperature stress and frost injury on plant productivity. Physiological processes limiting plant productivity, 253–269 (1981).

10 Thakur, P., Kumar, S., Malik, J. A., Berger, J. D. & Nayyar, H. Cold stress effects on reproductive development in grain crops: An overview. Environmental and Experimental Botany 67, 429–443, doi:https://doi.org/10.1016/j.envexpbot.2009.09.004 (2010).

11 Mahajan, S. & Tuteja, N. Cold, salinity and drought stresses: an overview. Archives of biochemistry and biophysics 444, 139–158 (2005).

12 Patzke, K. et al. The Plastidic Sugar Transporter pSuT Influences Flowering and Affects Cold Responses. Plant Physiol 179, 569–587, doi:10.1104/pp.18.01036 (2019).

13 Levitt, J. Responses of Plants to Environmental Stresses. (Academic Press, INC, 1980).

14 Herrmann, H. A., Schwartz, J.-M. & Johnson, G. N. Metabolic acclimation—a key to enhancing photosynthesis in changing environments? Journal of Experimental Botany 70, 3043–3056, doi:10.1093/jxb/erz157 (2019).

15 Thomashow, M. F. Molecular Basis of Plant Cold Acclimation: Insights Gained from Studying the CBF Cold Response Pathway. Plant Physiology 154, 571–577, doi:10.1104/pp.110.161794 (2010).

16 Gehan, M. A. et al. Natural variation in the C-repeat binding factor cold response pathway correlates with local adaptation of Arabidopsis ecotypes. Plant J 84, 682–693, doi:10.1111/tpj.13027 (2015).

17 Ding, Y., Shi, Y. & Yang, S. Molecular regulation of plant responses to environmental temperatures. Mol Plant 13, 544–564, doi:10.1016/j.molp.2020.02.004 (2020).

18 Koornneef, M., Alonso-Blanco, C. & Vreugdenhil, D. Naturally Occurring Genetic Variation in Arabidopsis Thaliana. Annual Review of Plant Biology 55, 141–172, doi:10.1146/annurev.arplant.55.031903.141605 (2004).

19 Mitchell-Olds, T. & Schmitt, J. Genetic mechanisms and evolutionary significance of natural variation in Arabidopsis. Nature 441, 947–952, doi:10.1038/nature04878 (2006).

20 Nordborg, M. et al. The pattern of polymorphism in Arabidopsis thaliana. PLoS biology 3 (2005).

21 Kleessen, S. et al. Structured patterns in geographic variability of metabolic phenotypes in Arabidopsis thaliana. Nature Communications 3, 1317–1319, doi:10.1038/ncomms2333 (2012).

22 Hannah, M. A. et al. Natural genetic variation of freezing tolerance in Arabidopsis. Plant Physiol 142, 98–112, doi:10.1104/pp.106.081141 (2006).

23 Horton, M. W., Willems, G., Sasaki, E., Koornneef, M. & Nordborg, M. The genetic architecture of freezing tolerance varies across the range of Arabidopsis thaliana. Plant Cell Environ 39, 2570–2579, doi:10.1111/pce.12812 (2016).

24 Zuther, E., Schulz, E., Childs, L. H. & Hincha, D. K. Clinal variation in the non-acclimated and cold-acclimated freezing tolerance of Arabidopsis thaliana accessions. Plant, Cell and Environment 35, 1860–1878, doi:10.1111/j.1365-3040.2012.02522.x (2012).

25 Martínez-Berdeja, A. et al. Functional variants of DOG1 control seed chilling responses and variation in seasonal life-history strategies in Arabidopsis thaliana. Proceedings of the National Academy of Sciences 117, 2526–2534, doi:10.1073/pnas.1912451117 (2020).

26 Adams, W. W., Stewart, J. J., Cohu, C. M., Muller, O. & Demmig-Adams, B. Habitat Temperature and Precipitation of Arabidopsis thaliana Ecotypes Determine the Response of Foliar Vasculature, Photosynthesis, and Transpiration to Growth Temperature. Frontiers in Plant Science 7, doi:10.3389/fpls.2016.01026 (2016).

27 Endelman, J. B. Ridge Regression and Other Kernels for Genomic Selection with R Package rrBLUP. The Plant Genome 4, 250–255, doi:10.3835/plantgenome2011.08.0024 (2011).

28 The 1001 Genomes Consortium. 1,135 Genomes Reveal the Global Pattern of Polymorphism in Arabidopsis thaliana. Cell 166, 481–491, doi:10.1016/j.cell.2016.05.063 (2016).

29 Wilson, J. L. et al. Inverse Data-Driven Modeling and Multiomics Analysis Reveals Phgdh as a Metabolic Checkpoint of Macrophage Polarization and Proliferation. Cell Rep 30, 1542–1552 e1547, doi:10.1016/j.celrep.2020.01.011 (2020).

30 Nägele, T. et al. Solving the differential biochemical jacobian from metabolomics covariance data. PLoS ONE 9 (2014).

31 Ferrero-serrano, Á. & Assmann, S. M. Phenotypic and genome-wide association of natural variation with the local environment of Arabidopsis. Nature Ecology & Evolution 3, 1–41, doi:10.1038/s41559-018-0754-5 (2019).

32 Zhen, Y. & Ungerer, M. C. Clinal variation in freezing tolerance among natural accessions of Arabidopsis thaliana. New Phytologist, 1–9, doi:10.1111/j.1469-8137.2007.02262.x (2008).

33 Kang, J. et al. Natural variation of C-repeat-binding factor (CBFs) genes is a major cause of divergence in freezing tolerance among a group of Arabidopsis thaliana populations along the Yangtze River in China. New Phytologist 199, 1069–1080, doi:10.1111/nph.12335 (2013).

34 Fournier-Level, A. et al. A map of local adaptation in Arabidopsis thaliana. Science (New York, N.Y.) 334, 86–89, doi:10.1126/science.1209271 (2011).

35 Hancock, A. M. et al. Adaptation to climate across the Arabidopsis thaliana genome. Science (New York, N.Y.) 334, 83–86, doi:10.1126/science.1209244 (2011).

36 Florez-Sarasa, I. et al. Differences in Metabolic and Physiological Responses between Local and Widespread Grapevine Cultivars under Water Deficit Stress. Agronomy 10, 1052 (2020).

37 Strand, A., Hurry, V., Gustafsson, P. & Gardestrom, P. Development of Arabidopsis thaliana leaves at low temperatures releases the suppression of photosynthesis and photosynthetic gene expression despite the accumulation of soluble carbohydrates. Plant Journal 12, 605–614 (1997).

38 Meyer, R. C. et al. The metabolic signature related to high plant growth rate in <em>Arabidopsis thaliana</em>. Proceedings of the National Academy of Sciences 104, 4759–4764, doi:10.1073/pnas.0609709104 (2007).

39 Vanholme, R., Demedts, B., Morreel, K., Ralph, J. & Boerjan, W. Lignin Biosynthesis and Structure. Plant Physiology 153, 895–905, doi:10.1104/pp.110.155119 (2010).

40 Schulz, E., Tohge, T., Zuther, E., Fernie, A. R. & Hincha, D. K. Flavonoids are determinants of freezing tolerance and cold acclimation in Arabidopsis thaliana. Scientific Reports 6, 34027, doi:ARTN 34027 10.1038/srep34027 (2016).

41 Dyson, B. C. et al. FUM2, a Cytosolic Fumarase, Is Essential for Acclimation to Low Temperature in <em>Arabidopsis thaliana</em>. Plant Physiology 172, 118–127, doi:10.1104/pp.16.00852 (2016).

42 Weiszmann, J., Fürtauer, L., Weckwerth, W. & Nägele, T. Vacuolar invertase activity shapes photosynthetic stress response of Arabidopsis thaliana and stabilizes central energy supply. bioRxiv (2017).

43 Riewe, D. et al. A naturally occurring promoter polymorphism of the Arabidopsis FUM2 gene causes expression variation, and is associated with metabolic and growth traits. Plant J 88, 826–838, doi:10.1111/tpj.13303 (2016).

44 Schulze, W. X., Schneider, T., Starck, S., Martinoia, E. & Trentmann, O. Cold acclimation induces changes in Arabidopsis tonoplast protein abundance and activity and alters phosphorylation of tonoplast monosaccharide transporters. The Plant Journal 69, 529–541, doi:10.1111/j.1365-313X.2011.04812.x (2012).

45 Planchais, S. et al. BASIC AMINO ACID CARRIER 2 gene expression modulates arginine and urea content and stress recovery in Arabidopsis leaves. Front Plant Sci 5, 330, doi:10.3389/fpls.2014.00330 (2014).

46 Witte, C. P. Urea metabolism in plants. Plant Sci 180, 431–438, doi:10.1016/j.plantsci.2010.11.010 (2011).

47 Guy, C., Kaplan, F., Kopka, J., Selbig, J. & Hincha, D. K. Metabolomics of temperature stress. Physiologia Plantarum 132, 220–235, doi:10.1111/j.1399-3054.2007.00999.x (2008).

48 Espinoza, C. et al. Interaction with Diurnal and Circadian Regulation Results in Dynamic Metabolic and Transcriptional Changes during Cold Acclimation in Arabidopsis. PLoS ONE 5, e14101–e14101, doi:10.1371/journal.pone.0014101 (2010).

49 Less, H. & Galili, G. Principal transcriptional programs regulating plant amino acid metabolism in response to abiotic stresses. Plant Physiol 147, 316–330, doi:10.1104/pp.108.115733 (2008).

50 Weiszmann, J., Fürtauer, L., Weckwerth, W. & Näxgele, T. Vacuolar sucrose cleavage prevents limitation of cytosolic carbohydrate metabolism and stabilizes photosynthesis under abiotic stress. FEBS J 285, 4082–4098, doi:10.1111/febs.14656 (2018).

51 Geigenberger, P. & Stitt, M. A futile cycle of sucrose synthesis and degradation is involved regulating partitioning between sucrose starch and respiration in cotyledons of germinating ricinus-communis l. Seedlings when phloem transport is inhibited. Planta 185, 81–90 (1991).

52 Ruan, Y.-L. Sucrose metabolism: gateway to diverse carbon use and sugar signaling. Annual review of plant biology, doi:10.1146/annurev-arplant-050213-040251 (2014).

53 Ben-Izhak Monselise, E., Parola, A. H. & Kost, D. Low-frequency electromagnetic fields induce a stress effect upon higher plants, as evident by the universal stress signal, alanine. Biochemical and Biophysical Research Communications 302, 427–434, doi:10.1016/s0006-291x(03)00194-3 (2003).

54 Zhang, Q. et al. Characterization of Arabidopsis serine:glyoxylate aminotransferase, AGT1, as an asparagine aminotransferase. Phytochemistry 85, 30–35, doi:10.1016/j.phytochem.2012.09.017 (2013).

55 Betsche, T. Aminotransfer from Alanine and Glutamate to Glycine and Serine during Photorespiration in Oat Leaves. Plant Physiology 71, 961–965, doi:10.1104/pp.71.4.961 (1983).

56 Obata, T. & Fernie, A. R. The use of metabolomics to dissect plant responses to abiotic stresses. Cellular and Molecular Life Sciences 69, 3225–3243, doi:10.1007/s00018-012-1091-5 (2012).

57 Maruyama, K. et al. Integrated Analysis of the Effects of Cold and Dehydration on Rice Metabolites, Phytohormones, and Gene Transcripts. Plant Physiology 164, 1759–1771, doi:10.1104/pp.113.231720 (2014).

58 Junker, A. et al. Optimizing experimental procedures for quantitative evaluation of crop plant performance in high throughput phenotyping systems. Frontiers in Plant Science 5, doi:10.3389/fpls.2014.00770 (2015).

59 Paine, C. E. T. et al. How to fit nonlinear plant growth models and calculate growth rates: an update for ecologists. Methods in Ecology and Evolution 3, 245–256, doi:10.1111/j.2041-210X.2011.00155.x (2012).

60 nlme: Linear and Nonlinear Mixed Effects Models (2020).

61 Bürkner, P.-C. brms: An R Package for Bayesian Multilevel Models Using Stan. 2017 80, 28, doi:10.18637/jss.v080.i01 (2017).

62 emmeans: Estimated Marginal Means, aka Least-Squares Means (2020).

63 R: A Language and Environment for Statistical Computing (R Foundation for Statistical Computing, 2019).

64 Wickham, H. et al. Welcome to the Tidyverse. Journal of Open Source Software 4, 1686 (2019).

65 Yuan, T., Masaaki, H. & Wenxuan, L. ggfortify: Unified Interface to Visualize Statistical Result of Popular R Packages. The R Journal 8 (2016).

66 Houshyani, B. et al. Characterization of the natural variation in Arabidopsis thaliana metabolome by the analysis of metabolic distance. Metabolomics 8, 131–145, doi:10.1007/s11306-011-0375-3 (2012).

67 Hmisc: Harrell Miscellaneous (2020).

68 Fick, S. E. & Hijmans, R. J. WorldClim 2: new 1-km spatial resolution climate surfaces for global land areas. International Journal of Climatology 37, 4302–4315, doi:10.1002/joc.5086 (2017).

69 caret: Classification and Regression Training (2019).

70 leaps: Regression Subset Selection (2020).

71 Friedman, J., Hastie, T. & Tibshirani, R. Regularization Paths for Generalized Linear Models via Coordinate Descent. J Stat Softw 33, 1–22, doi:10.18637/jss.v033.i01 (2010).

72 Oksanen, J. et al. Package ‘vegan’. Community ecology package, version 2, 1–295 (2013).

73 Nägele, T. Linking metabolomics data to underlying metabolic regulation. Frontiers in Molecular Biosciences 1, 1–6, doi:10.3389/fmolb.2014.00022 (2014).

74 Kim, S. et al. Recombination and linkage disequilibrium in Arabidopsis thaliana. Nat Genet 39, 1151–1155, doi:10.1038/ng2115 (2007).

75 Perlaza-Jiménez, L. & Walther, D. A genome-wide scan for correlated mutations detects macromolecular and chromatin interactions in Arabidopsis thaliana. Nucleic Acids Research 46, 8114–8132, doi:10.1093/nar/gky576 (2018).

76 Lippert, C., Casale, F. P., Rakitsch, B. & Stegle, O. LIMIX: genetic analysis of multiple traits. bioRxiv, 003905, doi:10.1101/003905 (2014).

