## Supplementary material for "Plasticity of the primary metabolome in 241 cold grown *Arabidopsis thaliana* accessions and its relation to natural habitat temperature": S_Table1

Table S 1 Spearman’s rank correlation coefficients of metabolic distance and all climate variables. Additionally, results of partial Mantel tests, correcting for population structure via a genetic kinship matrix are given for climate summary variables

|  | Spearman correlation | | Mantel Test (kinship correction) | |
| --- | --- | --- | --- | --- |
| **variable** | **Spearman’s ρ** | **p-value** | **r** | **p-value** |
| maximum temperature Feb | -0.558 | <1E-16 |  |  |
| maximum temperature Jan, Feb, Mar (Q1) | -0.551 | <1E-16 | 0.188 | 0.001 |
| maximum temperature Jan | -0.549 | <1E-16 |  |  |
| average temperature Jan, Feb, Mar (Q1) | -0.536 | <1E-16 | 0.196 | 0.001 |
| mean temperature of coldest quarter [BIO11] | -0.531 | <1E-16 |  |  |
| average temperature Mar | -0.531 | <1E-16 |  |  |
| maximum temperature Mar | -0.530 | <1E-16 |  |  |
| average temperature Feb | -0.529 | <1E-16 |  |  |
| average temperature Jan | -0.523 | <1E-16 |  |  |
| annual mean temperature [BIO01] | -0.518 | <1E-16 |  |  |
| maximum temperature Dec | -0.518 | <1E-16 |  |  |
| maximum temperature Oct, Nov, Dec (Q4) | -0.510 | <1E-16 | 0.127 | 0.001 |
| average temperature Dec | -0.508 | <1E-16 |  |  |
| maximum temperature Oct | -0.506 | <1E-16 |  |  |
| minimum temperature Feb | -0.505 | <1E-16 |  |  |
| minimum temperature Mar | -0.505 | <1E-16 |  |  |
| average temperature Oct, Nov, Dev (Q4) | -0.504 | <1E-16 | 0.154 | 0.002 |
| maximum temperature Nov | -0.502 | <1E-16 |  |  |
| minimum temperature Jan, Feb, Mar (Q1) | -0.501 | <1E-16 | 0.183 | 0.001 |
| average temperature Oct | -0.496 | 2.22E-16 |  |  |
| average temperature Sept | -0.495 | 2.22E-16 |  |  |
| average temperature Apr | -0.492 | 4.44E-16 |  |  |
| vapour pressure Mar | -0.487 | 8.88E-16 |  |  |
| maximum temperature Sept | -0.485 | 1.33E-15 |  |  |
| isothermality (bio_2/bio_7) (×100) [BIO03] | -0.484 | 1.33E-15 |  |  |
| average temperature Nov | -0.483 | 1.78E-15 |  |  |
| vapour pressure Jan, Feb, Mar (Q1) | -0.477 | 4.44E-15 | 0.136 | 0.002 |
| solar radiation Nov | -0.474 | 6.66E-15 |  |  |
| minimum temperature Apr | -0.472 | 9.33E-15 |  |  |
| solar radiation Oct, Nov, Dec (Q4) | -0.470 | 1.15E-14 | 0.064 | 0.039 |
| minimum temperature Sept | -0.470 | 1.15E-14 |  |  |
| solar radiation Dec | -0.468 | 1.55E-14 |  |  |
| maximum temperature Apr | -0.467 | 1.82E-14 |  |  |
| vapour pressure Feb | -0.466 | 2.13E-14 |  |  |
| vapour pressure Apr | -0.463 | 3.24E-14 |  |  |
| solar radiation Oct | -0.461 | 4.49E-14 |  |  |
| average temperature Apr, May, Jun (Q2) | -0.460 | 5.00E-14 | 0.101 | 0.006 |
| solar radiation Jan | -0.459 | 5.68E-14 |  |  |
| average temperature Jul, Aug, Sept (Q3) | -0.458 | 6.79E-14 | 0.050 | 0.091 |
| min temperature of coldest month [BIO06] | -0.455 | 9.81E-14 |  |  |
| solar radiation Feb | -0.454 | 1.22E-13 |  |  |
| maximum temperature Jul, Aug, Sept (Q3) | -0.452 | 1.45E-13 | 0.032 | 0.197 |
| minimum temperature Oct | -0.452 | 1.60E-13 |  |  |
| minimum temperature Oct, Nov, Dec (Q4) | -0.447 | 2.95E-13 | 0.157 | 0.003 |
| vapour pressure Jan | -0.444 | 4.44E-13 |  |  |
| minimum temperature Apr, May, Jun (Q2) | -0.444 | 4.49E-13 | 0.125 | 0.003 |
| vapour pressure Oct | -0.443 | 5.01E-13 |  |  |
| minimum temperature Jan | -0.440 | 7.36E-13 |  |  |
| average temperature May | -0.439 | 9.09E-13 |  |  |
| vapour pressure Dec | -0.438 | 9.78E-13 |  |  |
| vapour pressure Oct, Nov, Dec (Q4) | -0.436 | 1.41E-12 | 0.132 | 0.001 |
| average temperature Aug | -0.434 | 1.62E-12 |  |  |
| maximum temperature Aug | -0.433 | 2.06E-12 |  |  |
| solar radiation Jan, Feb, Mar (Q1) | -0.432 | 2.10E-12 | 0.044 | 0.061 |
| minimum temperature Nov | -0.432 | 2.22E-12 |  |  |
| minimum temperature Dec | -0.430 | 2.75E-12 |  |  |
| mean temperature of warmest quarter [BIO10] | -0.425 | 5.18E-12 |  |  |
| minimum temperature May | -0.422 | 7.85E-12 |  |  |
| vapour pressure May | -0.419 | 1.09E-11 |  |  |
| maximum temperature Apr, May, Jun (Q2) | -0.413 | 2.30E-11 | 0.071 | 0.026 |
| maximum temperature Jul | -0.410 | 3.67E-11 |  |  |
| average temperature Jul | -0.409 | 3.75E-11 |  |  |
| max temperature of warmest month [BIO05] | -0.403 | 8.17E-11 |  |  |
| solar radiation Sept | -0.401 | 9.97E-11 |  |  |
| vapour pressure Sept | -0.401 | 1.02E-10 |  |  |
| maximum temperature May | -0.400 | 1.17E-10 |  |  |
| vapour pressure Apr, May, Jun (Q2) | -0.399 | 1.20E-10 | 0.100 | 0.012 |
| minimum temperature Jul, Aug, Sept (Q3) | -0.399 | 1.32E-10 | 0.082 | 0.031 |
| vapour pressure Nov | -0.393 | 2.54E-10 |  |  |
| solar radiation Mar | -0.392 | 2.88E-10 |  |  |
| average temperature Jun | -0.389 | 3.75E-10 |  |  |
| minimum temperature Jun | -0.380 | 1.12E-09 |  |  |
| mean temperature of driest quarter [BIO09] | -0.378 | 1.30E-09 |  |  |
| minimum temperature Aug | -0.367 | 4.23E-09 |  |  |
| maximum temperature Jun | -0.348 | 2.80E-08 |  |  |
| solar radiation Aug | -0.337 | 7.98E-08 |  |  |
| precipitation May | -0.318 | 4.68E-07 |  |  |
| minimum temperature Jul | -0.318 | 4.69E-07 |  |  |
| solar radiation Jul, Aug, Sept (Q3) | -0.318 | 4.72E-07 | 0.011 | 0.357 |
| temperature seasonality (standard deviation ×100) [BIO04] | 0.306 | 1.33E-06 |  |  |
| solar radiation Apr | -0.294 | 3.41E-06 |  |  |
| vapour pressure Jun | -0.289 | 4.90E-06 |  |  |
| mean diurnal range (mean of monthly (max temp - min temp)) [BIO02] | -0.287 | 5.75E-06 |  |  |
| wind Sept | 0.275 | 1.48E-05 |  |  |
| precipitation Apr | -0.270 | 2.13E-05 |  |  |
| precipitation Jul, Aug, Sept (Q3) | 0.266 | 2.95E-05 | -0.036 | 0.828 |
| vapour pressure Jul, Aug, Sept (Q3) | -0.265 | 3.07E-05 | 0.073 | 0.047 |
| wind Oct | 0.265 | 3.09E-05 |  |  |
| wind Jun | 0.254 | 6.62E-05 |  |  |
| precipitation Feb | -0.253 | 7.12E-05 |  |  |
| wind Nov | 0.252 | 7.82E-05 |  |  |
| precipitation Jul | 0.251 | 8.04E-05 |  |  |
| wind Jul, Aug, Sept (Q3) | 0.250 | 8.73E-05 | 0.000 | 0.496 |
| wind Oct, Nov, Dec (Q4) | 0.245 | 1.24E-04 | -0.009 | 0.610 |
| precipitation Aug | 0.244 | 1.26E-04 |  |  |
| wind Aug | 0.238 | 1.92E-04 |  |  |
| wind Dec | 0.220 | 5.79E-04 |  |  |
| wind Jul | 0.218 | 6.46E-04 |  |  |
| wind Jan | 0.214 | 8.06E-04 |  |  |
| wind May | 0.210 | 1.03E-03 |  |  |
| wind Apr, May, Jun (Q2) | 0.210 | 1.05E-03 | -0.013 | 0.641 |
| vapour pressure Aug | -0.205 | 1.36E-03 |  |  |
| precipitation of coldest quarter [BIO19] | -0.201 | 1.69E-03 |  |  |
| precipitation Jan, Feb, Mar (Q1) | -0.197 | 2.15E-03 | -0.039 | 0.817 |
| precipitation of warmest quarter [BIO18] | 0.196 | 2.27E-03 |  |  |
| wind Jan, Feb, Mar (Q1) | 0.192 | 2.74E-03 | -0.026 | 0.794 |
| precipitation Mar | -0.191 | 2.90E-03 |  |  |
| precipitation Sept | 0.183 | 4.34E-03 |  |  |
| mean temperature of wettest quarter [BIO08] | -0.182 | 4.49E-03 |  |  |
| wind Feb | 0.180 | 4.99E-03 |  |  |
| solar radiation Jul | -0.178 | 5.58E-03 |  |  |
| precipitation Dec | -0.175 | 6.43E-03 |  |  |
| wind Mar | 0.167 | 9.22E-03 |  |  |
| precipitation Apr, May, Jun (Q2) | -0.164 | 0.011 | -0.066 | 0.990 |
| wind Apr | 0.162 | 0.011 |  |  |
| solar radiation Apr, May, Jun (Q2) | -0.156 | 0.015 | -0.031 | 0.822 |
| precipitation Jan | -0.131 | 0.042 |  |  |
| vapour pressure Jul | -0.130 | 0.043 |  |  |
| solar radiation May | -0.126 | 0.052 |  |  |
| annual precipitation [BIO12] | -0.093 | 0.150 |  |  |
| temperature annual range (BIO05-BIO06) [BIO07] | 0.080 | 0.214 |  |  |
| precipitation Oct, Nov, Dec (Q4) | -0.051 | 0.433 | -0.052 | 0.928 |
| precipitation seasonality (coefficient of variation) [BIO15] | 0.037 | 0.570 |  |  |
| precipitation Jun | 0.033 | 0.611 |  |  |
| precipitation of wettest month [BIO13] | -0.031 | 0.636 |  |  |
| precipitation of wettest quarter [BIO16] | -0.028 | 0.670 |  |  |
| solar radiation Jun | 0.022 | 0.740 |  |  |
| precipitation Oct | -0.017 | 0.791 |  |  |
| precipitation of driest quarter [BIO17] | -0.007 | 0.919 |  |  |
| precipitation Nov | 0.006 | 0.927 |  |  |
| precipitation of driest month [BIO14] | 0.003 | 0.967 |  |  |
