## Supplementary material for "Plasticity of the primary metabolome in 241 cold grown *Arabidopsis thaliana* accessions and its relation to natural habitat temperature": S_Table5

mGWAS of metabolite concentrations in the 16 °C condition, red line indicates significance threshold after Bonferroni correction:

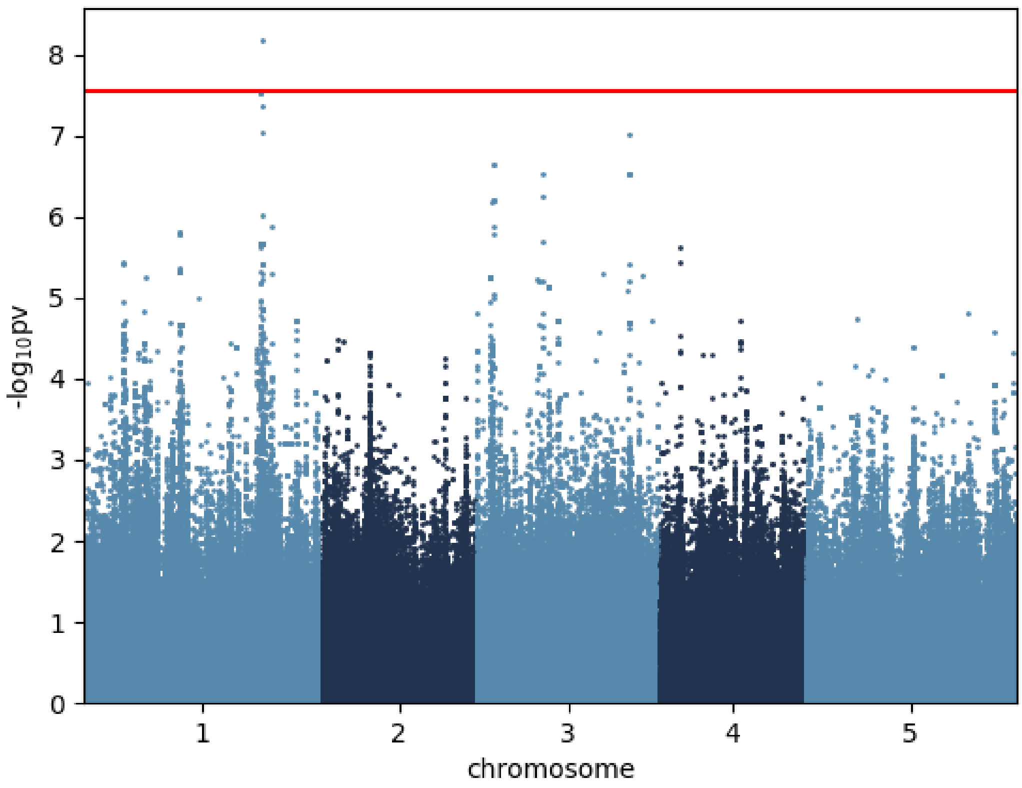


Alanine_16 °C

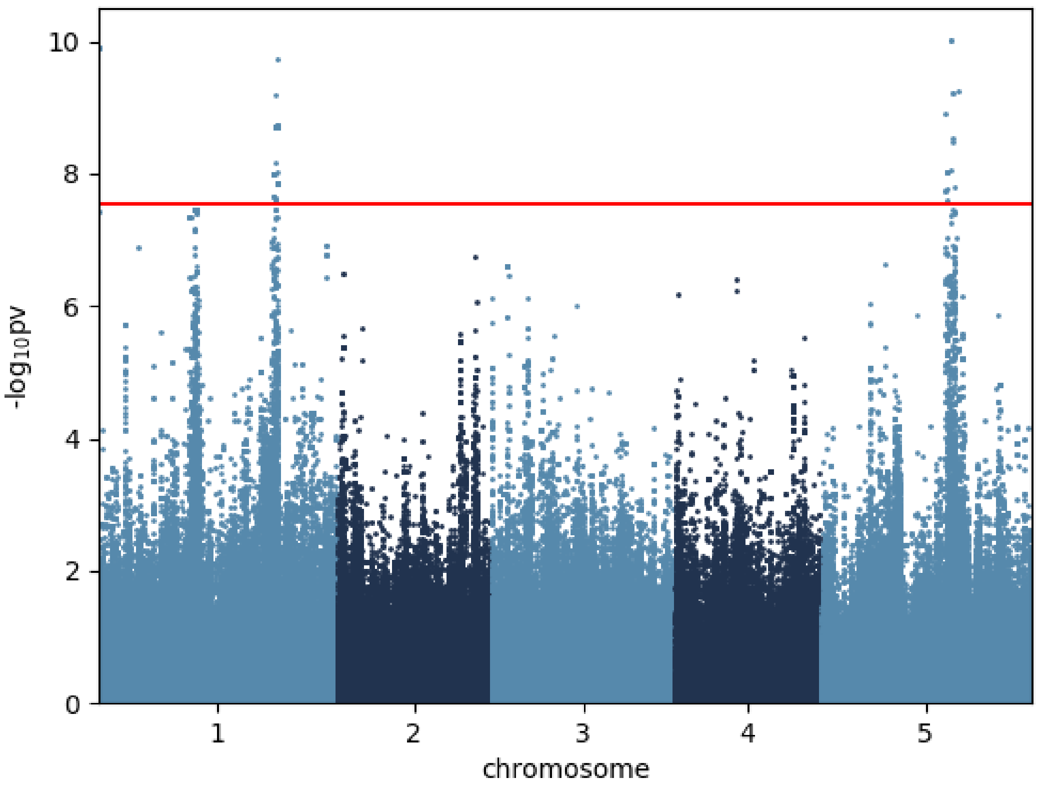


Asparagine_16 °C

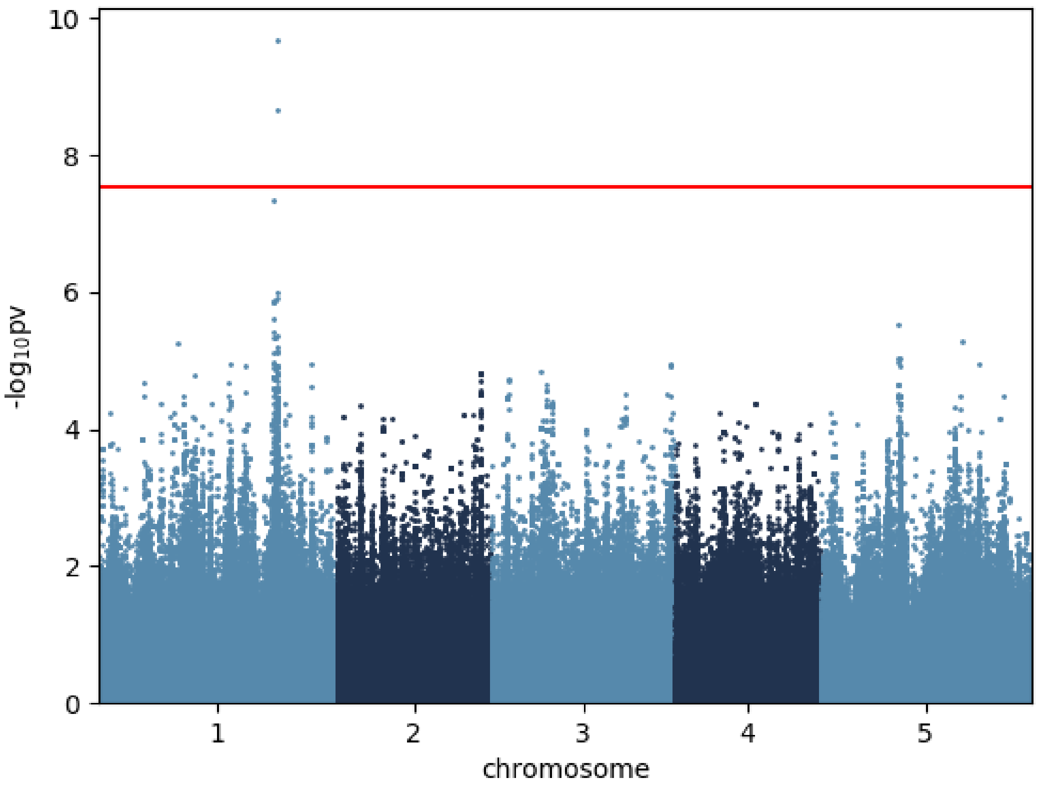


Aspartic.acid_16 °C

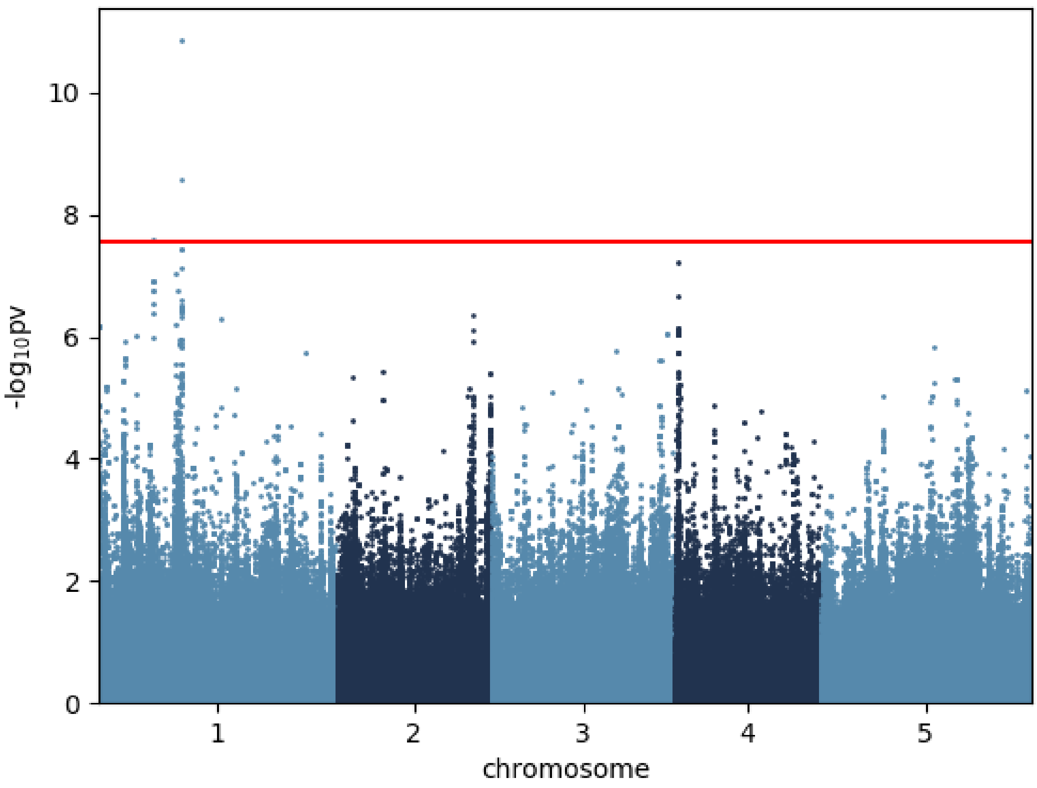


Butanoic.acid.4.amino._16 °C

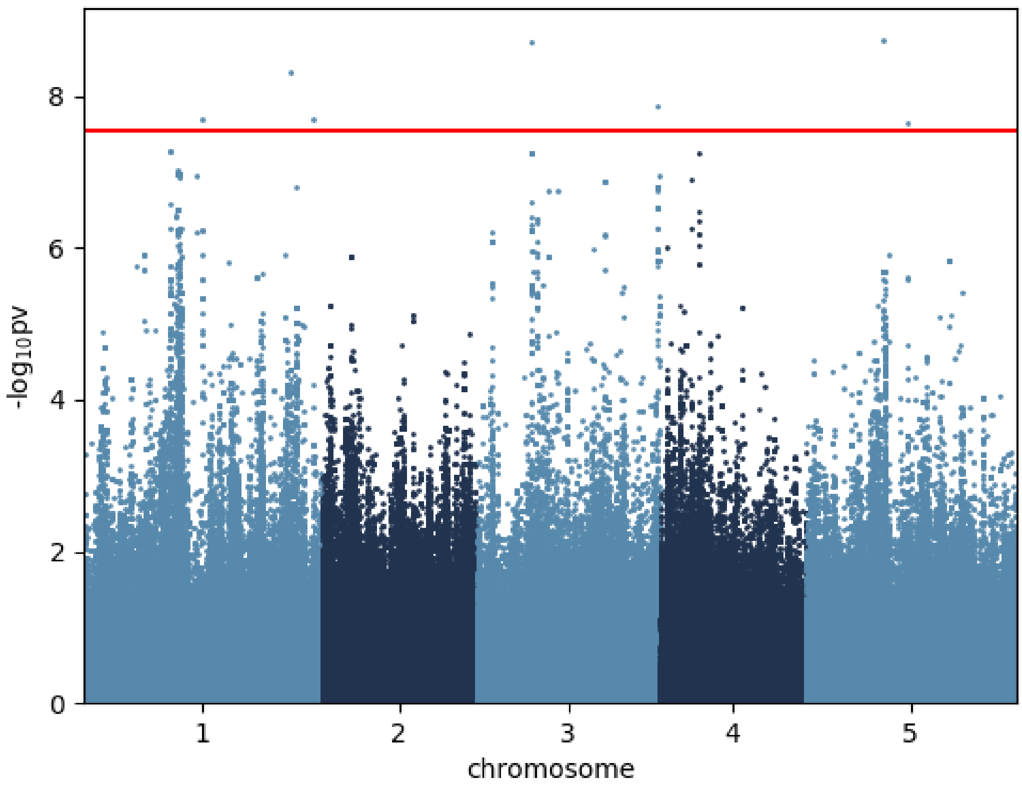


Citric.acid_16 °C

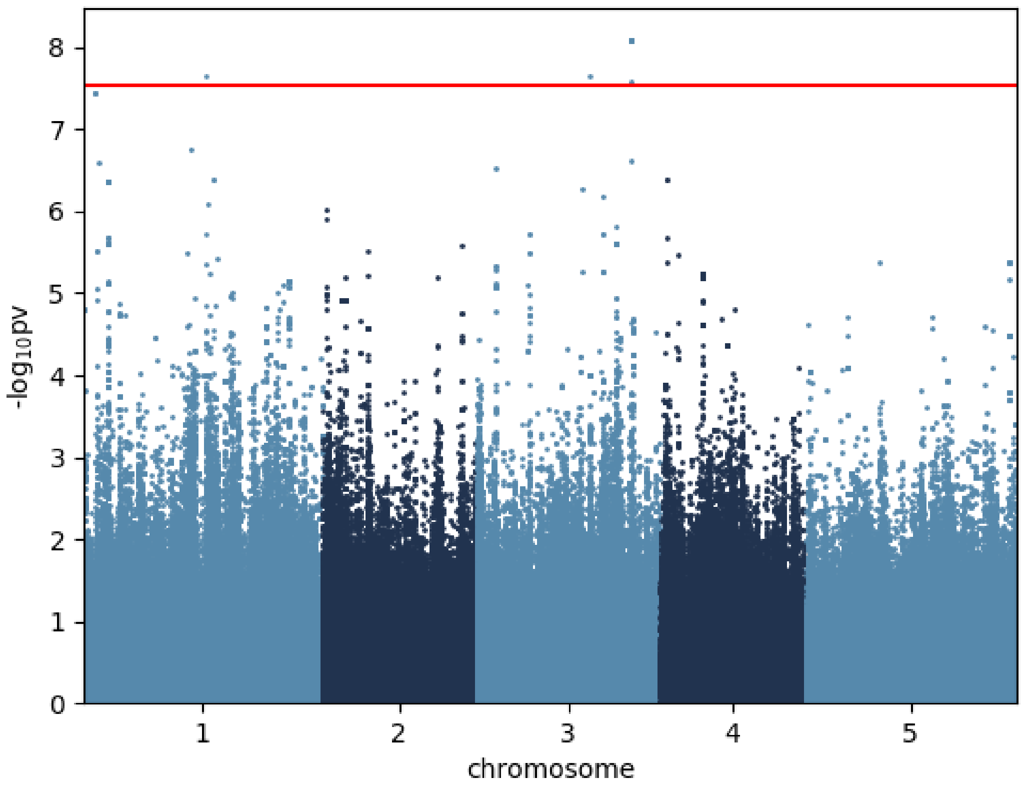


Fructose_16 °C

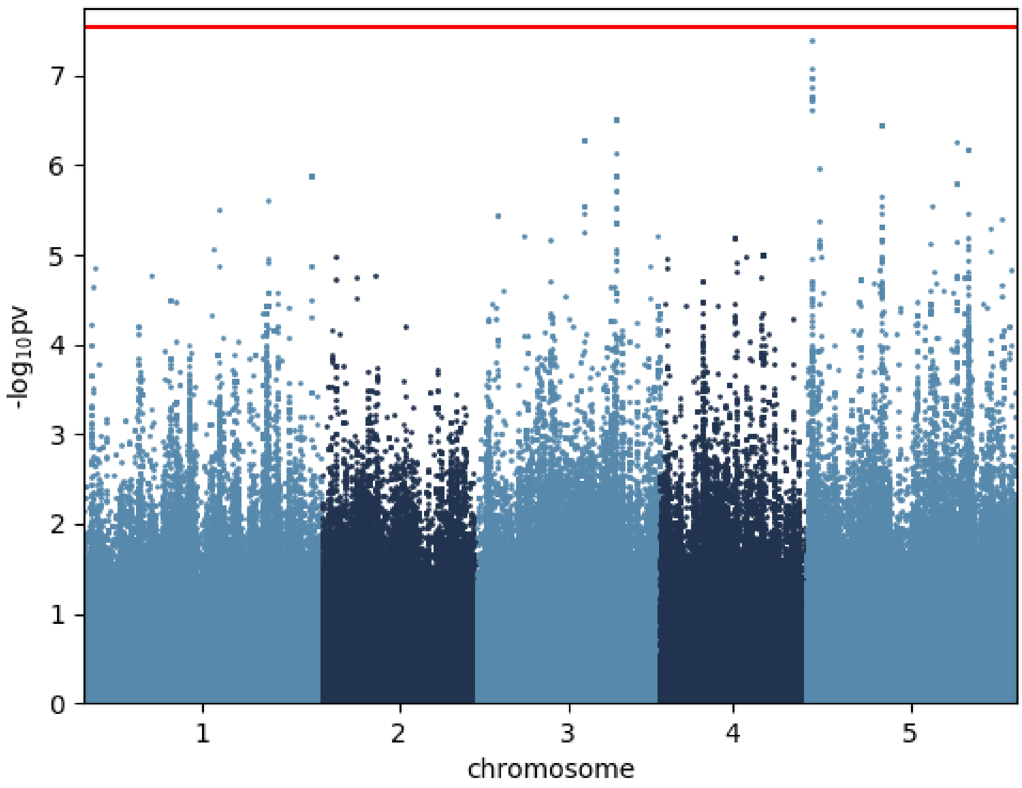


Fumaric.acid_16 °C

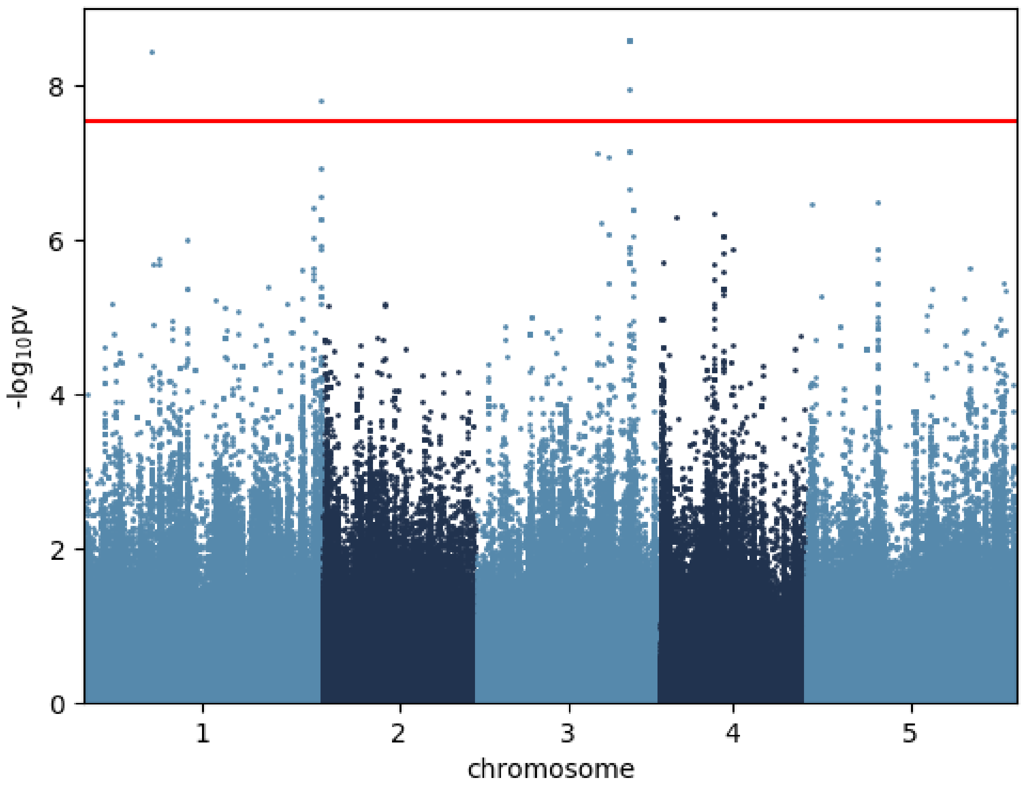


Galactinol_16 °C

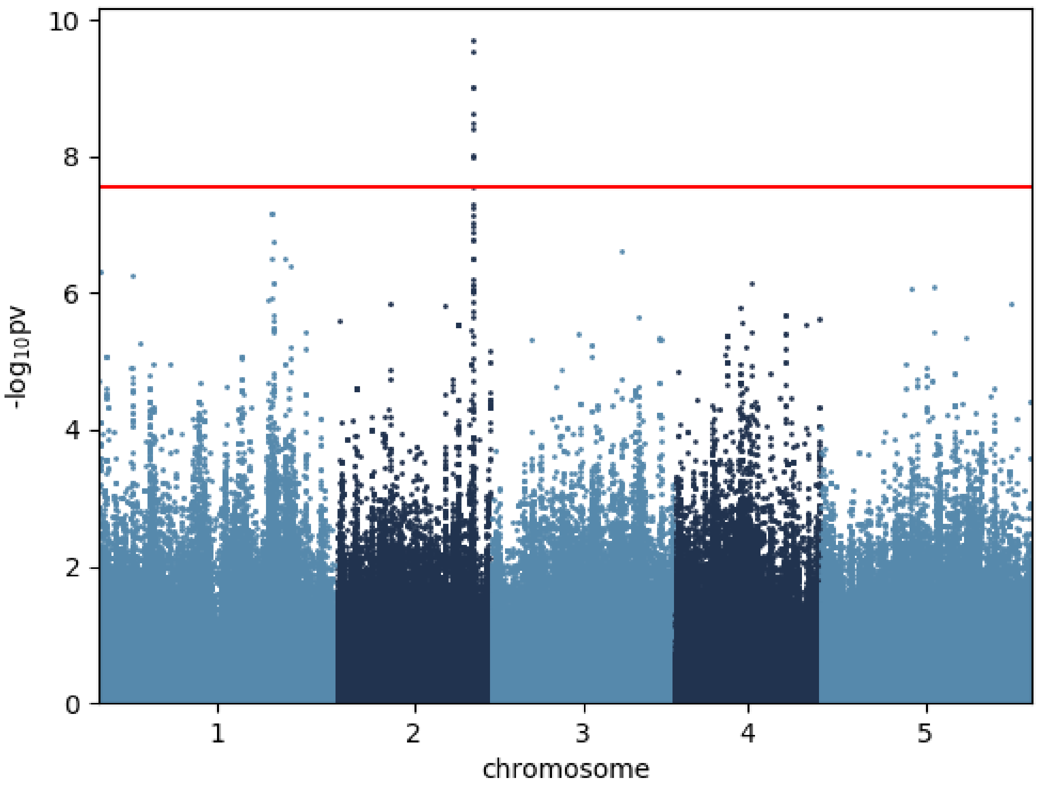


Galactose_16 °C

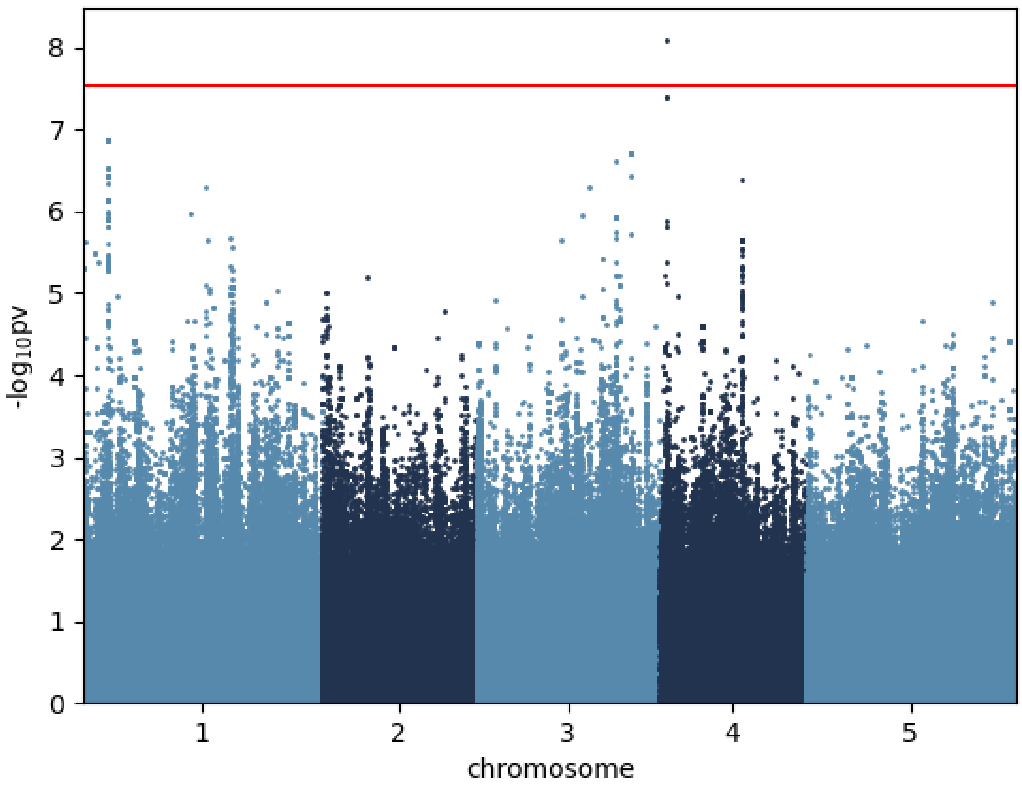


Glucose_16 °C

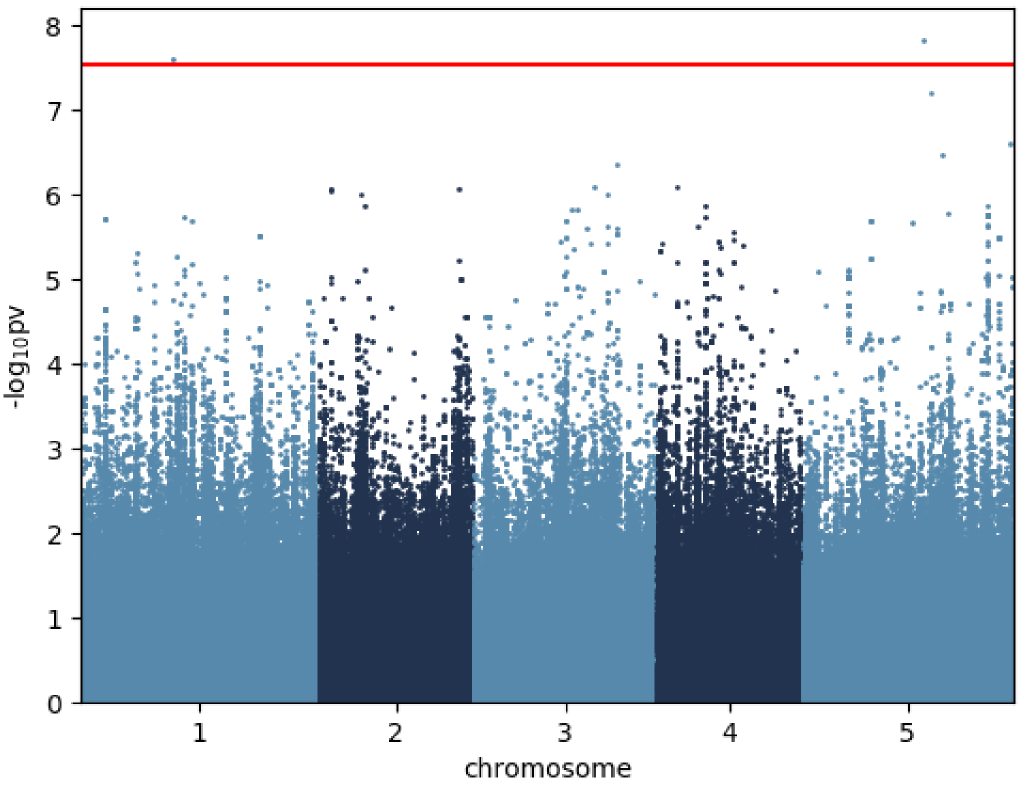


Glutamic.acid_16 °C

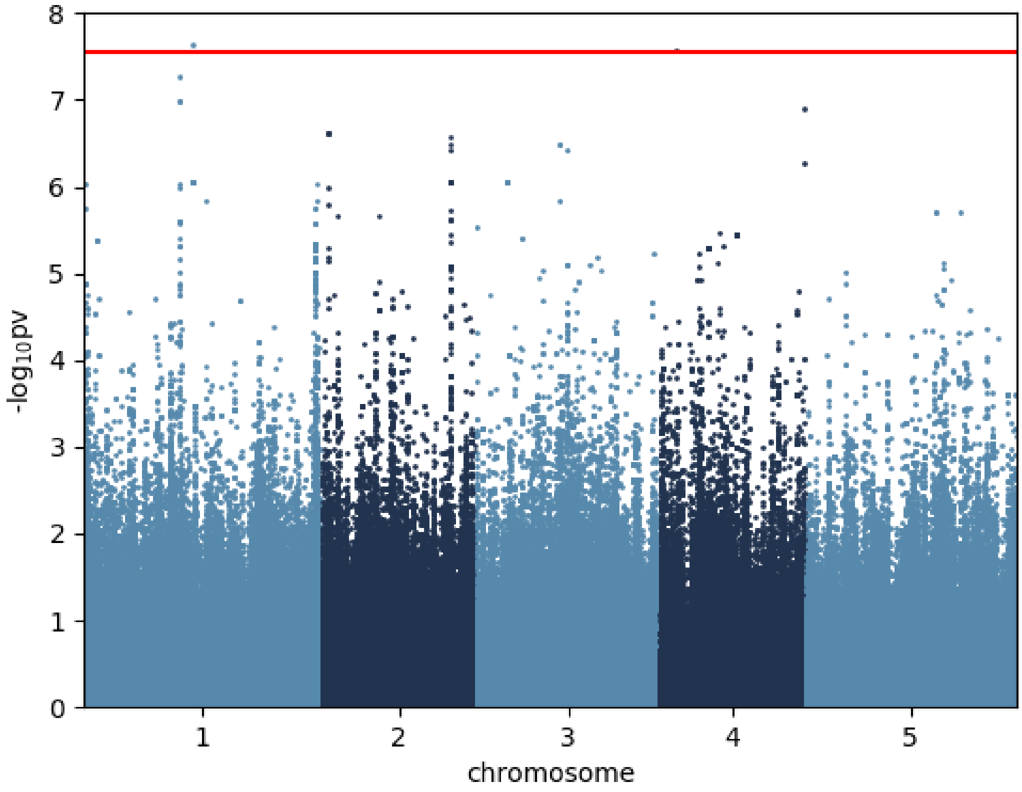


Glutamine_16 °C

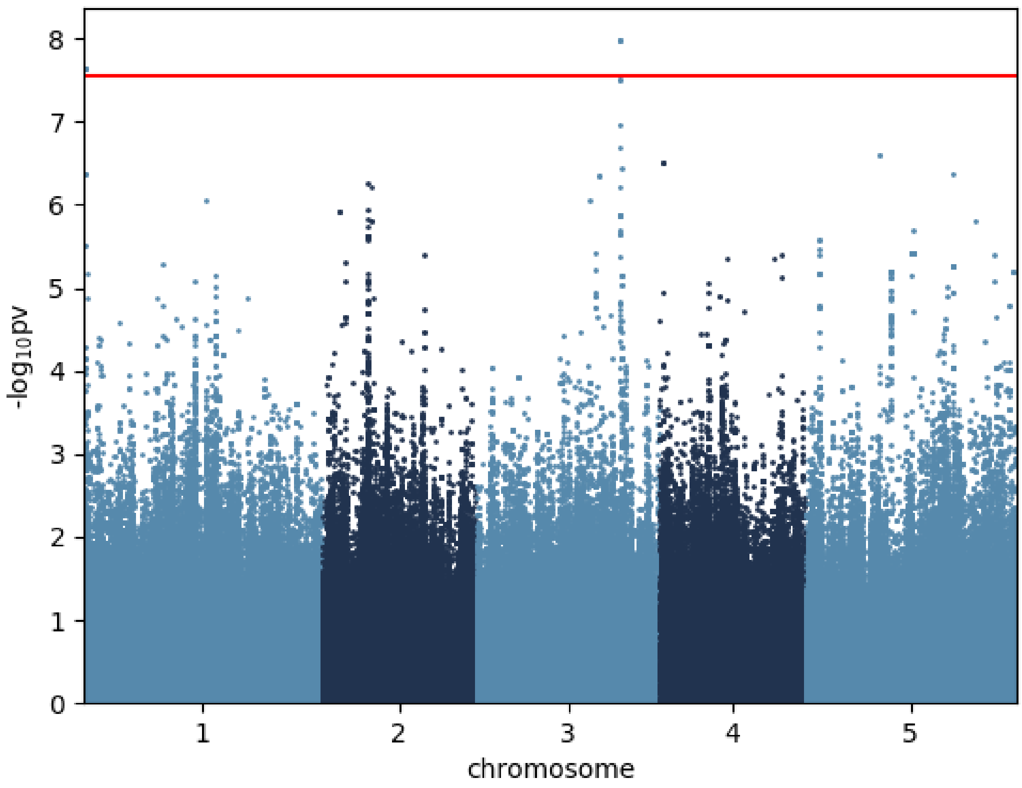


Glycine_16 °C

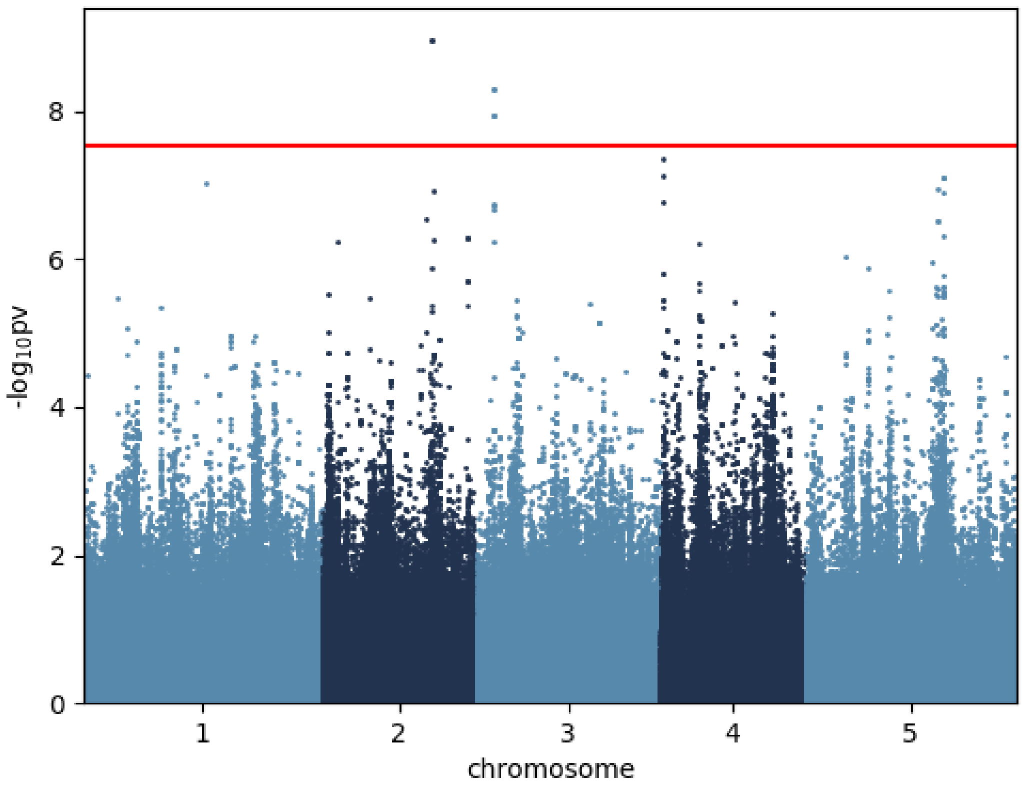


Isoleucine_16 °C

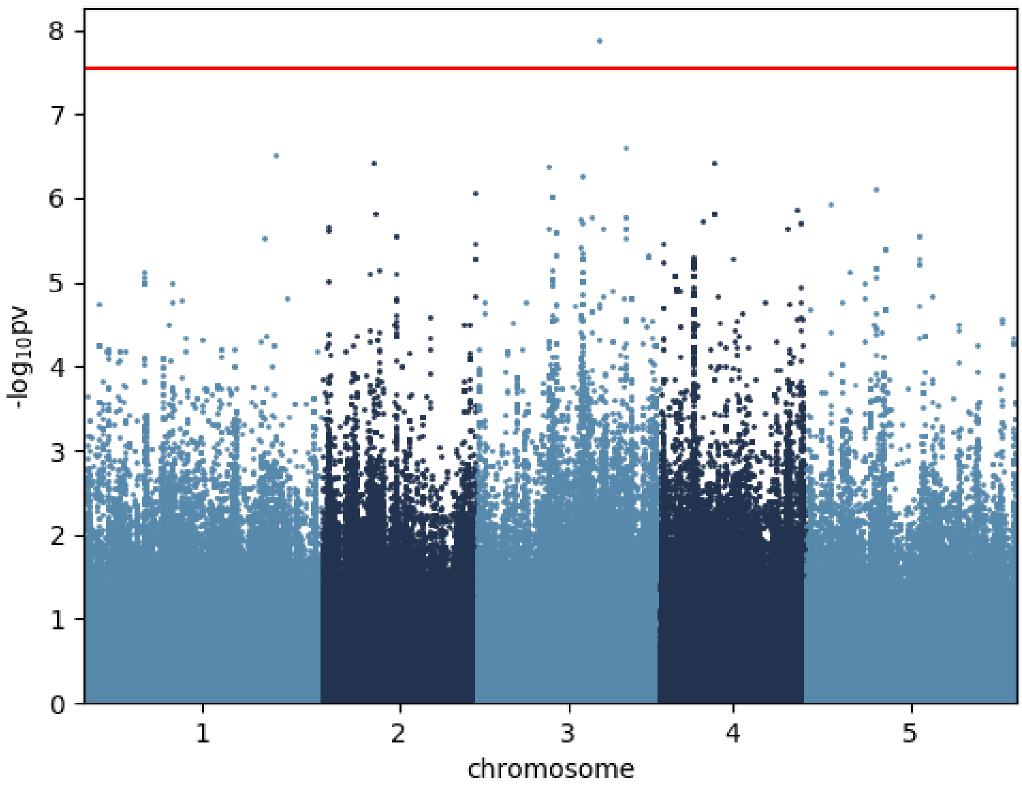


Lactic.acid_16 °C

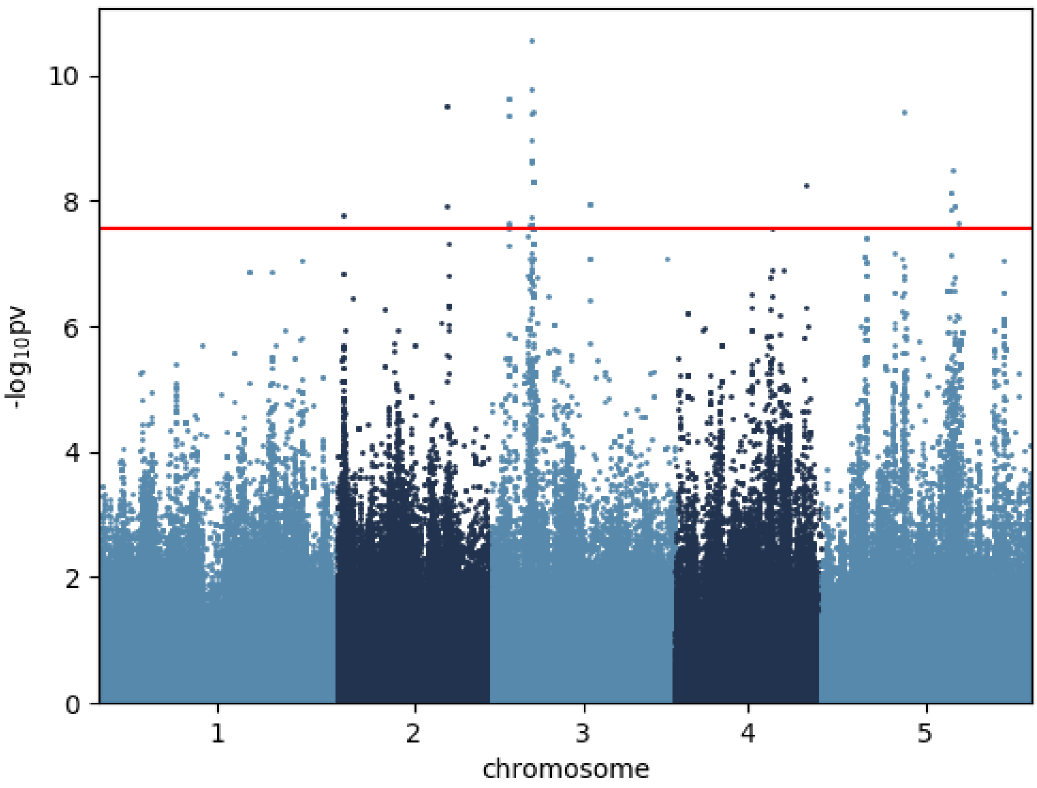


Leucine_16 °C

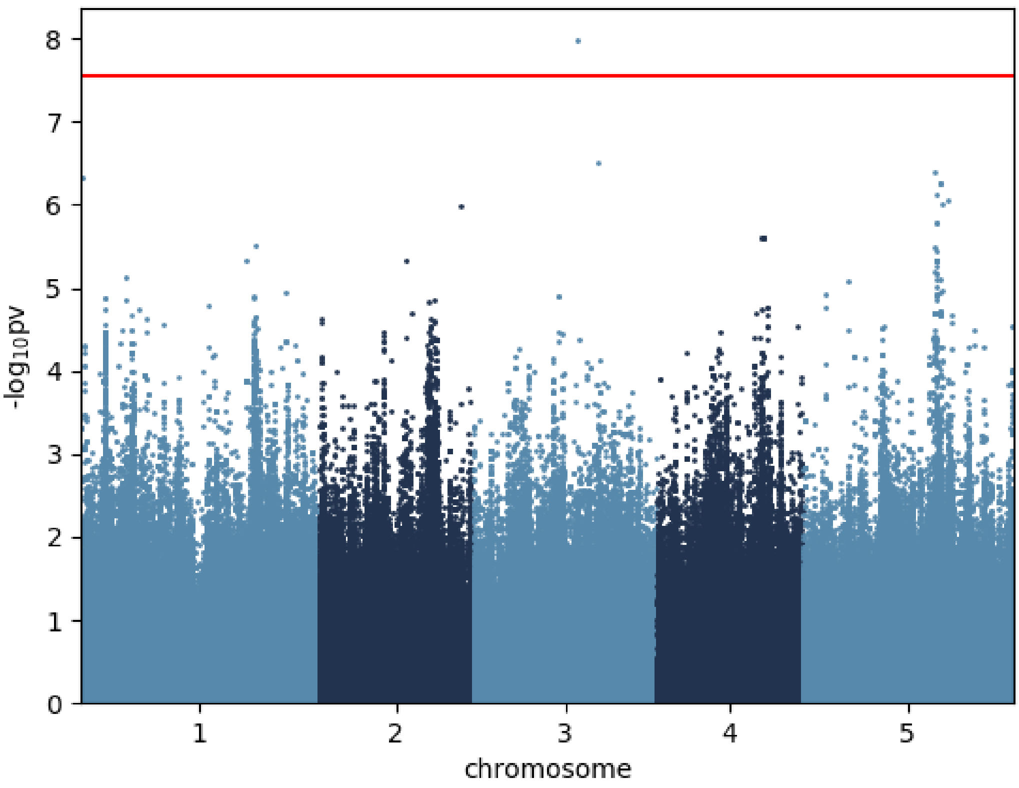


Lysine_16 °C

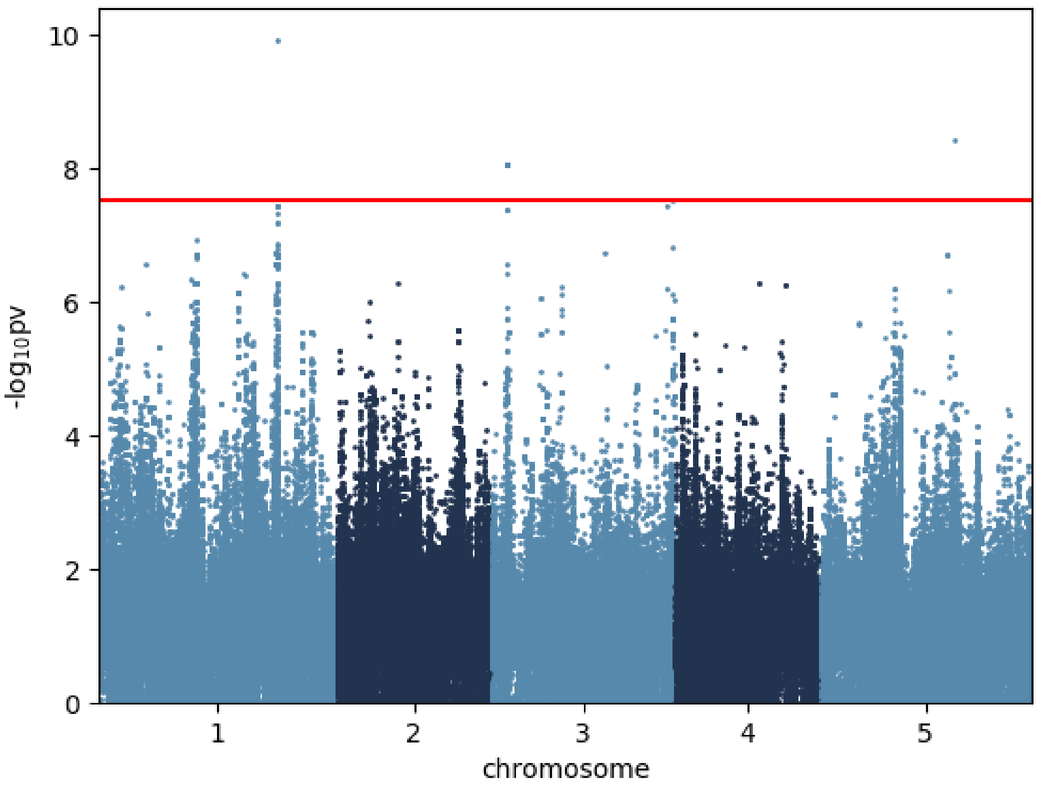


Malic.acid_16 °C

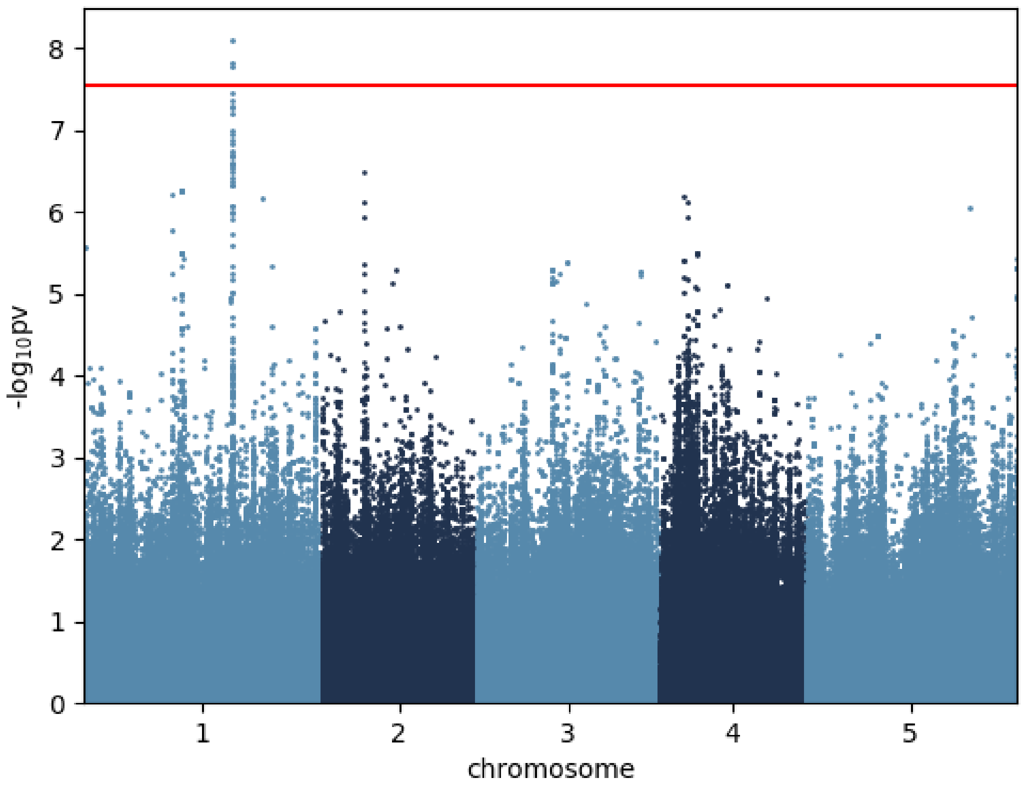


Maltose_16 °C

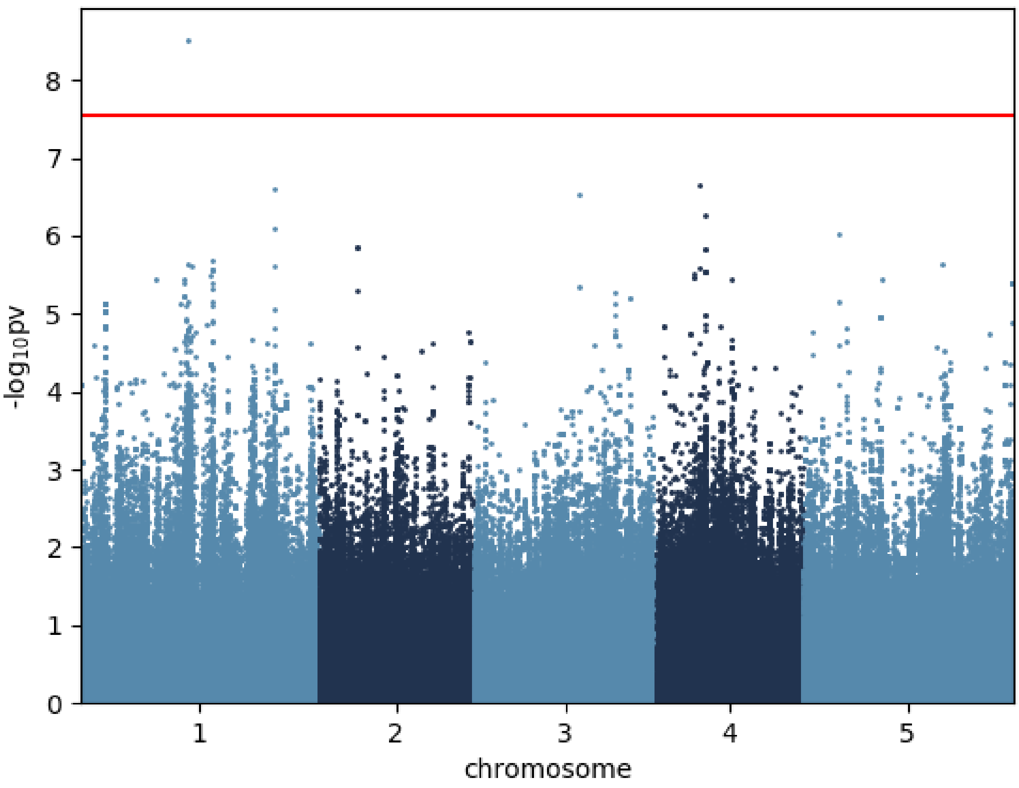


Myo.Inositol_16 °C

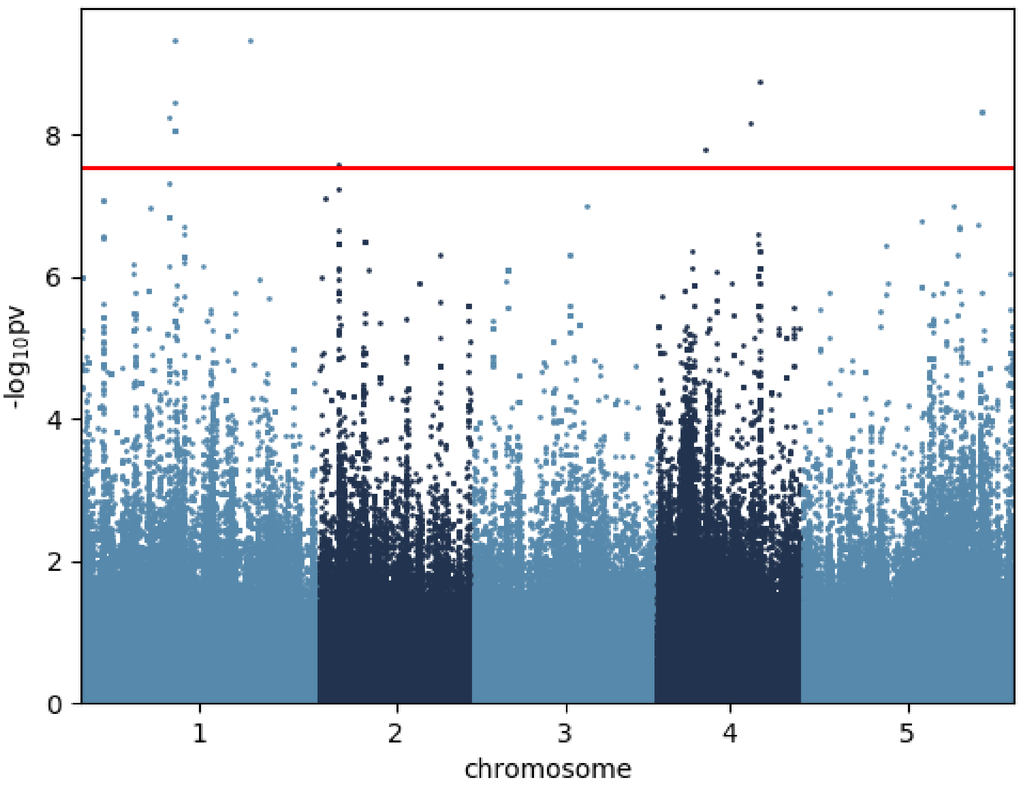


Ornithine_16 °C

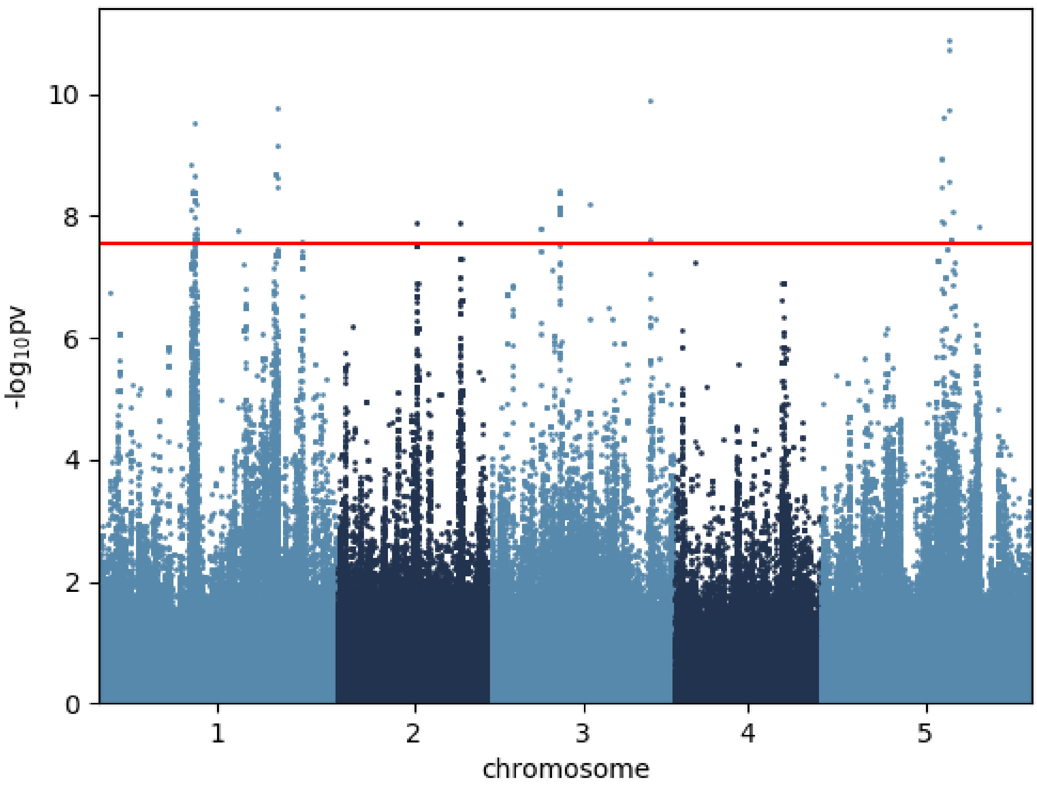


Oxoglutaric.acid_16 °C

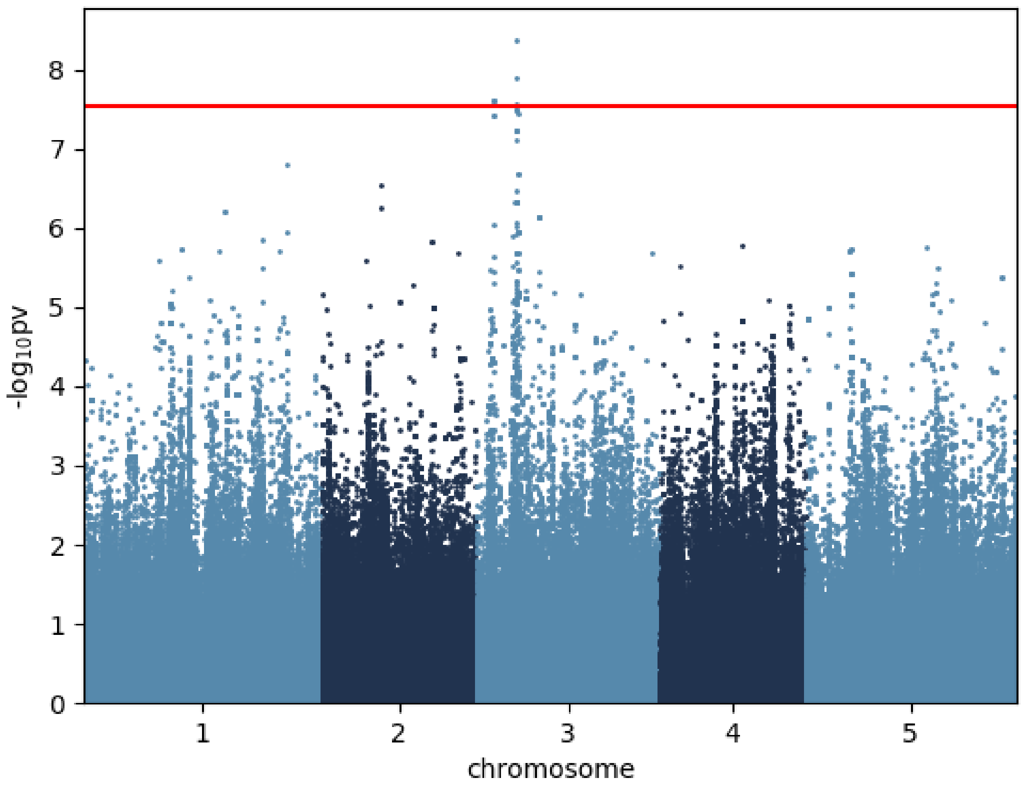


Phenylalanine_16 °C

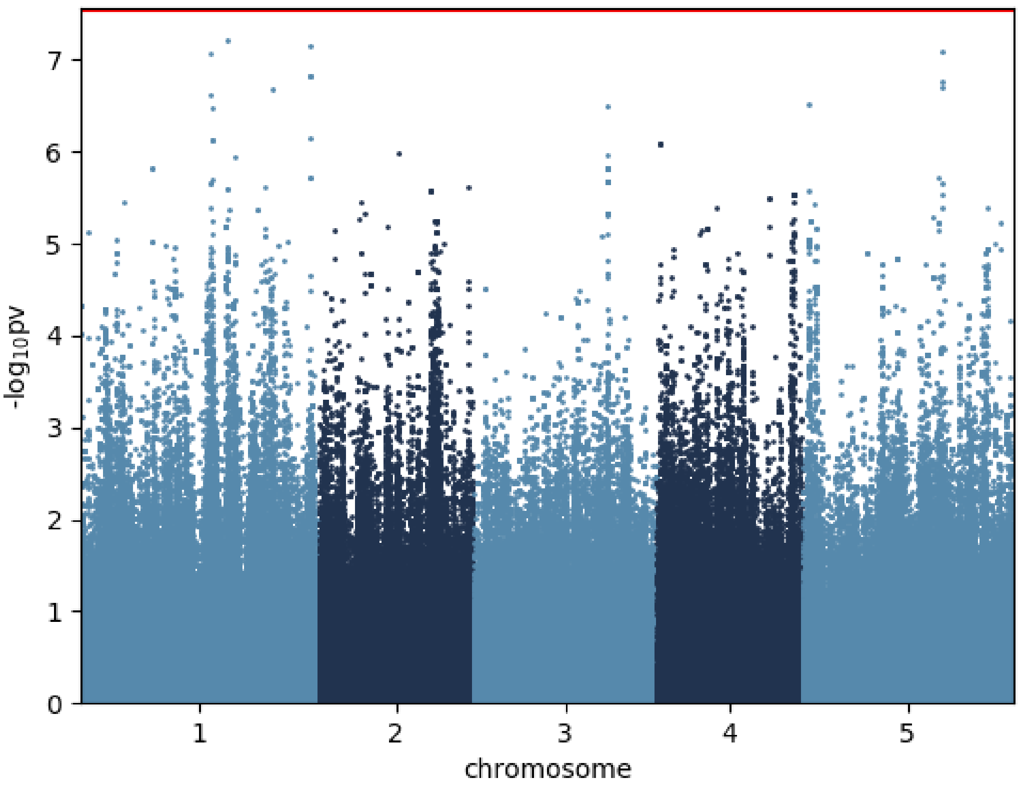


Proline_16 °C

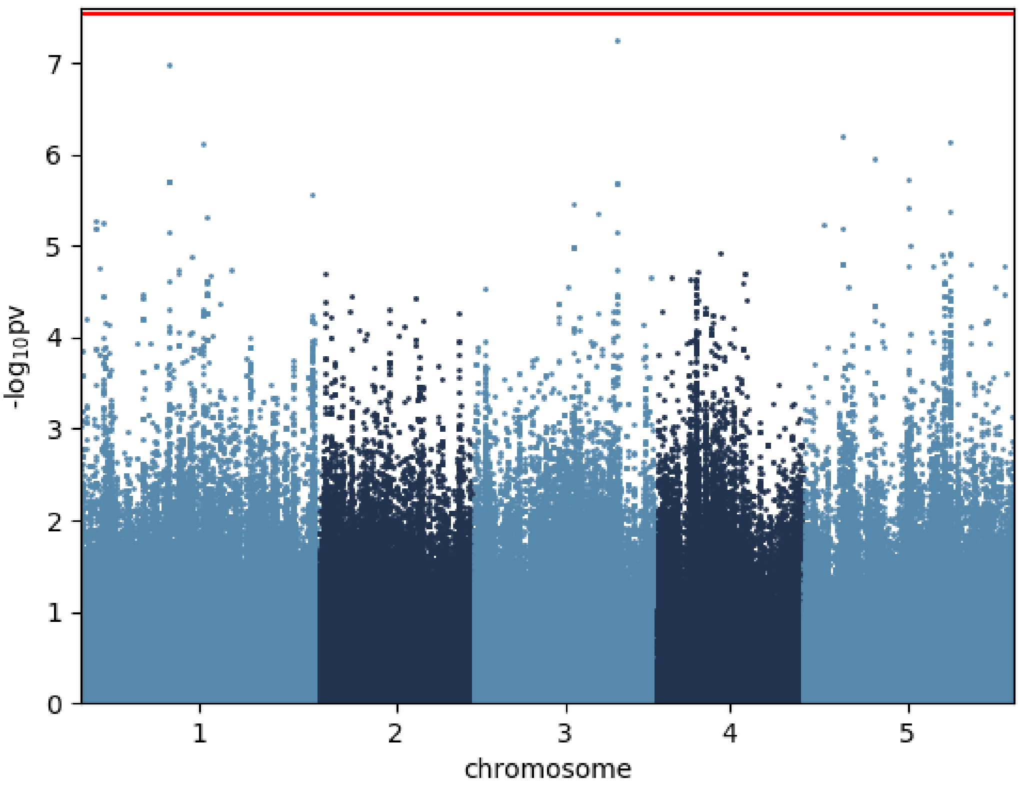


Putrescine_16 °C

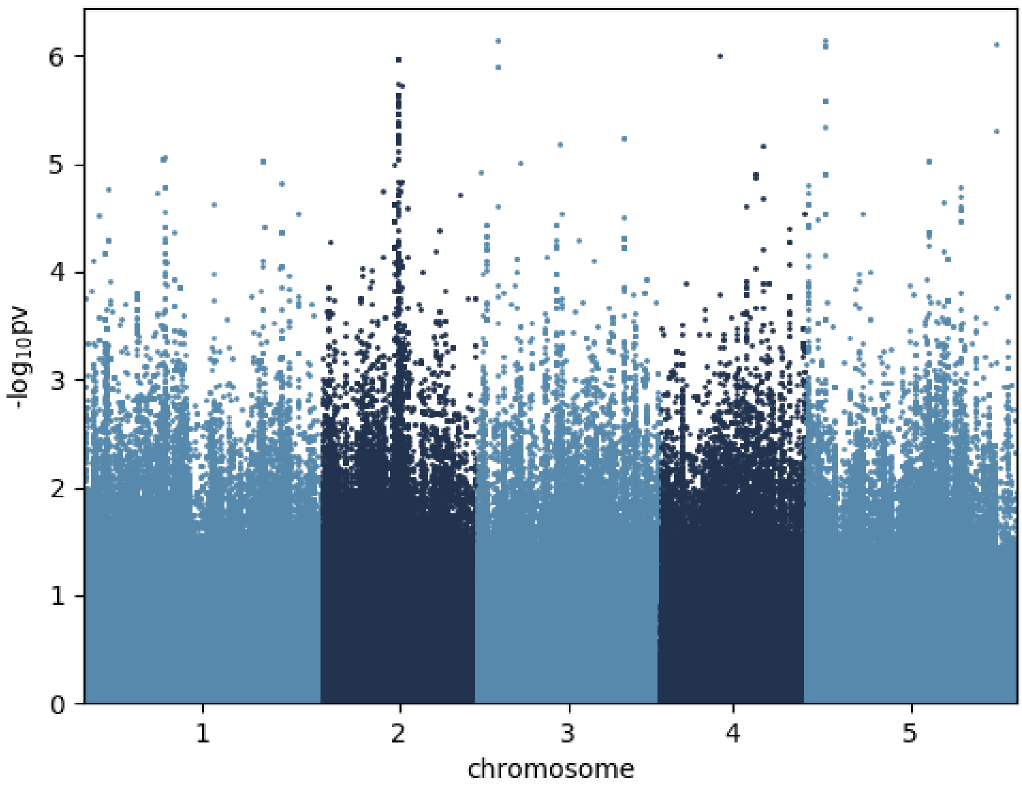


Pyruvic.acid_16 °C

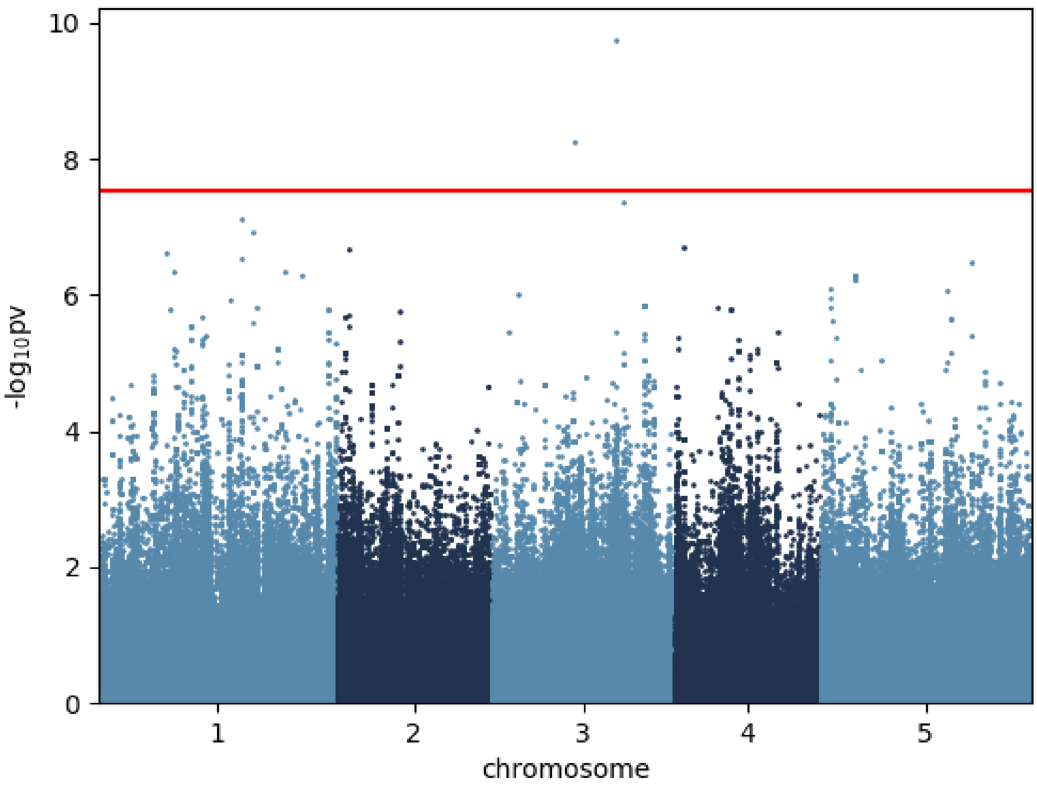


Raffinose_16 °C

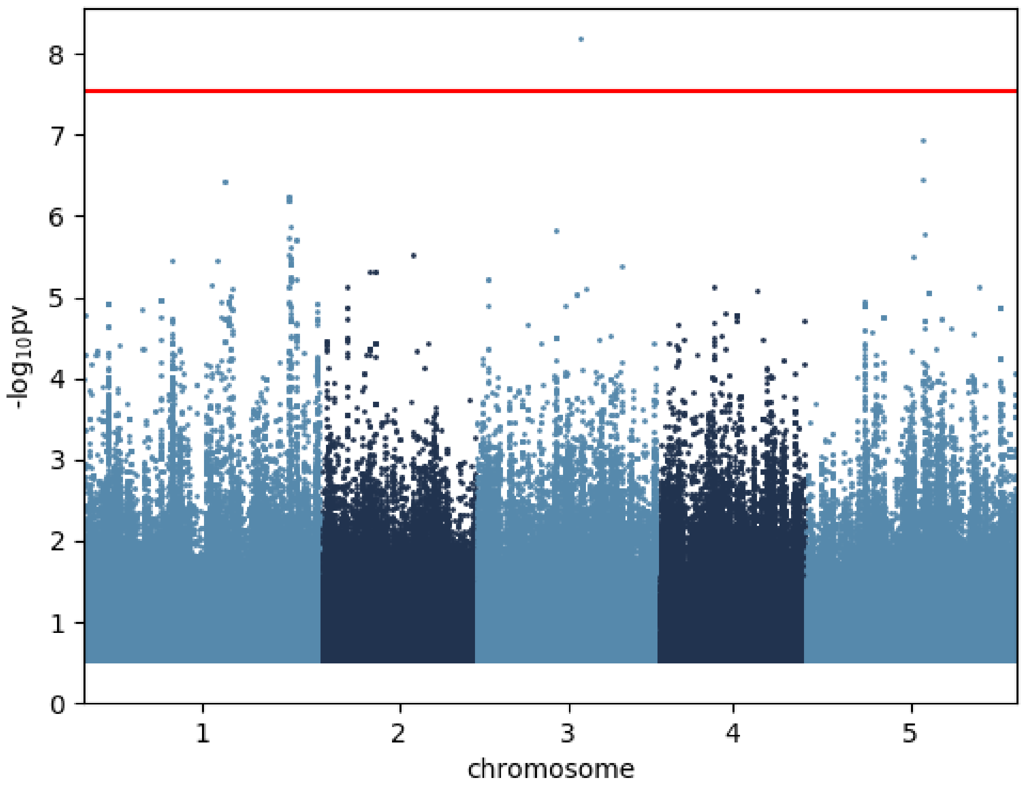


Serine_16 °C

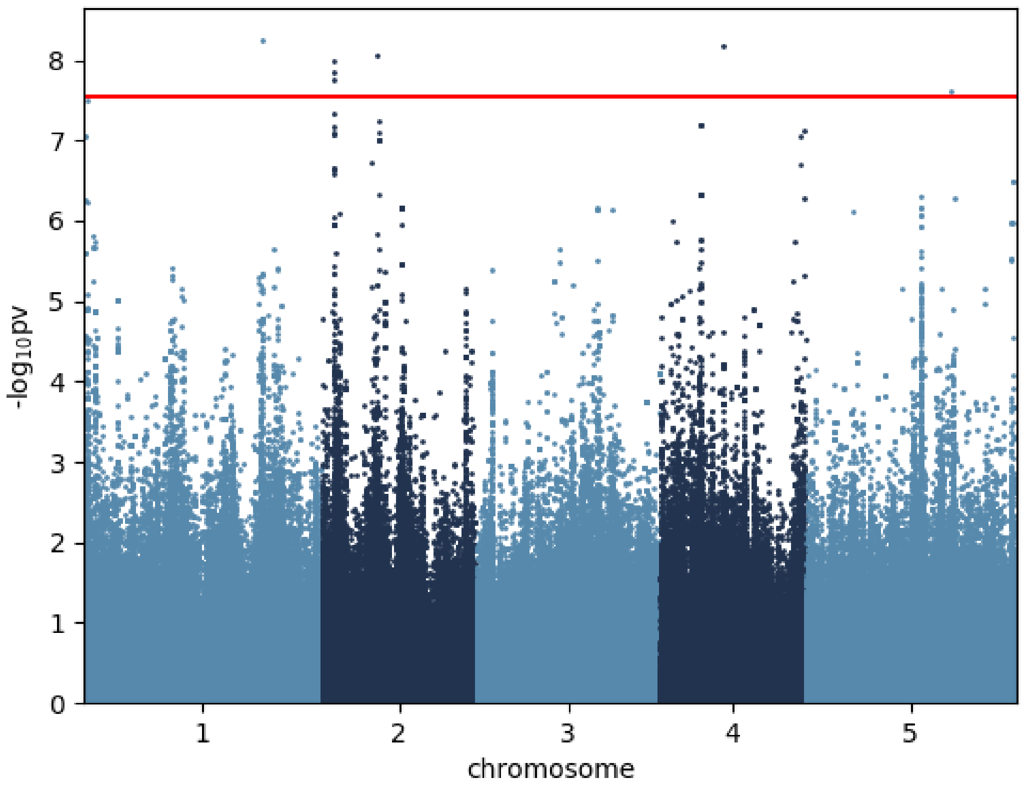


Spermidine_16 °C

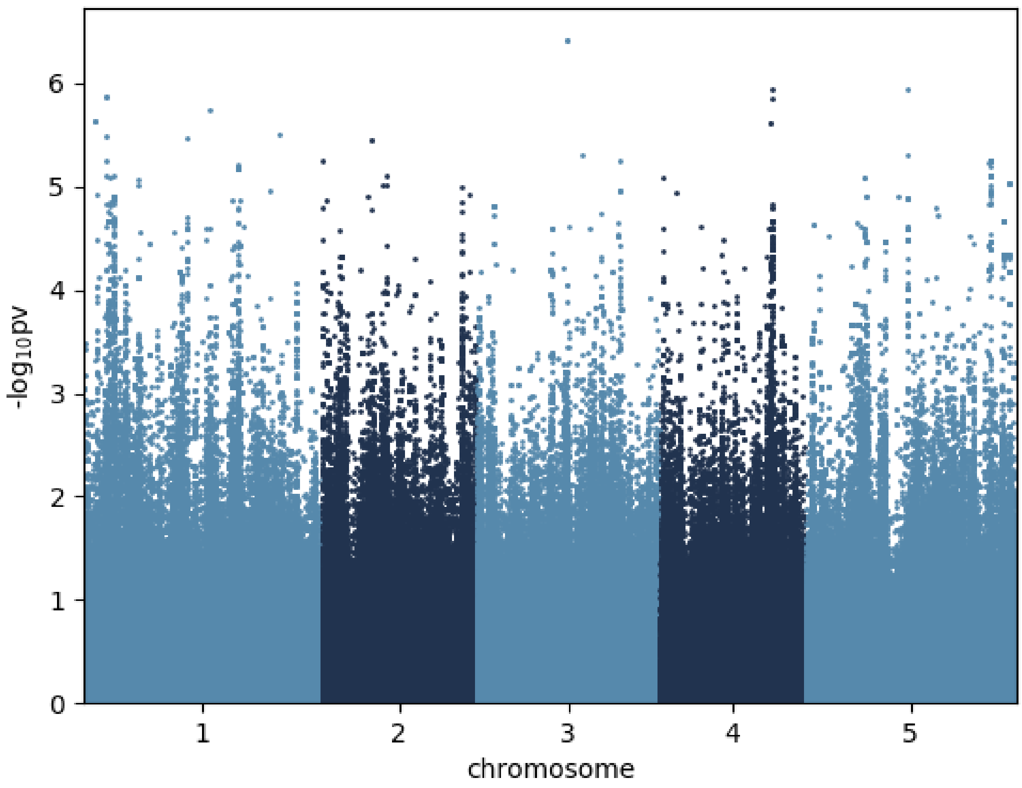


Succinic.acid_16 °C

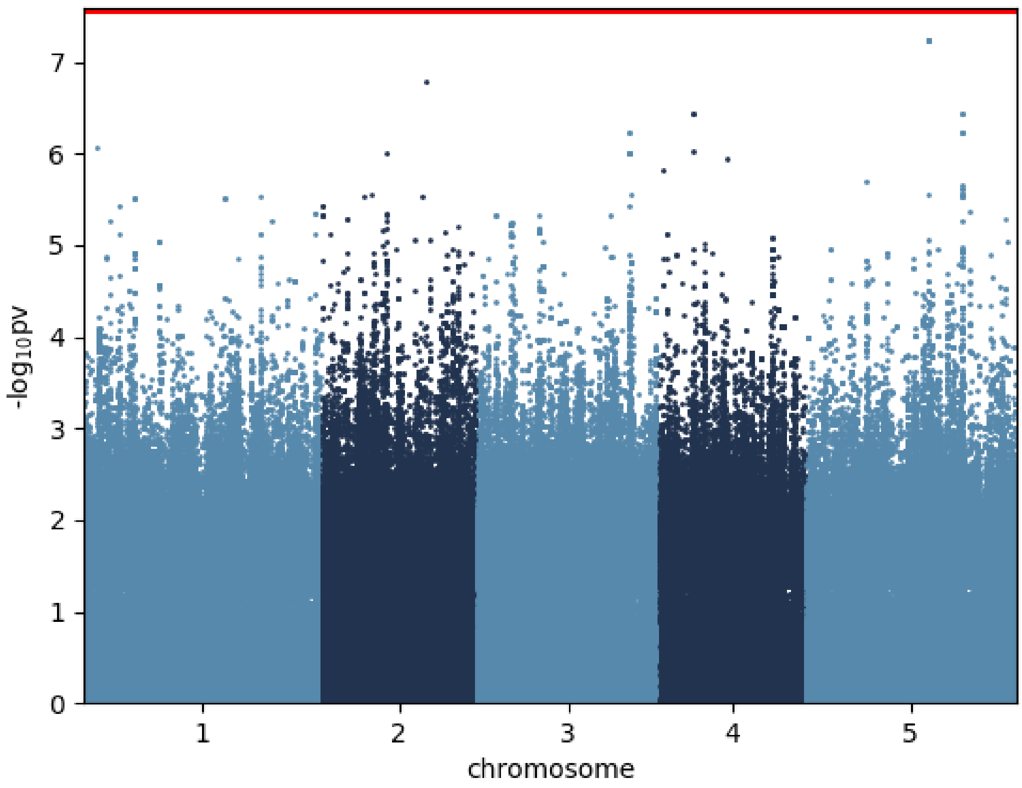


Sucrose_16 °C

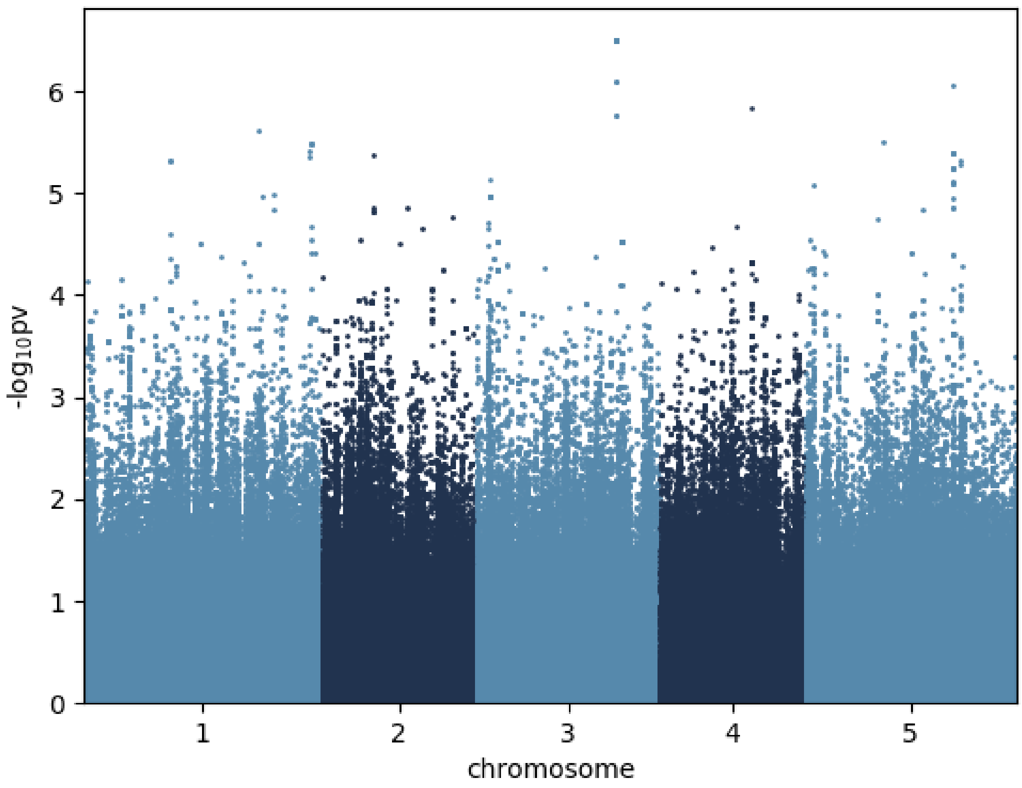


Threitol_16 °C

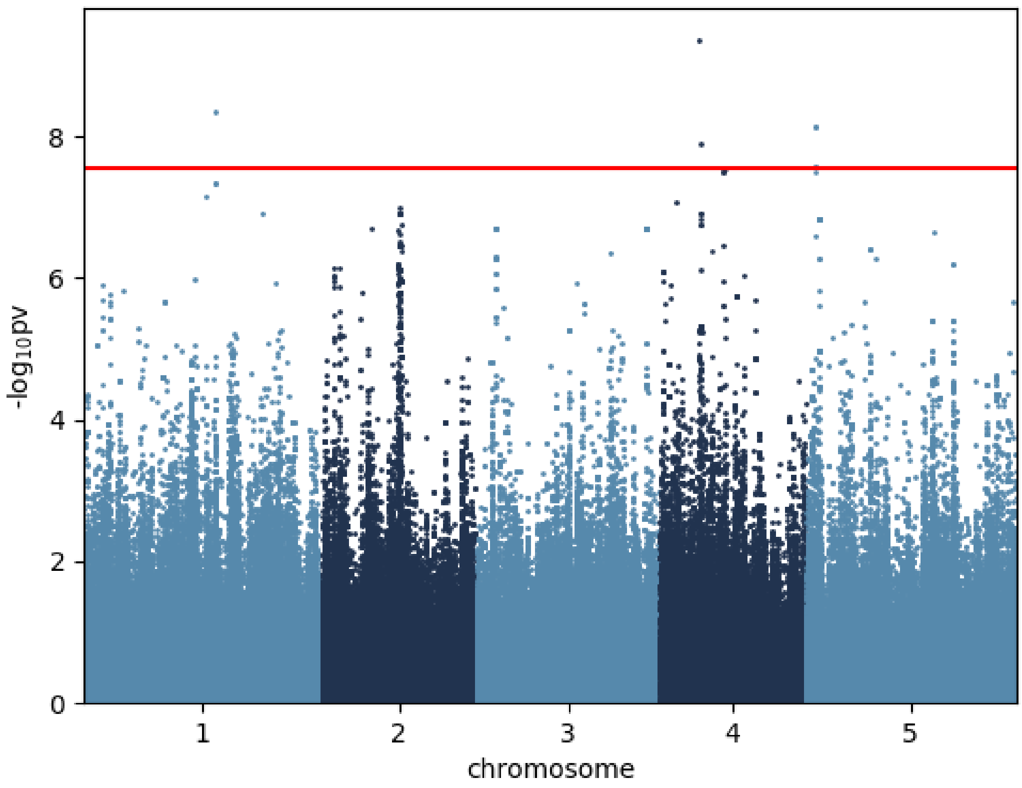


Threonic.acid_16 °C

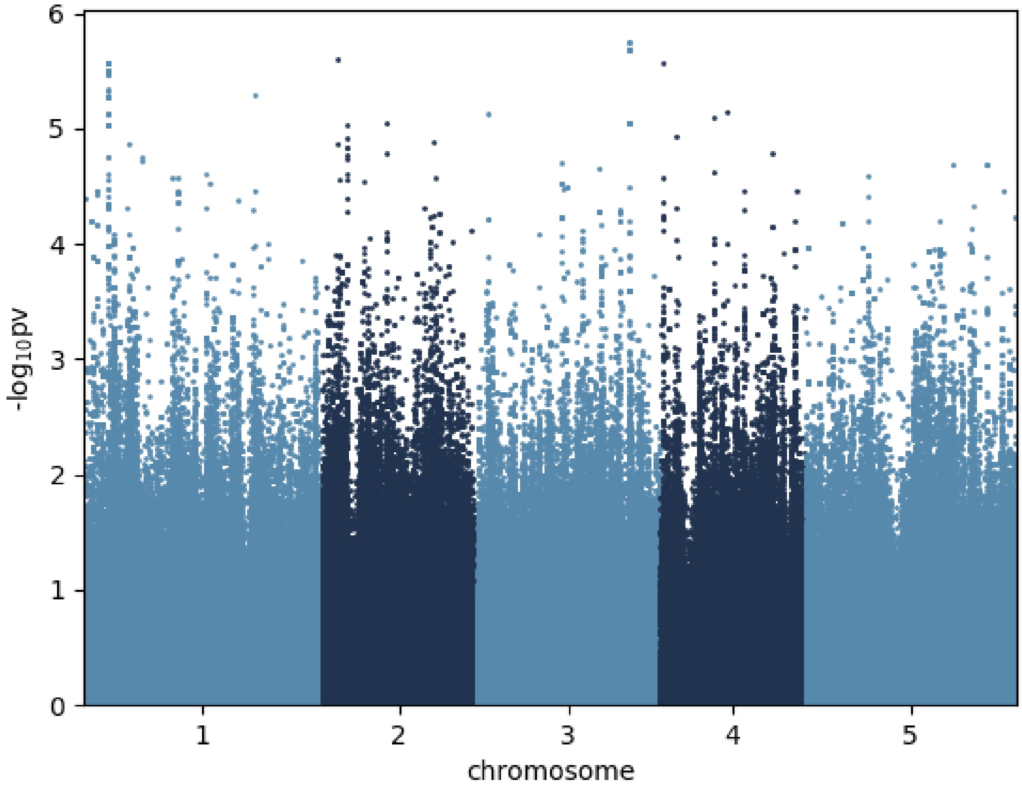


Threonine_16 °C

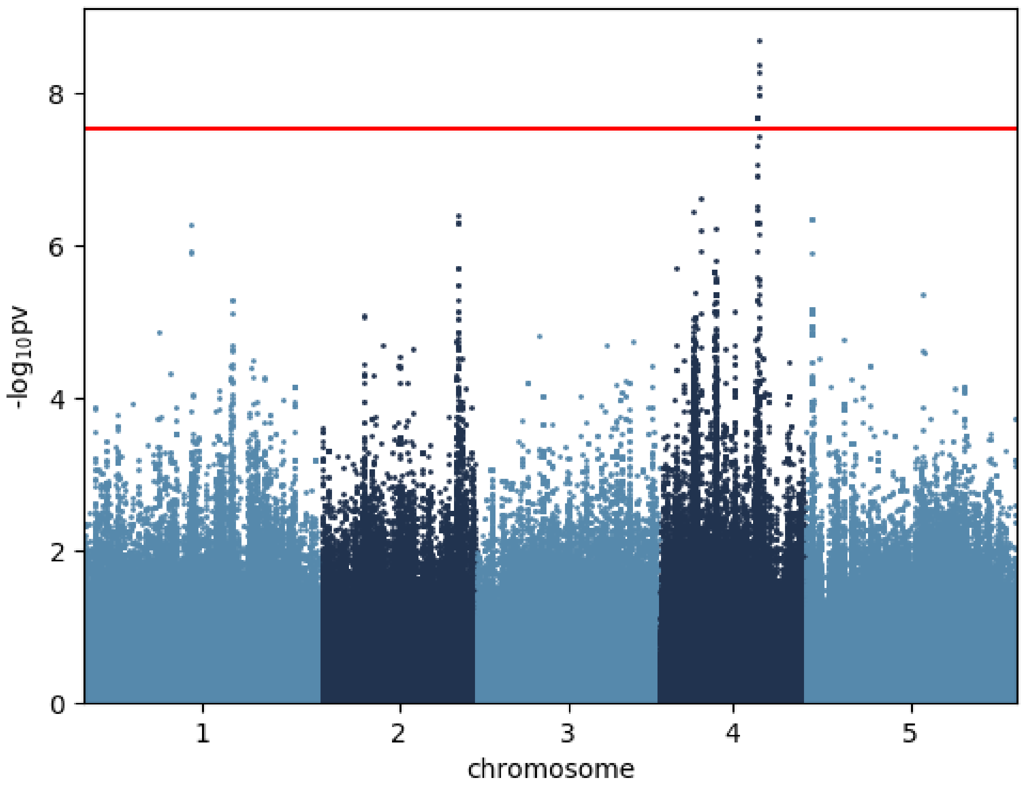


Trehalose_16 °C

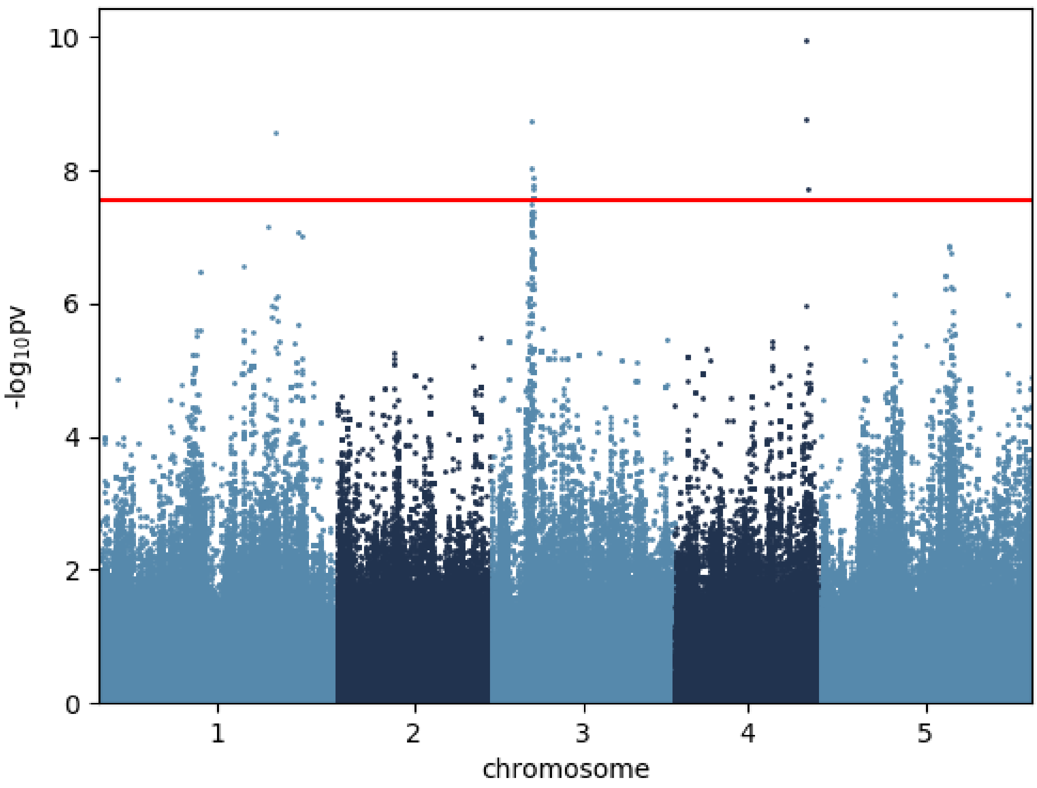


Tyrosine_16 °C

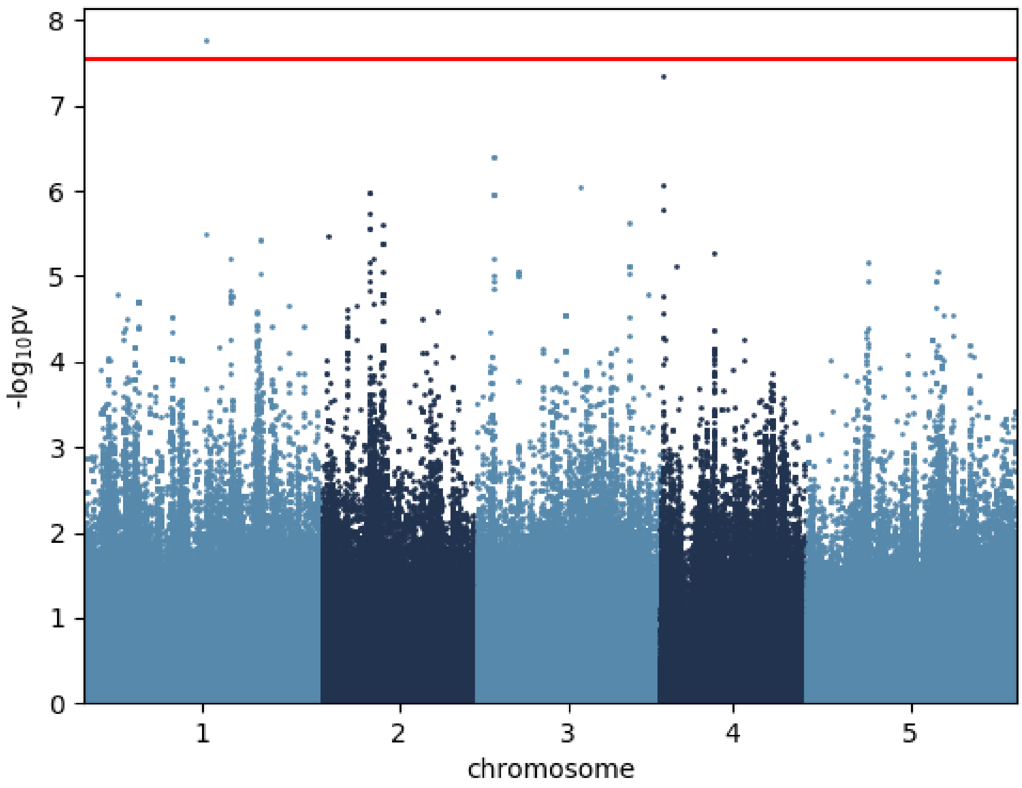


Valine_16 °C

mGWAS of metabolite concentrations in the 6 °C condition, red line indicates significance threshold after Bonferroni correction:

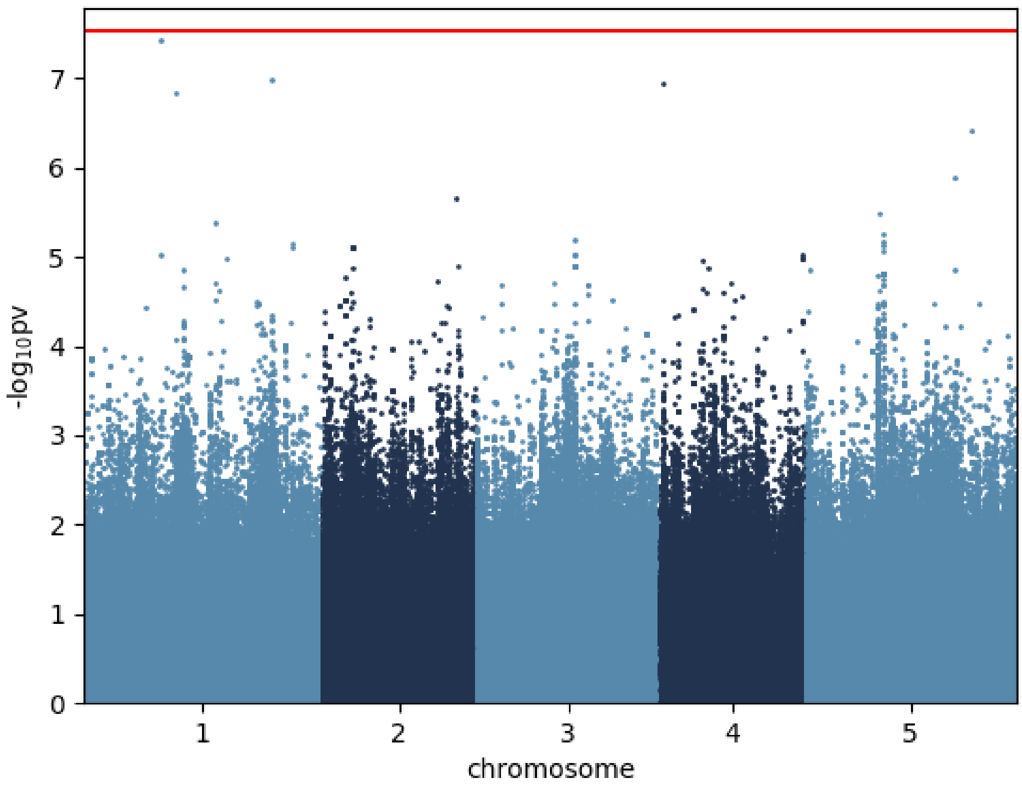


Alanine_6 °C

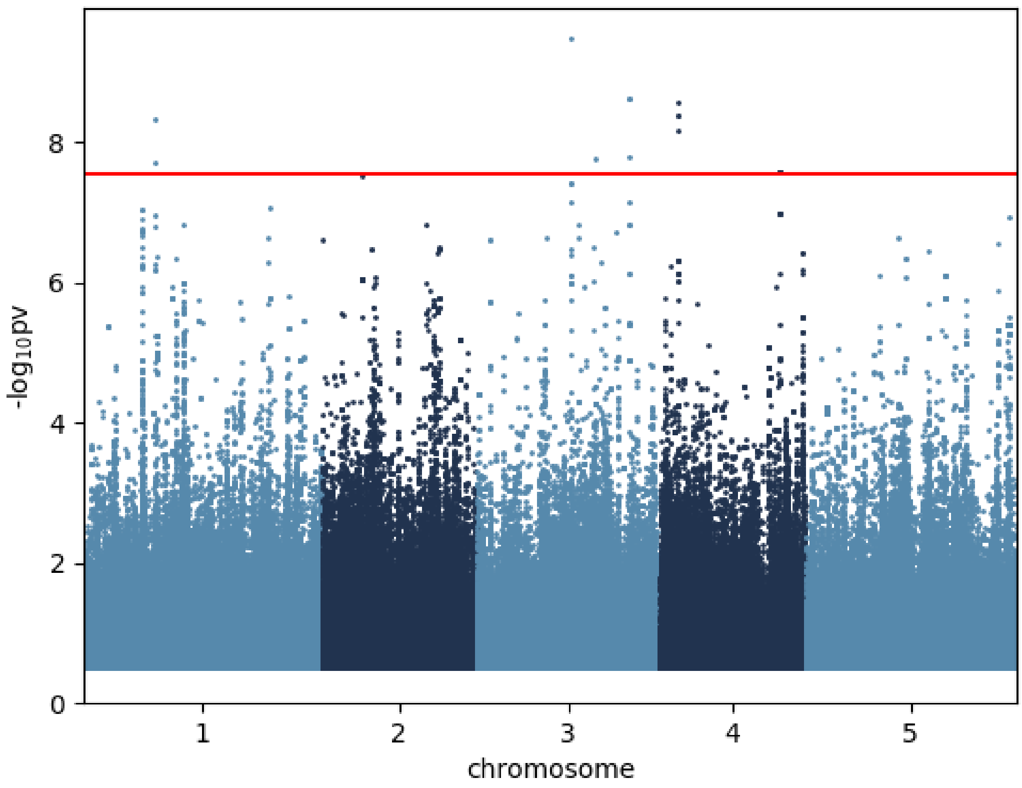


Asparagine_6 °C

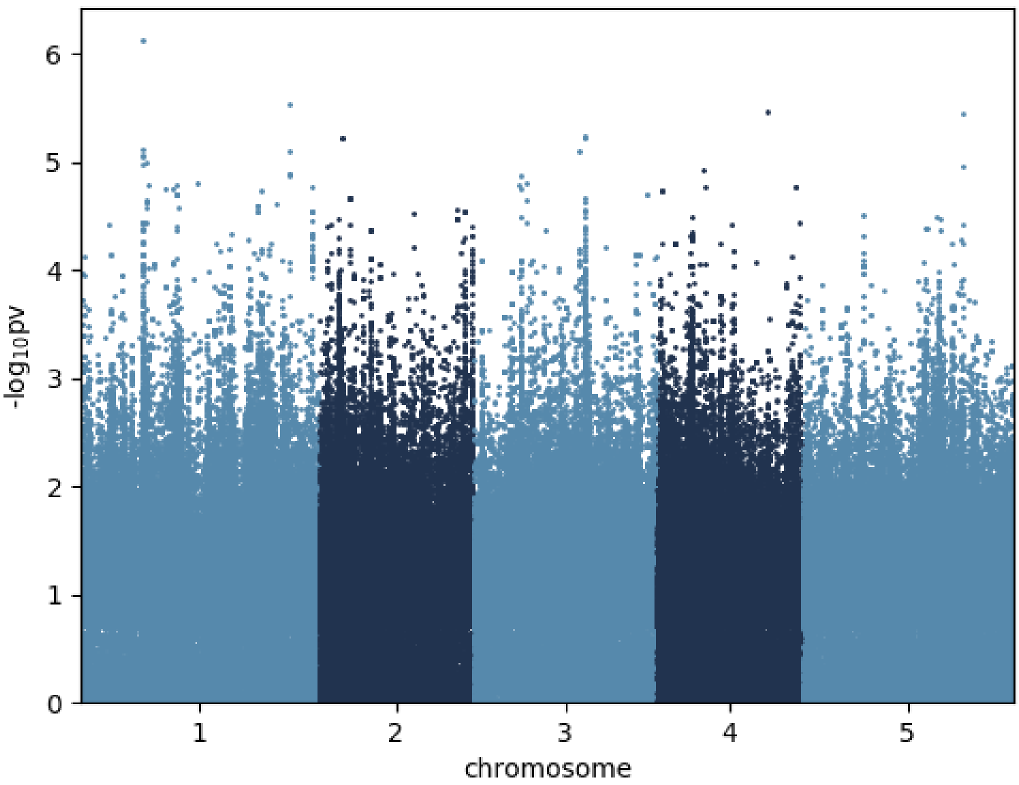


Aspartic.acid_6 °C

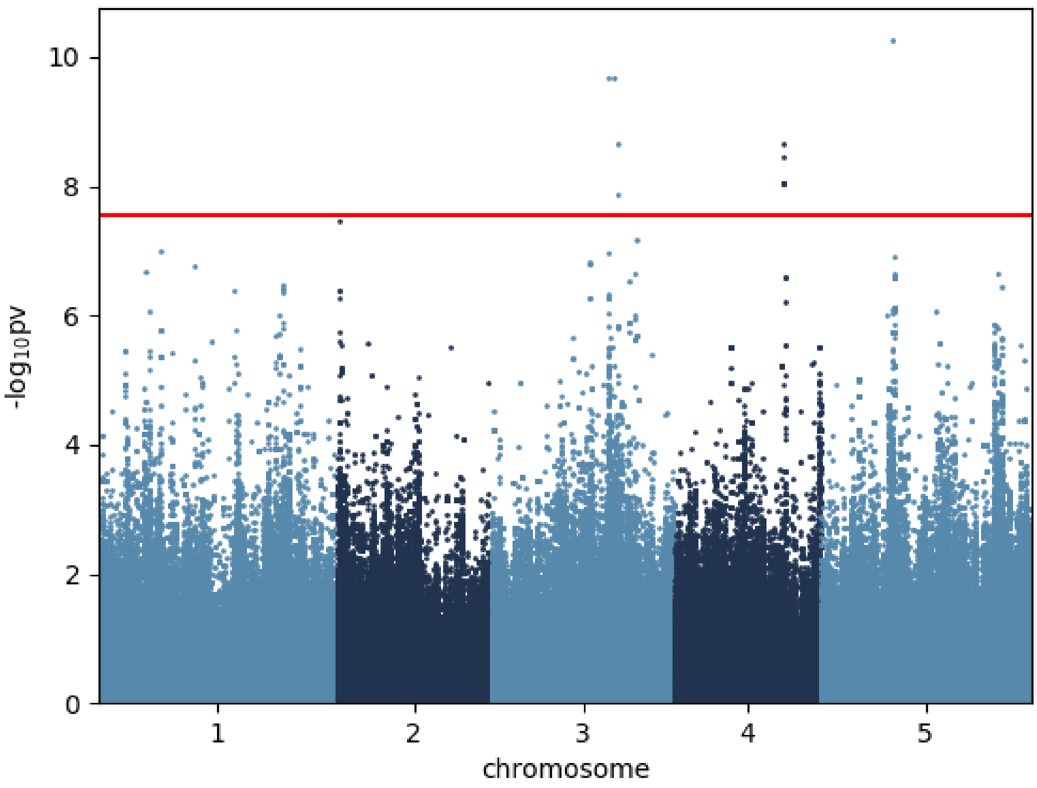


Butanoic.acid.4.amino._6 °C

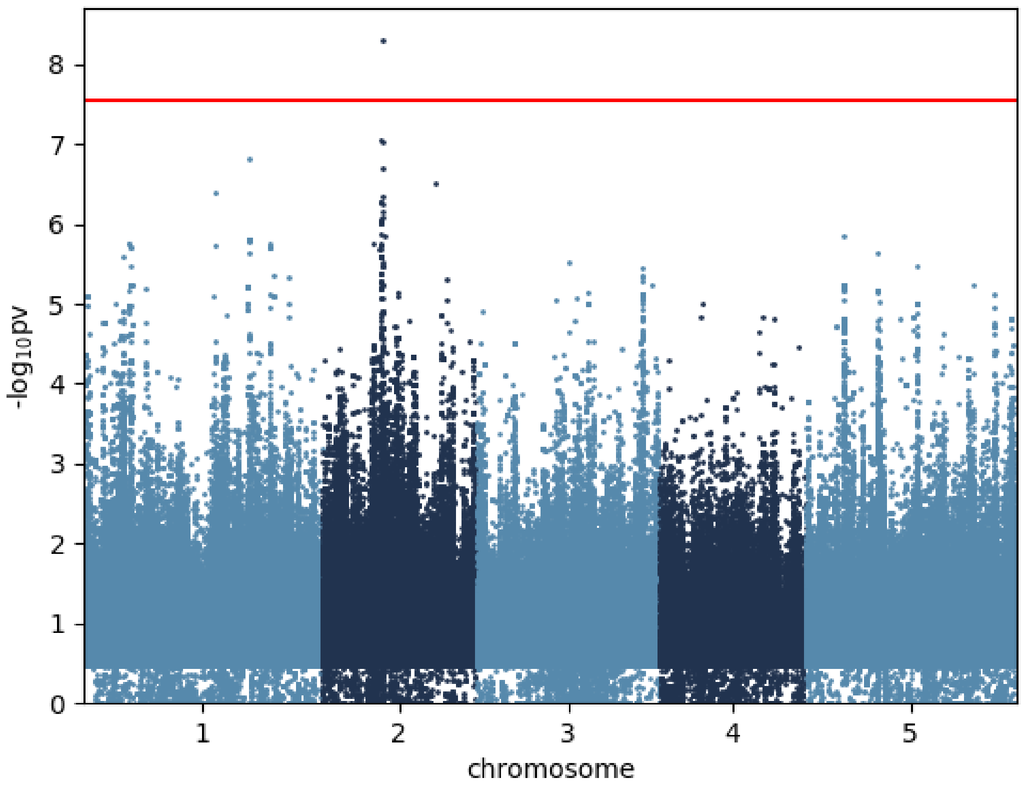


Citric.acid_6 °C

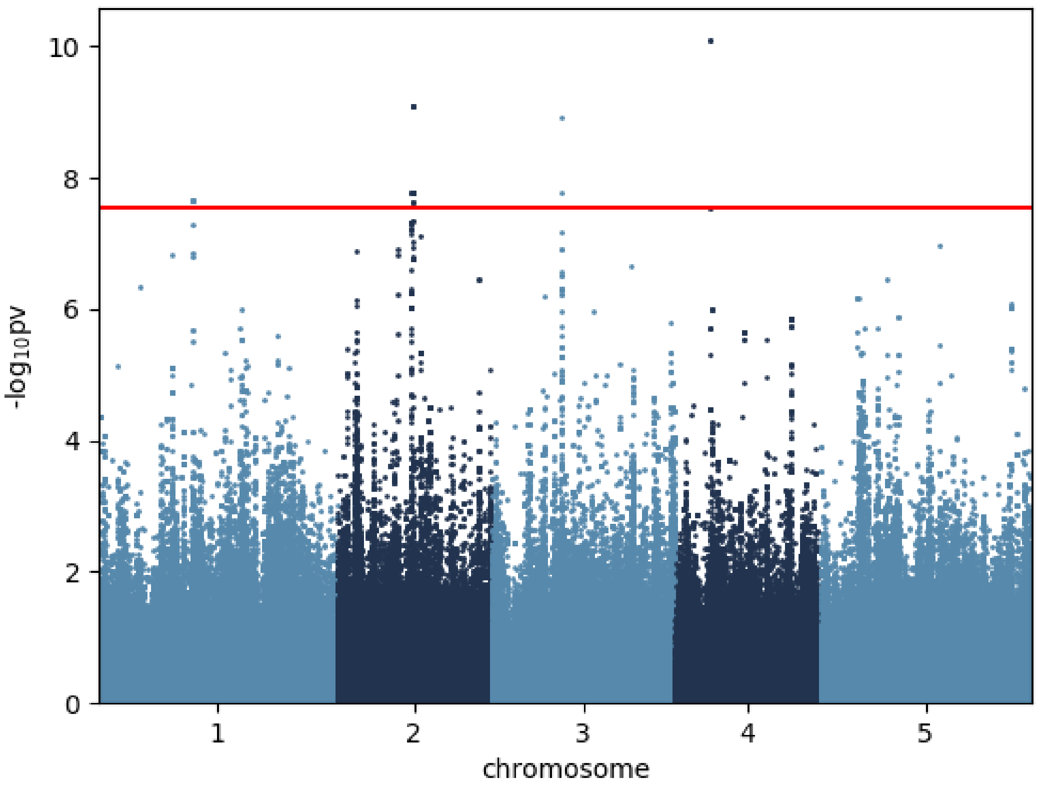


Fructose_6 °C

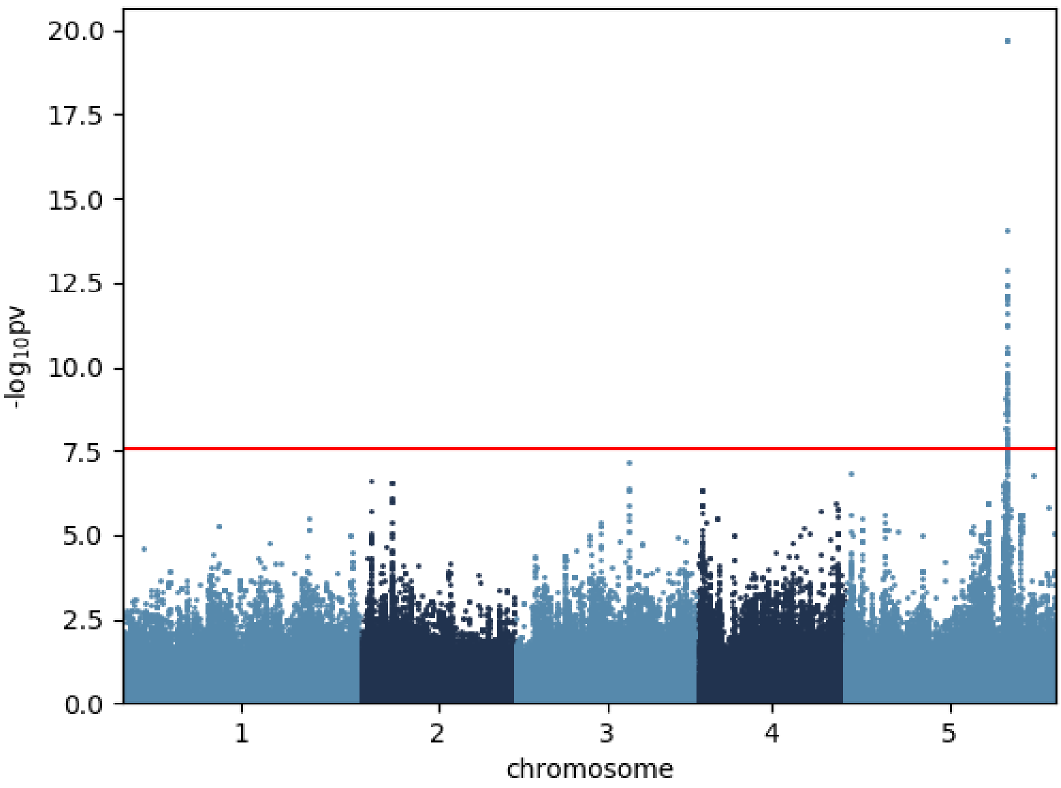


Fumaric.acid_6 °C

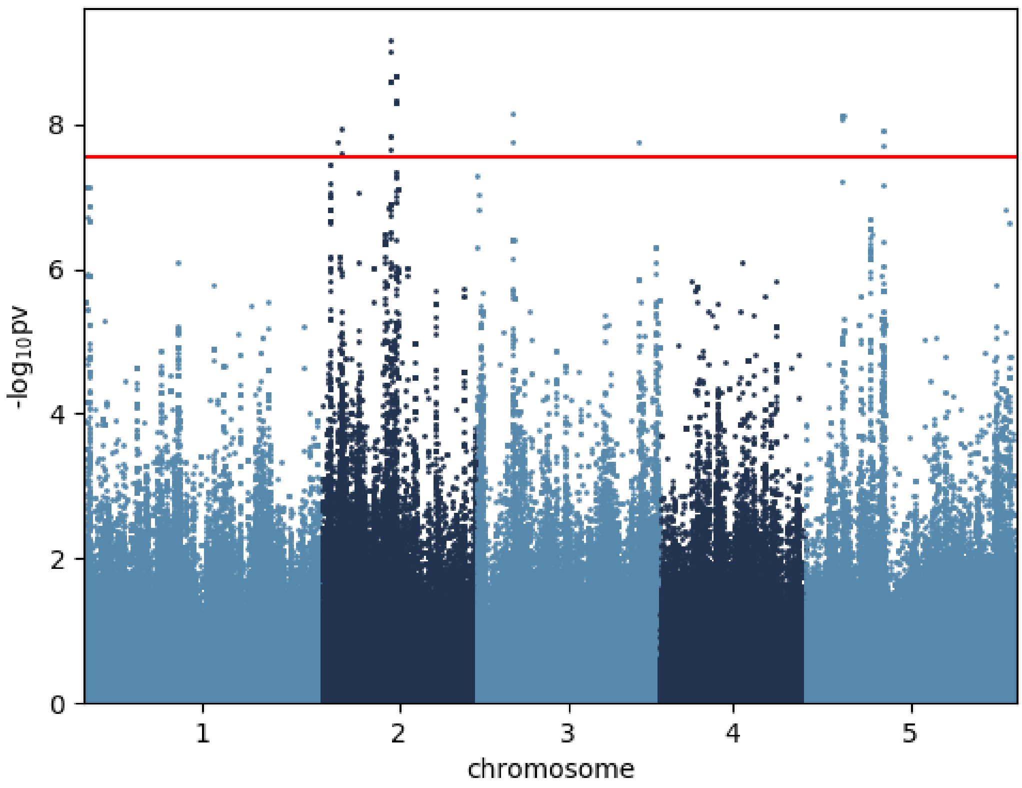


Galactinol_6 °C

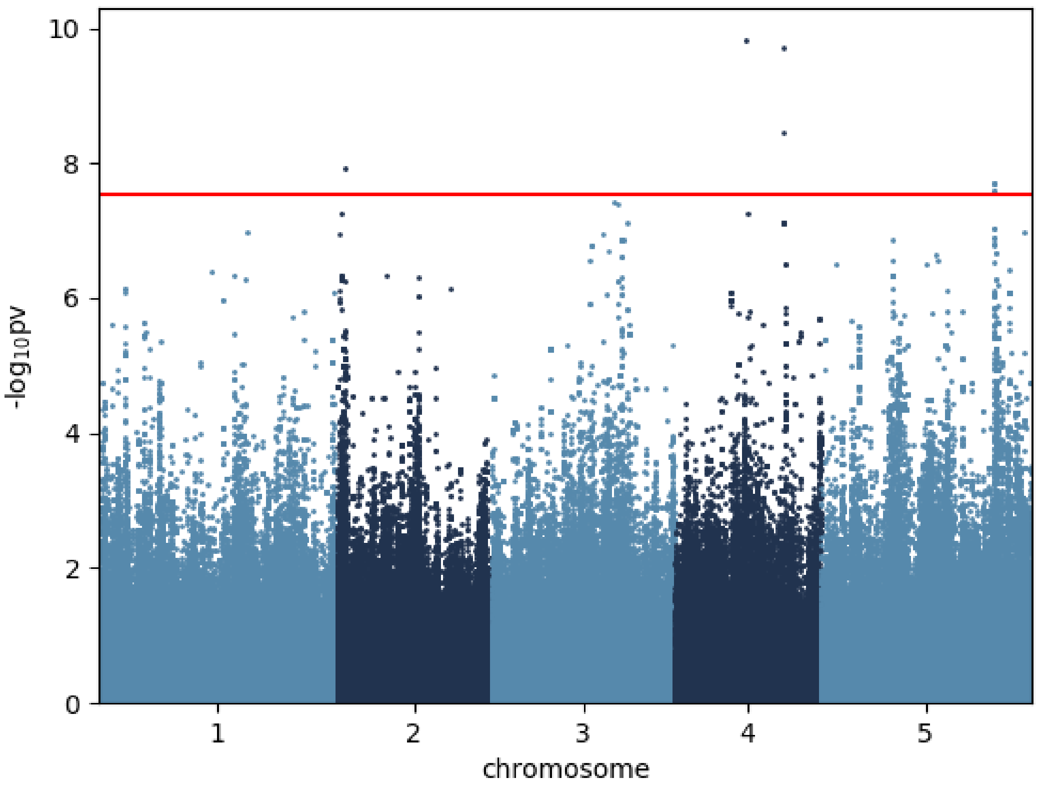


Galactose_6 °C

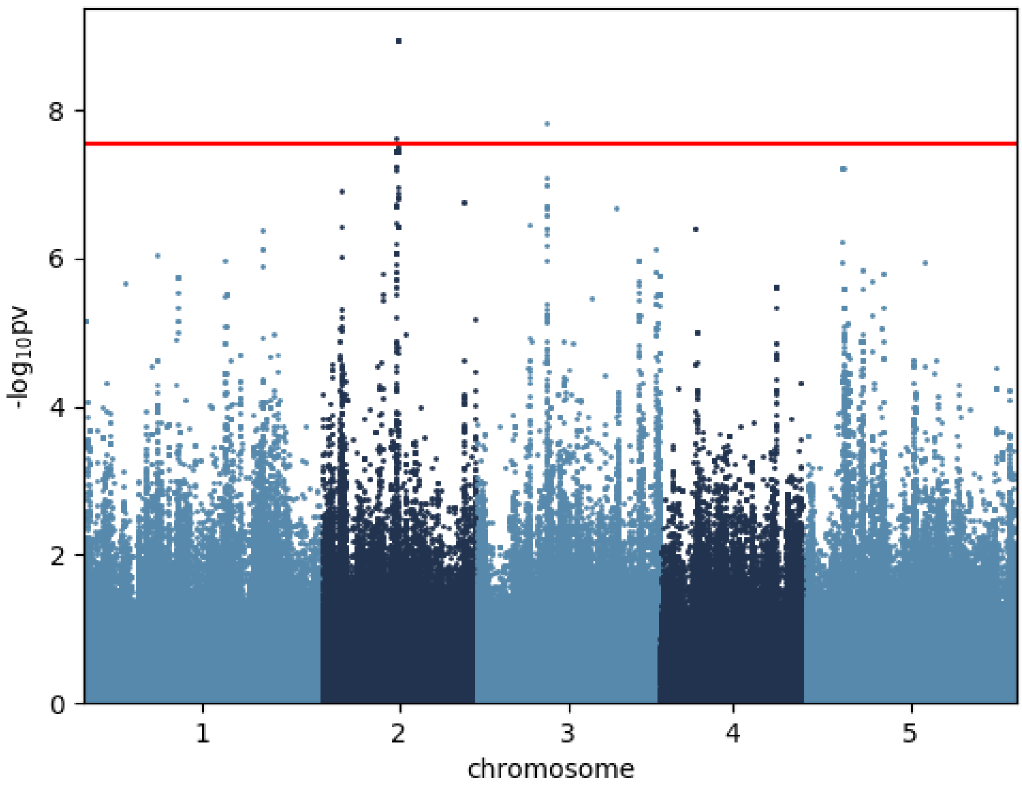


Glucose_6 °C

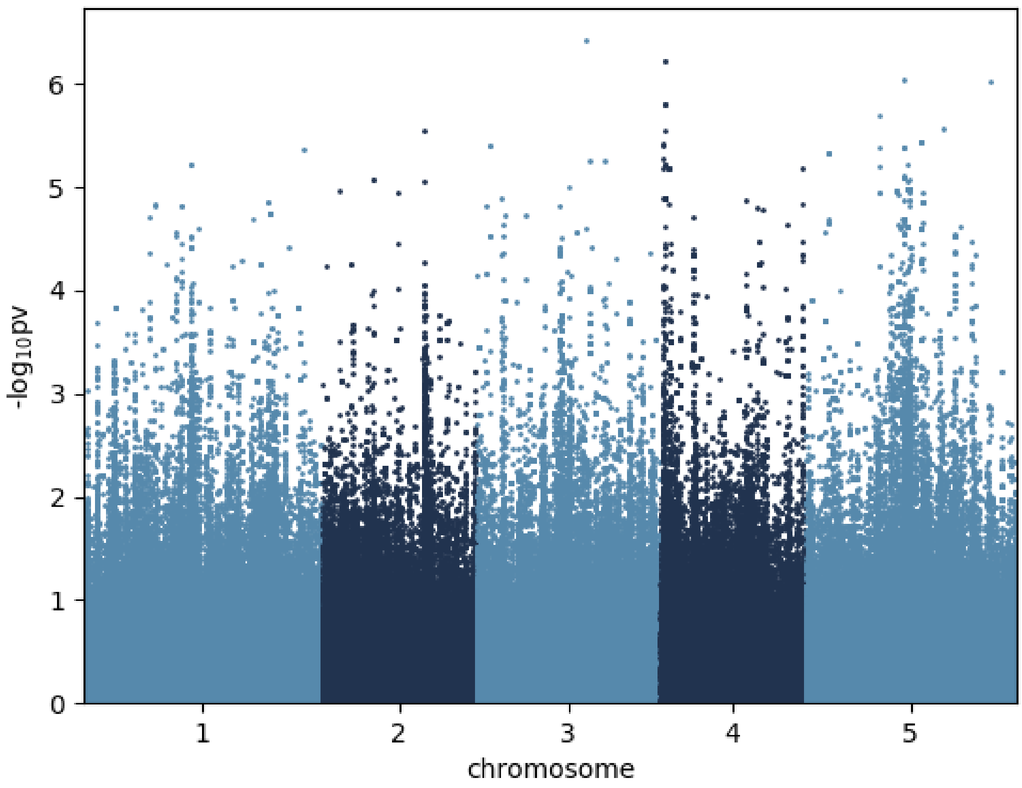


Glutamic.acid_6 °C

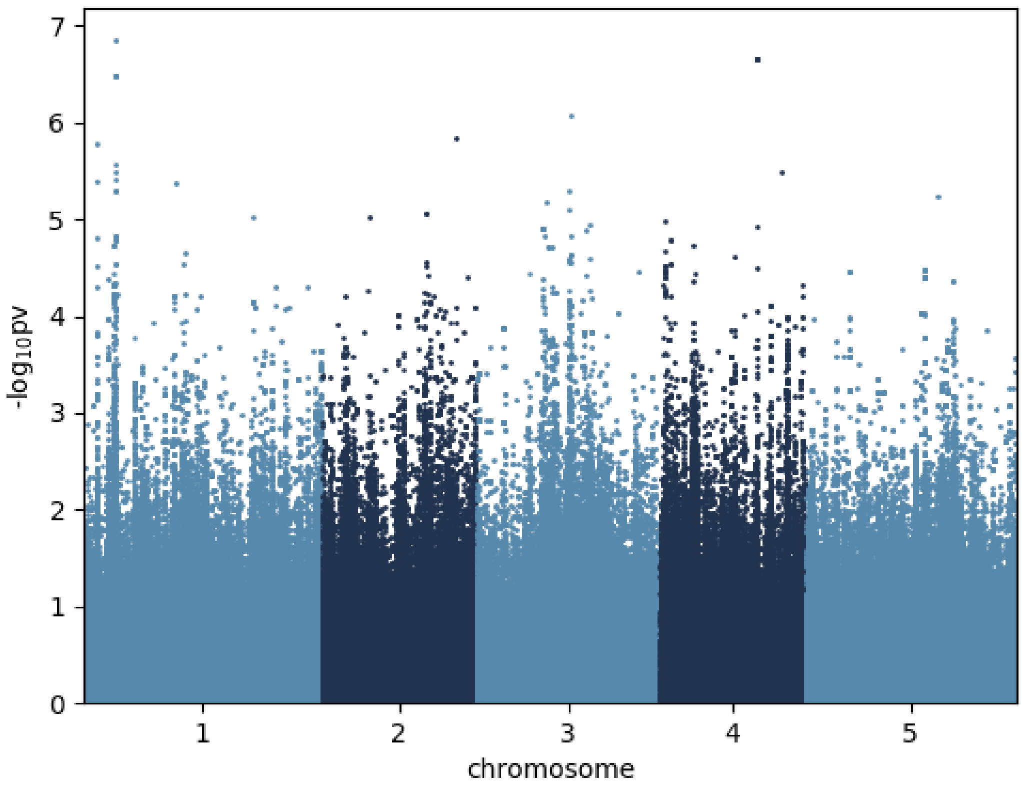


Glutamine_6 °C

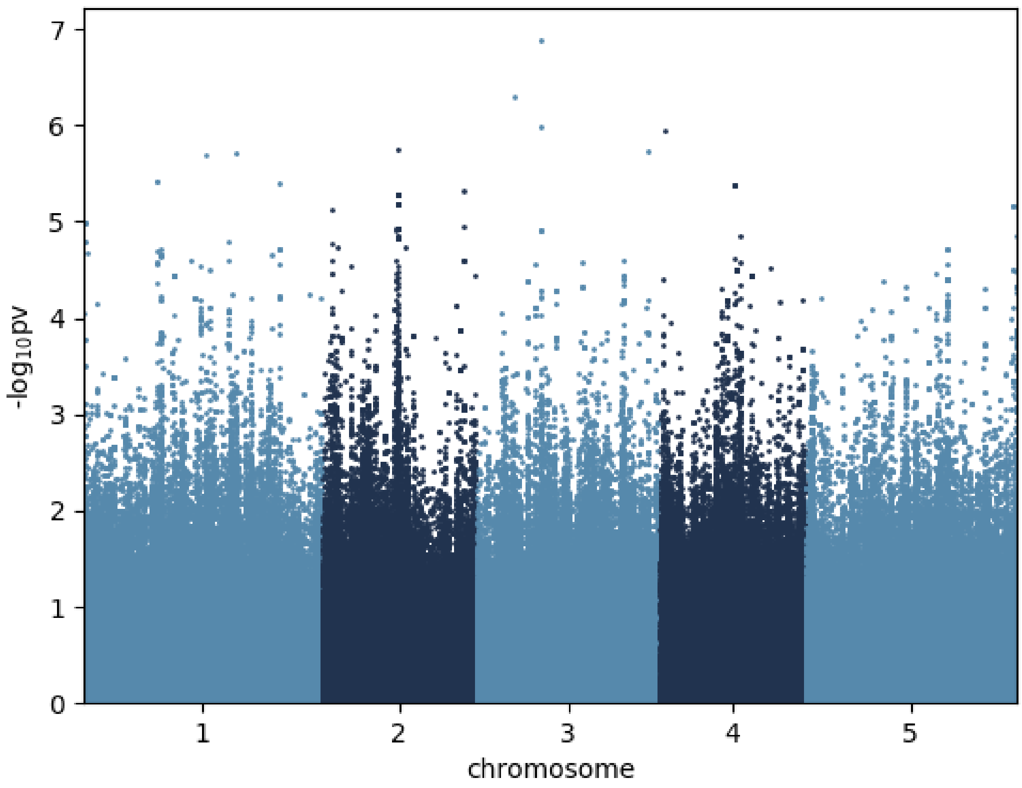


Glycine_6 °C

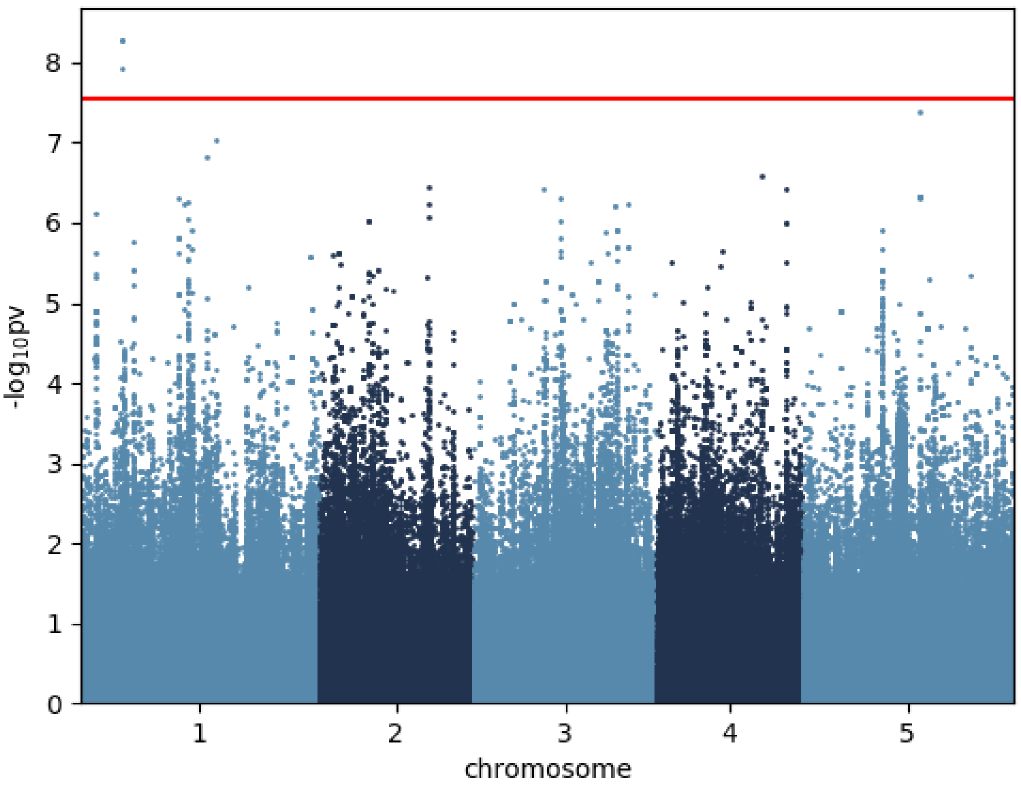


Lactic.acid_6 °C

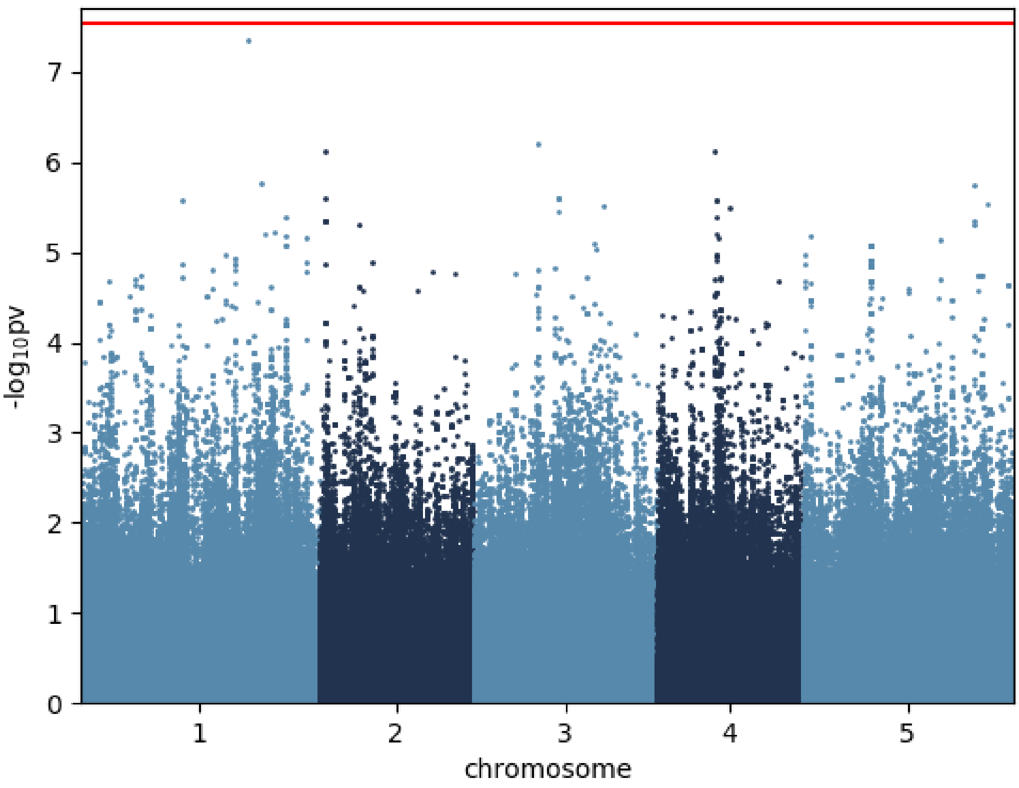


Leucine_6 °C

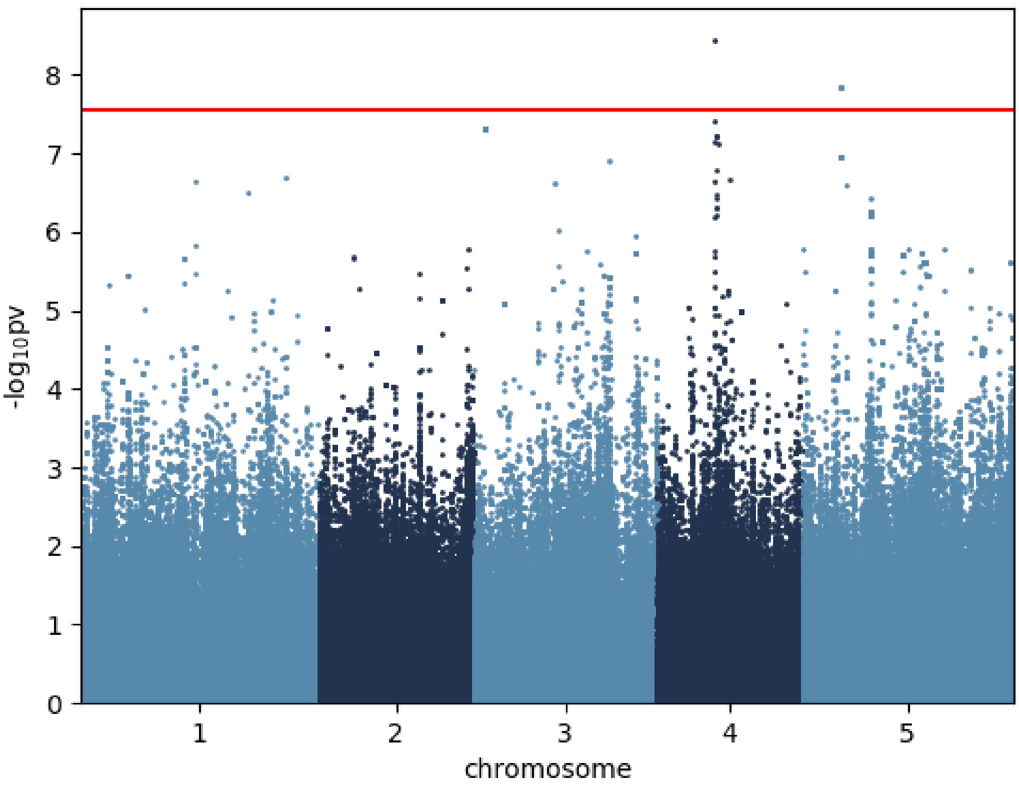


Lysine_6 °C

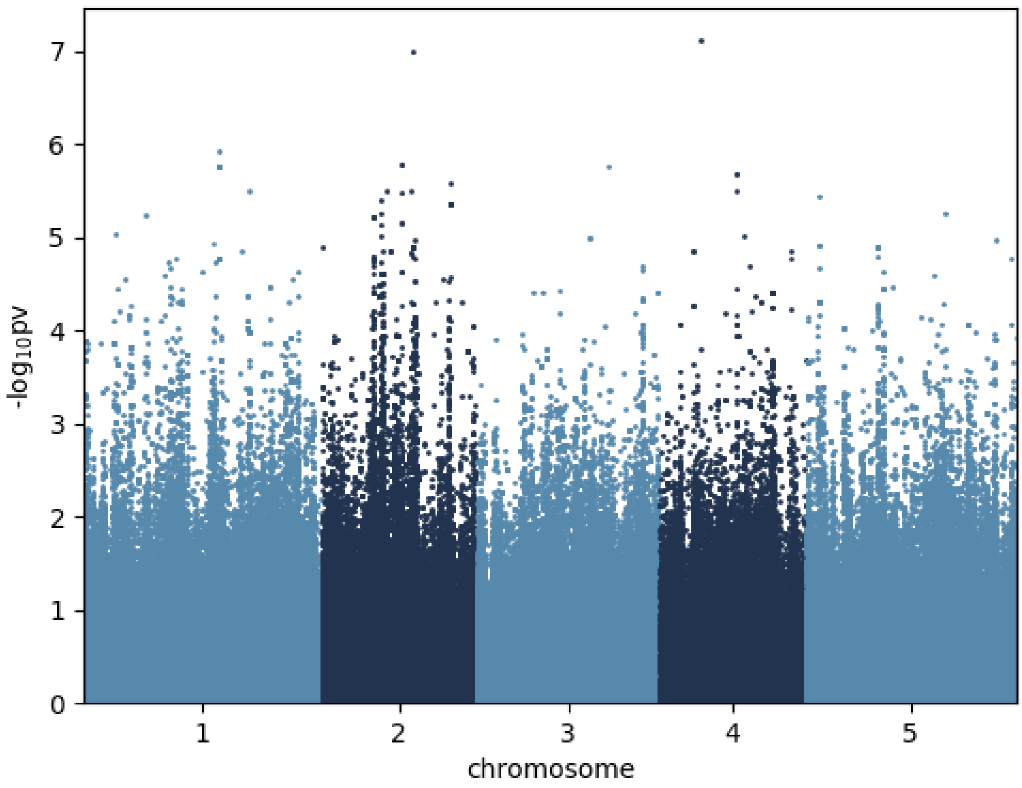


Malic.acid_6 °C

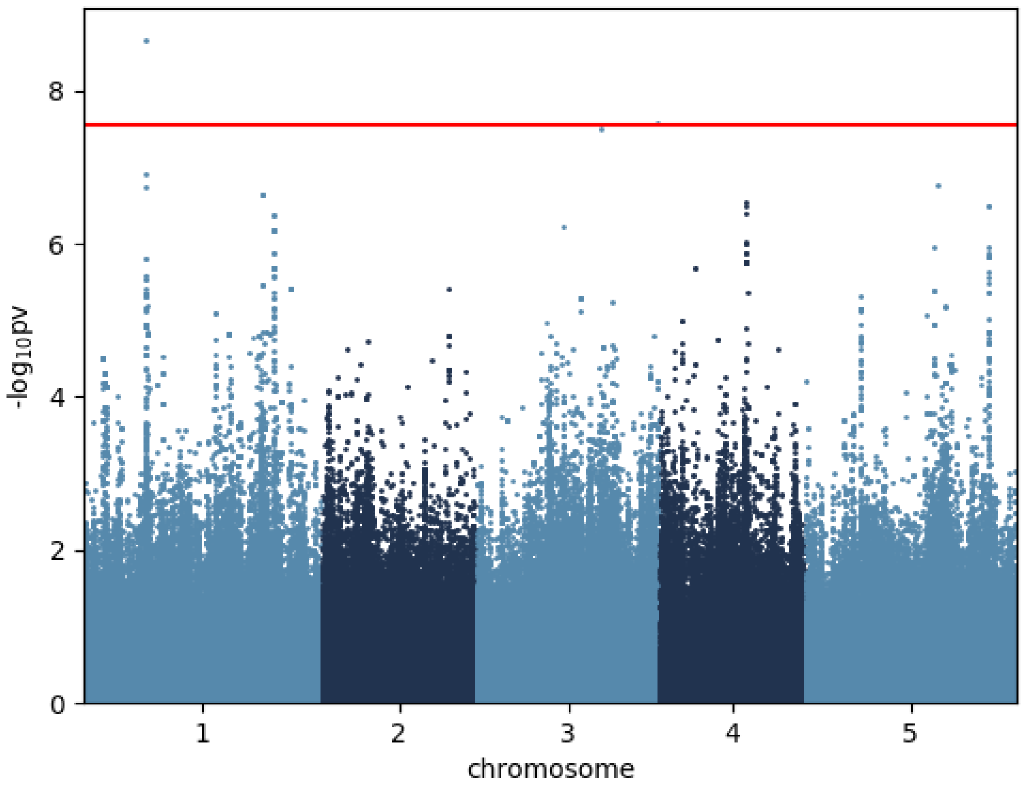


Maltose_6 °C

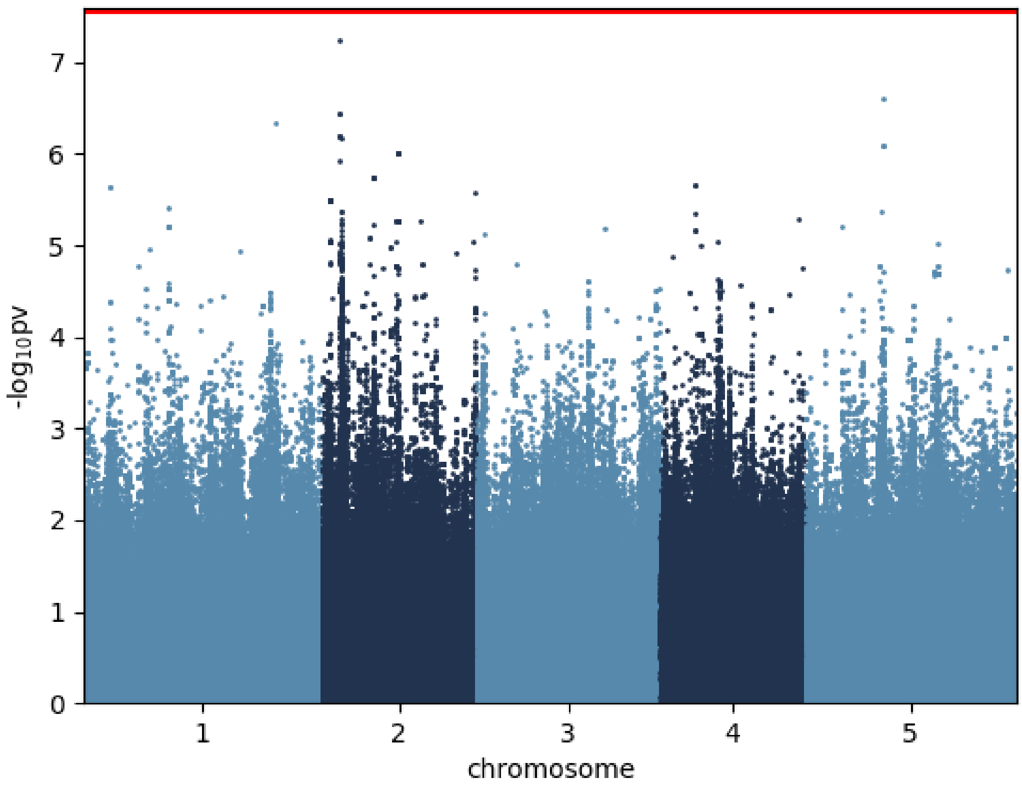


Myo.Inositol_6 °C

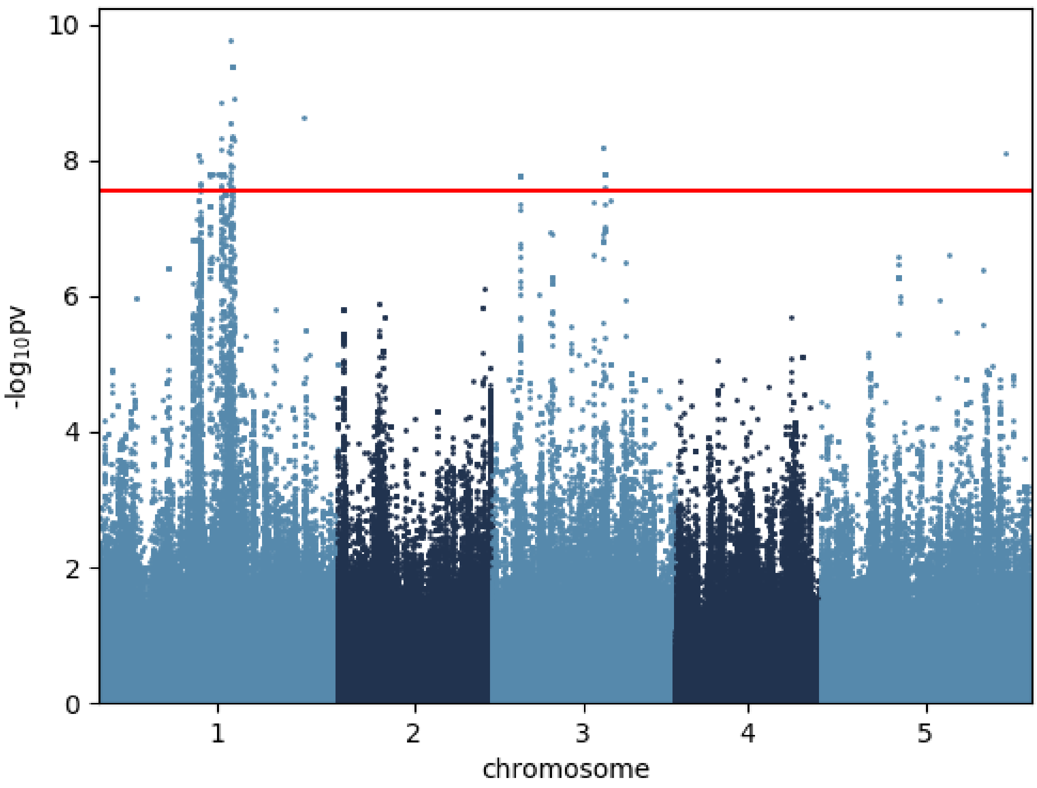


Ornithine_6 °C

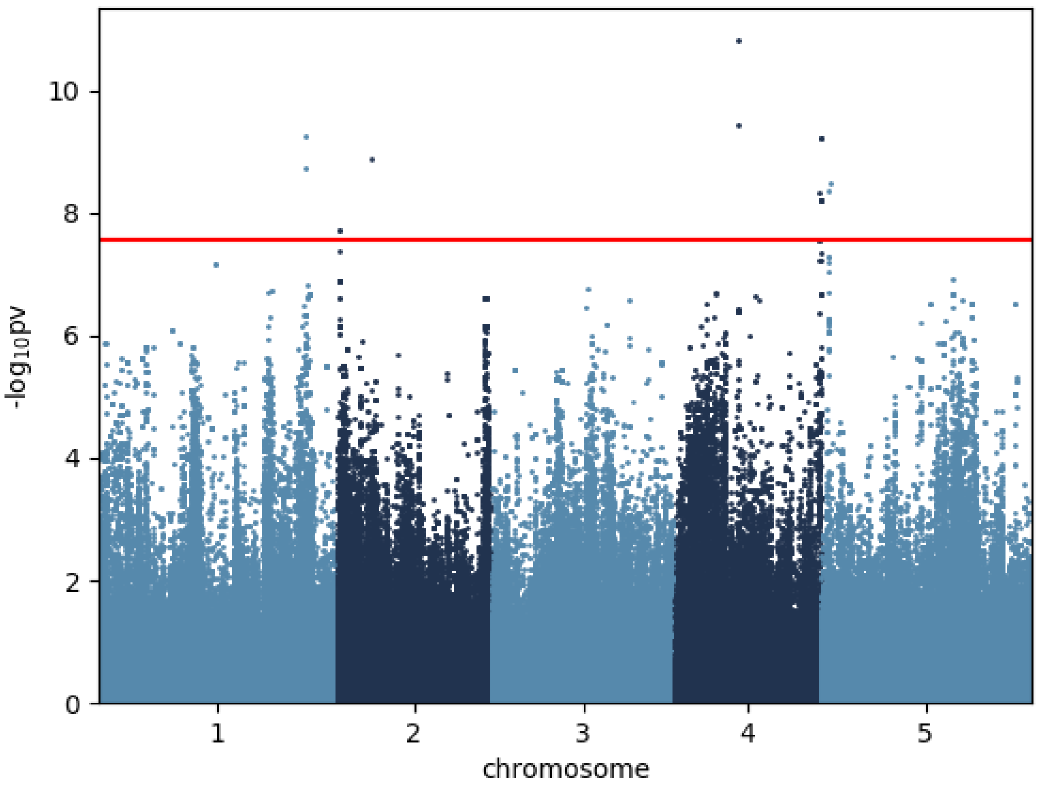


Oxoglutaric.acid_6 °C

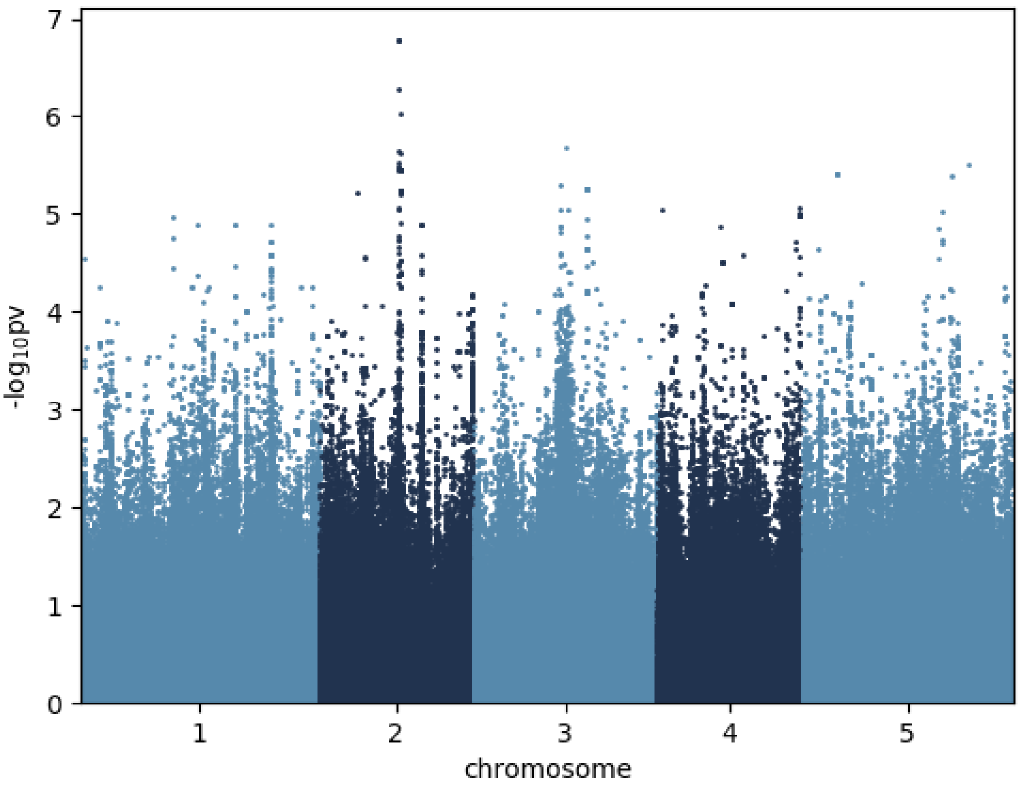


Phenylalanine_6 °C

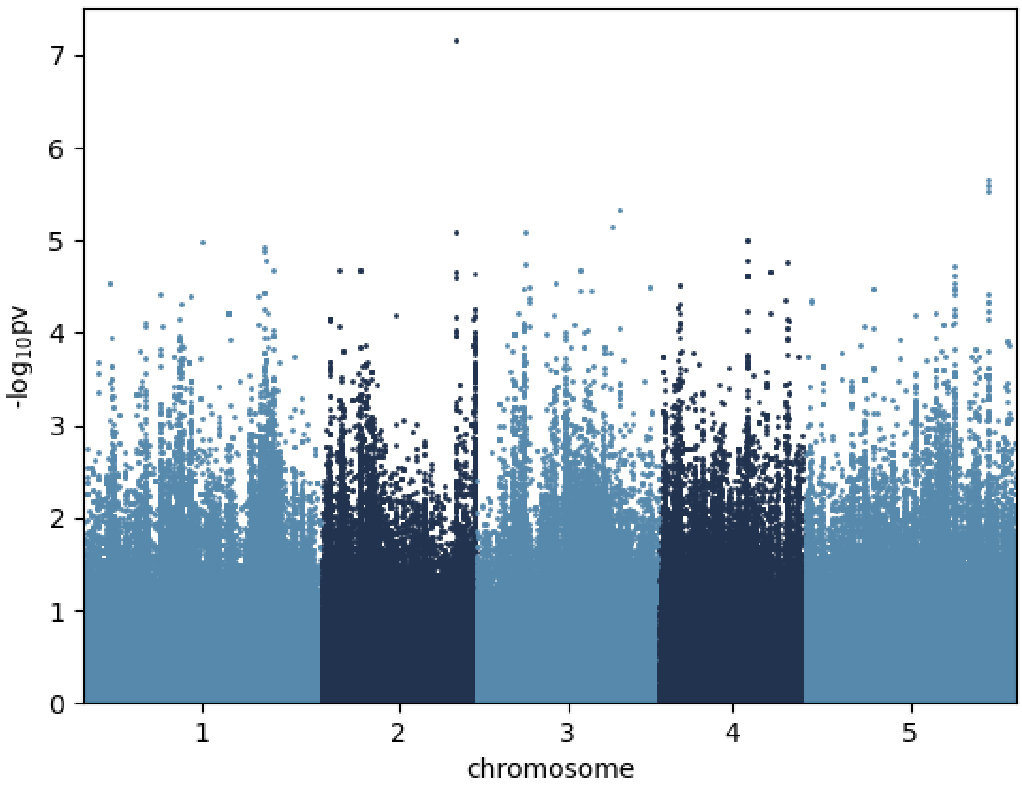


Proline_6 °C

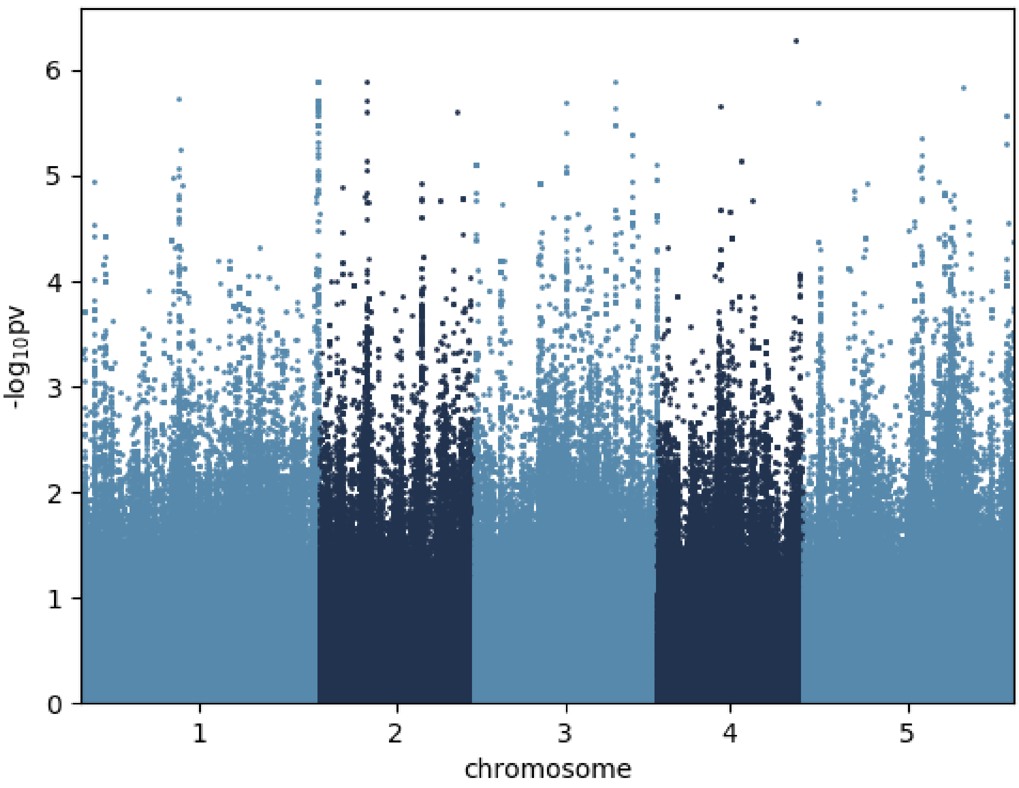


Putrescine_6 °C

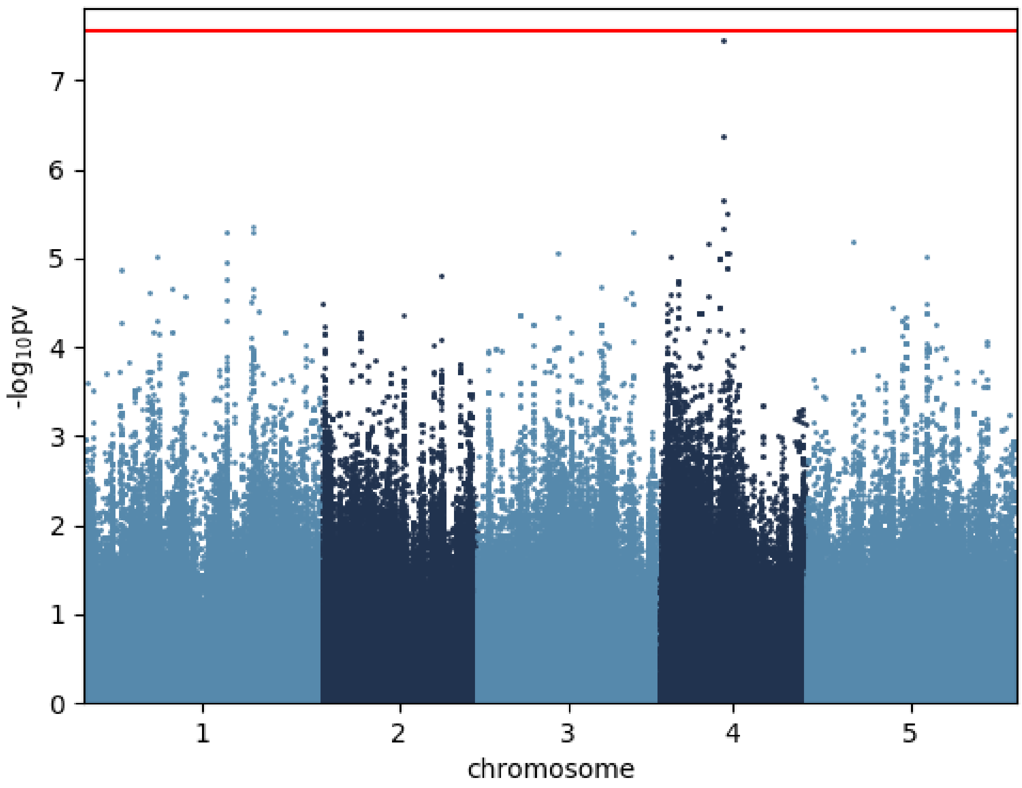


Pyruvic.acid_6 °C

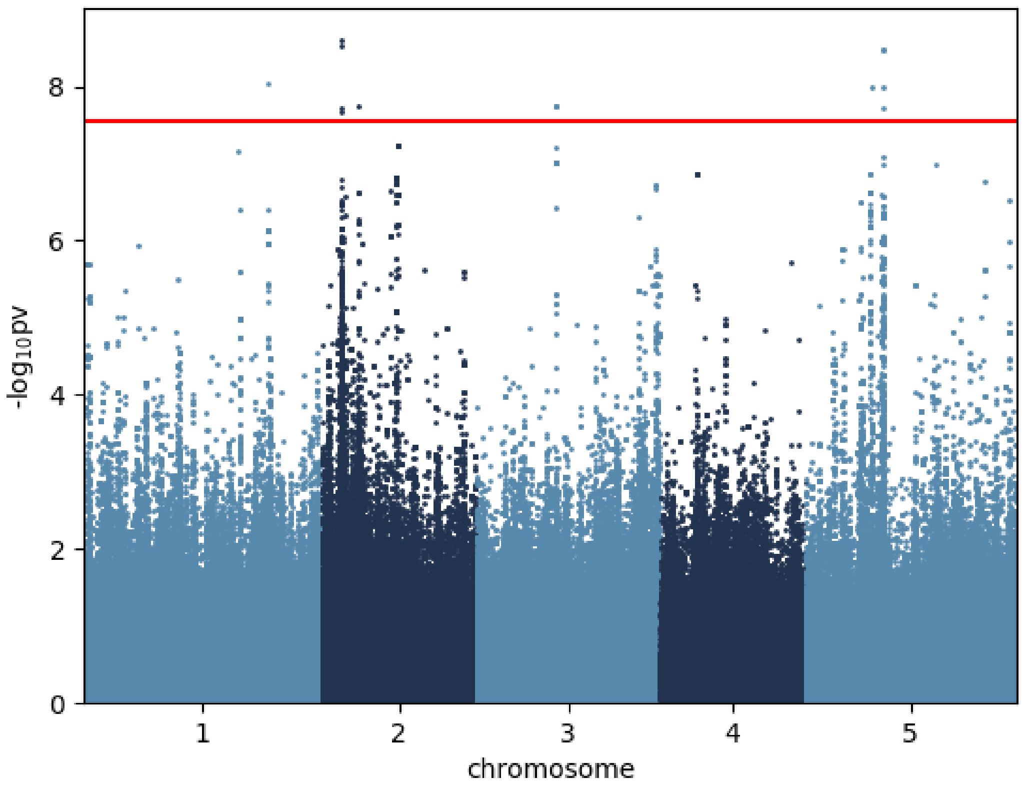


Raffinose_6 °C

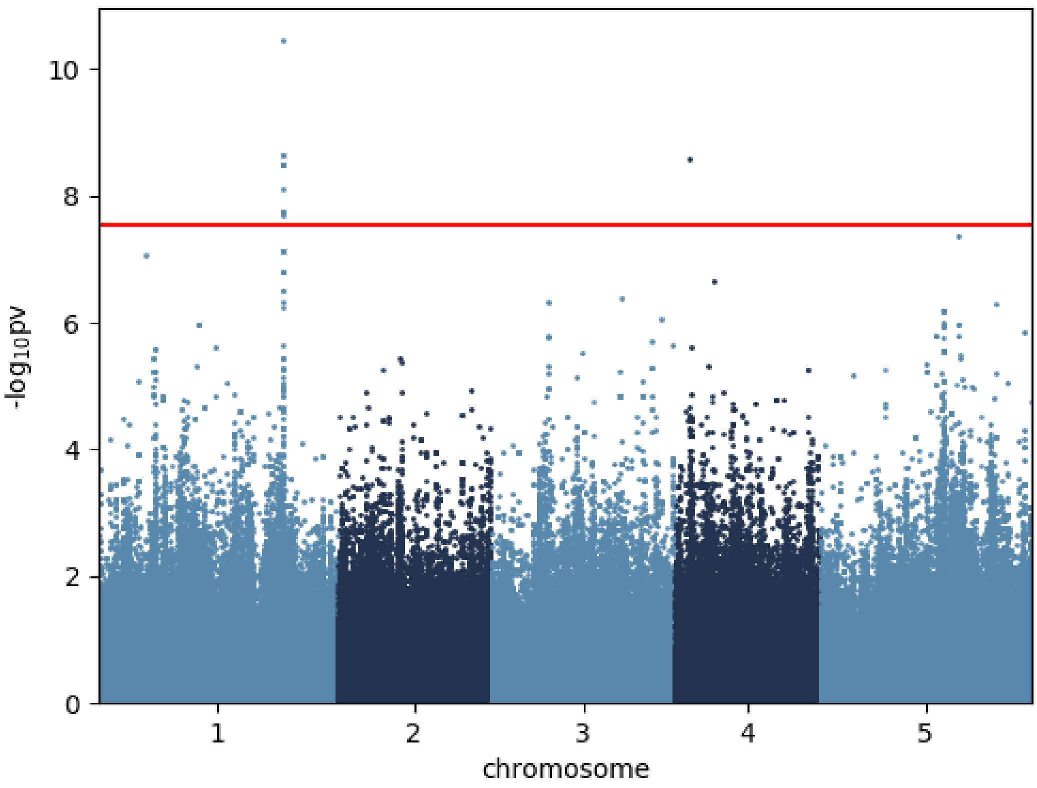


Serine_6 °C

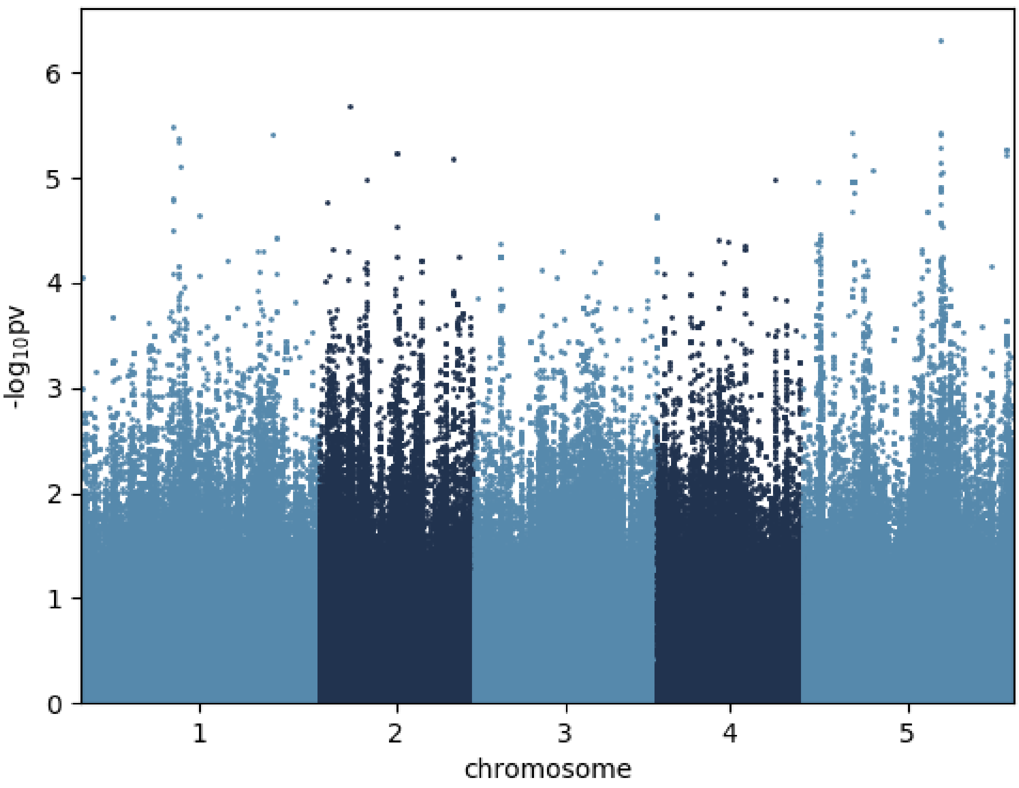


Spermidine_6 °C

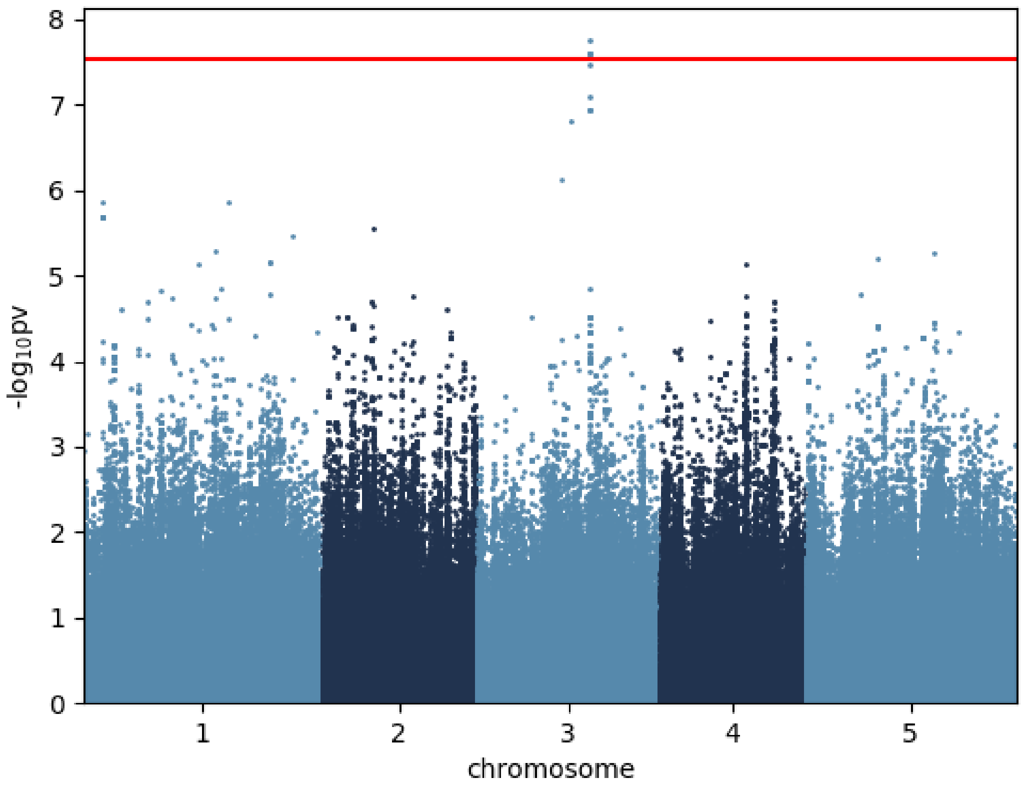


Succinic.acid_6 °C

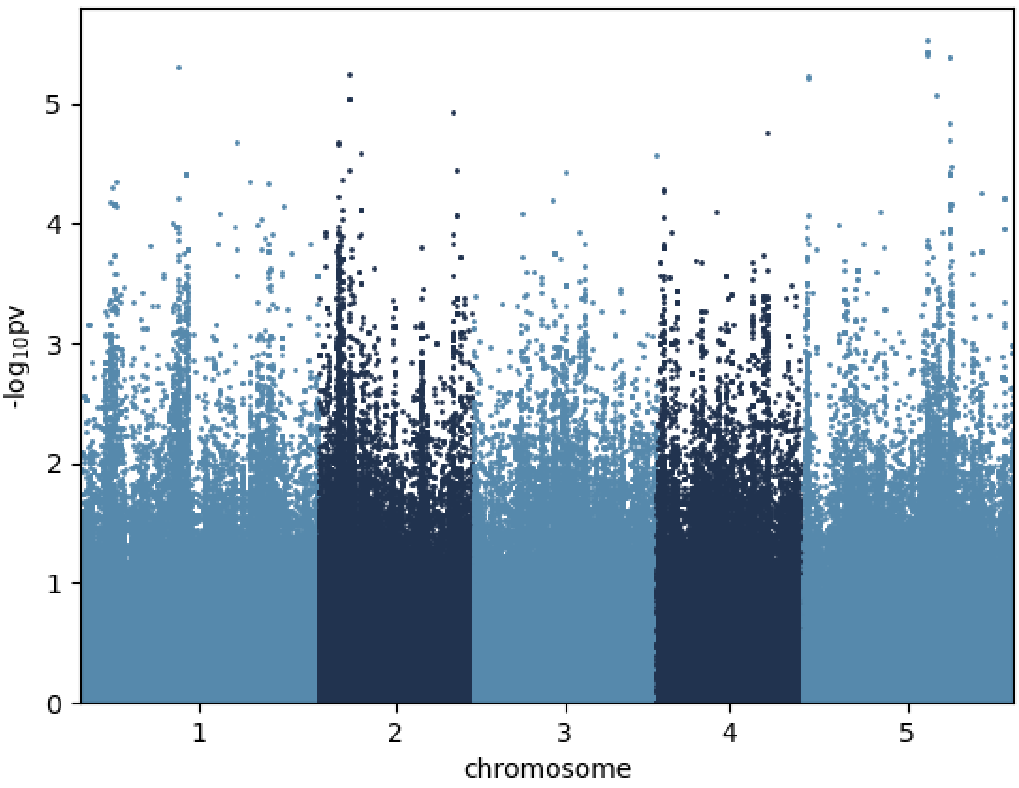


Sucrose_6 °C

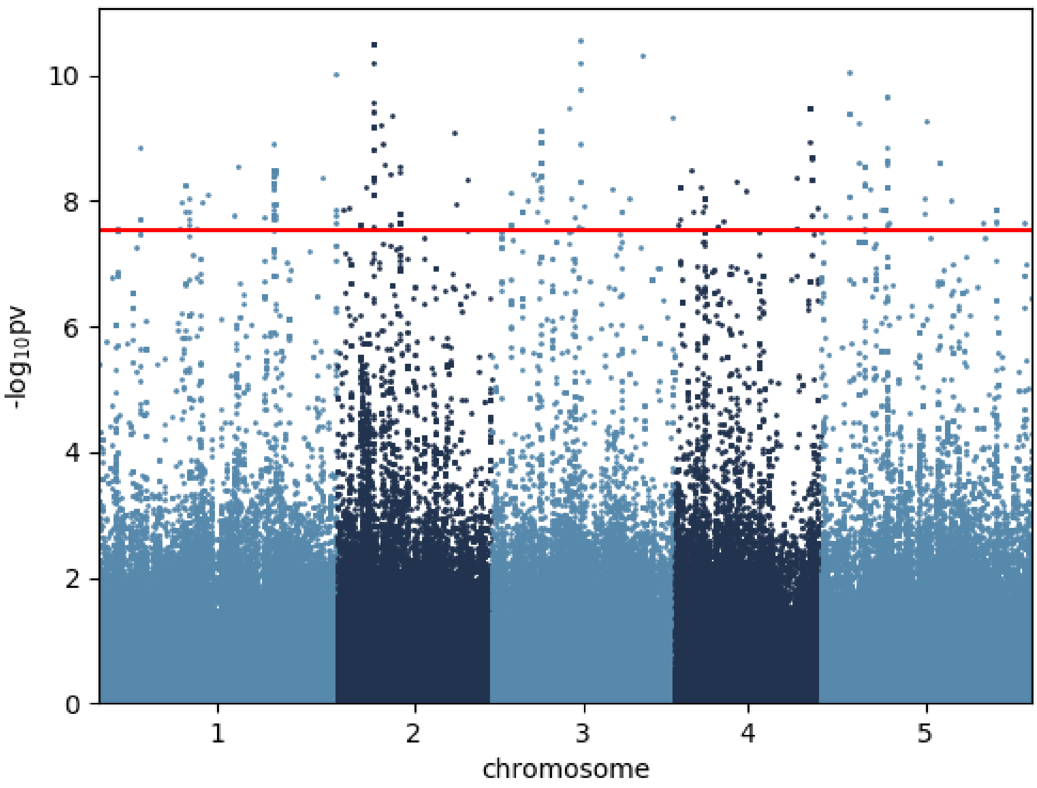


Threitol_6 °C

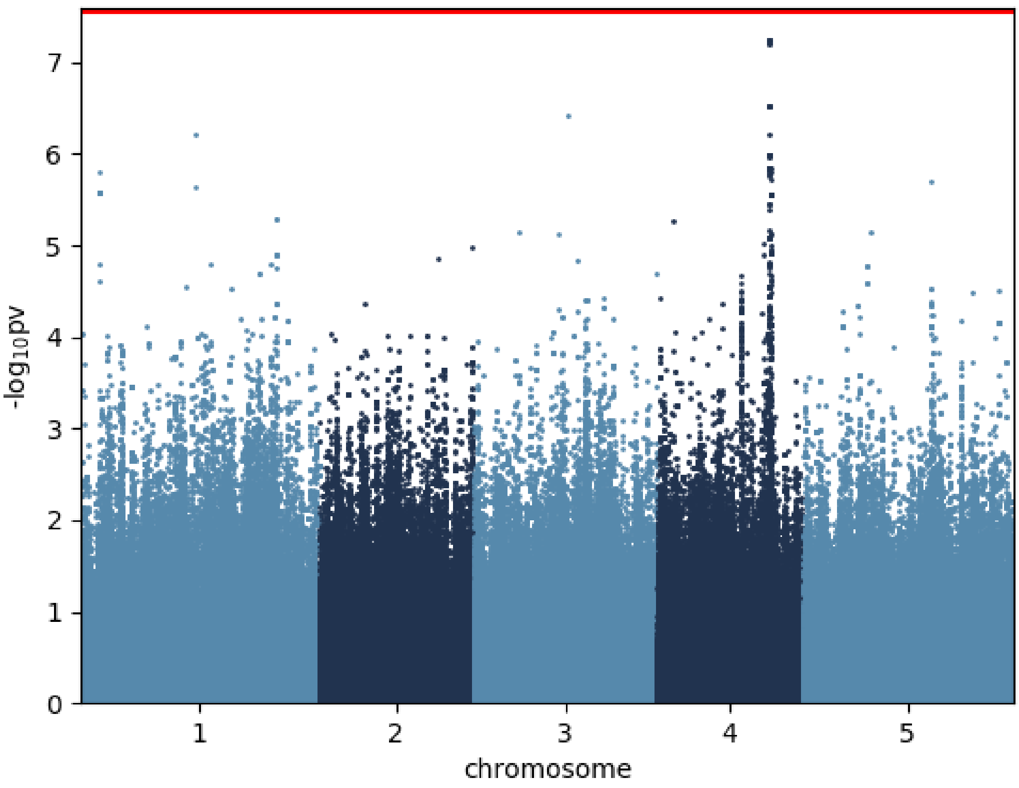


Threonic.acid_6 °C

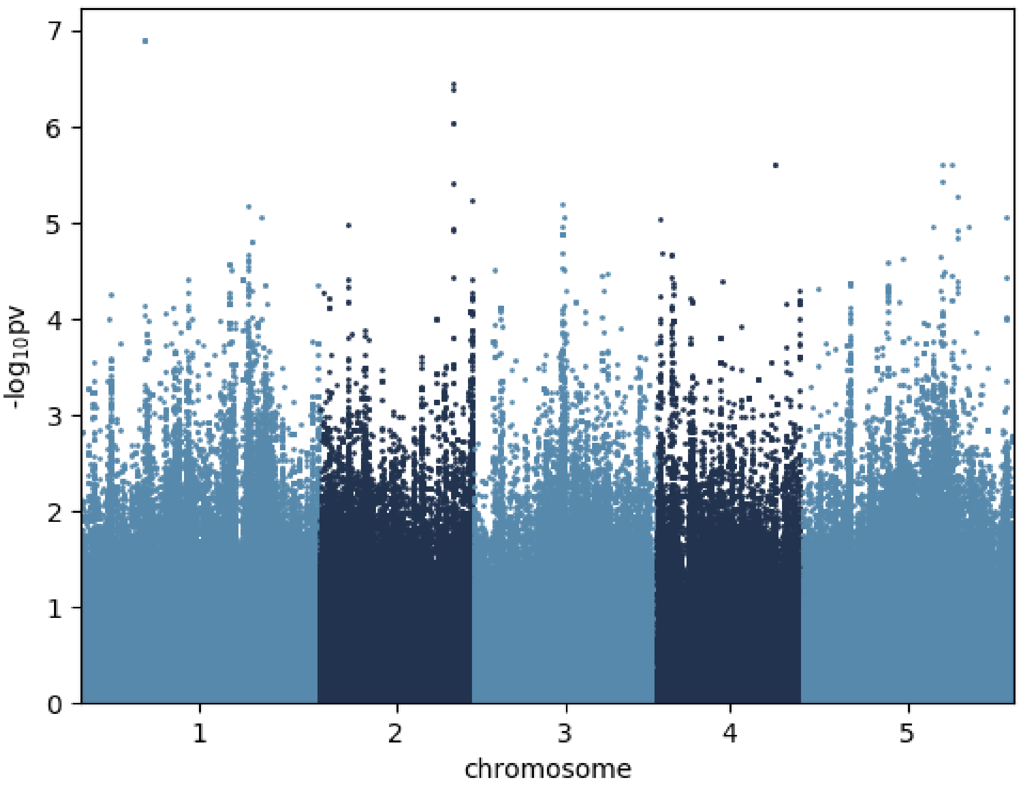


Threonine_6 °C

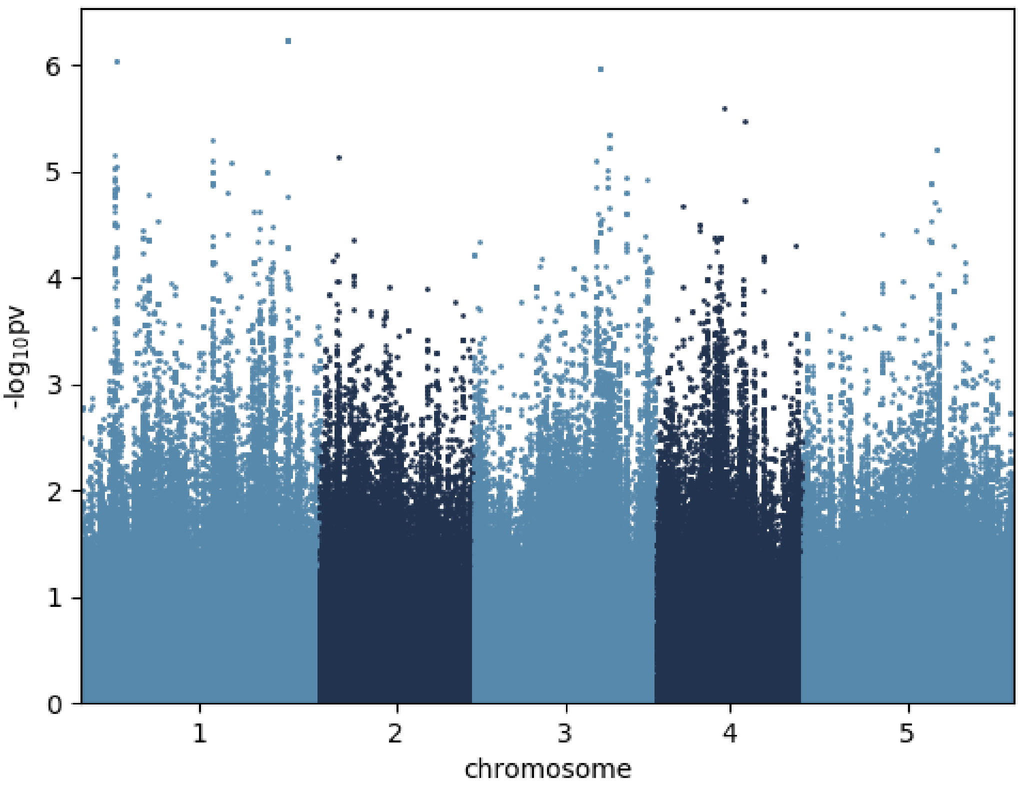


Trehalose_6 °C

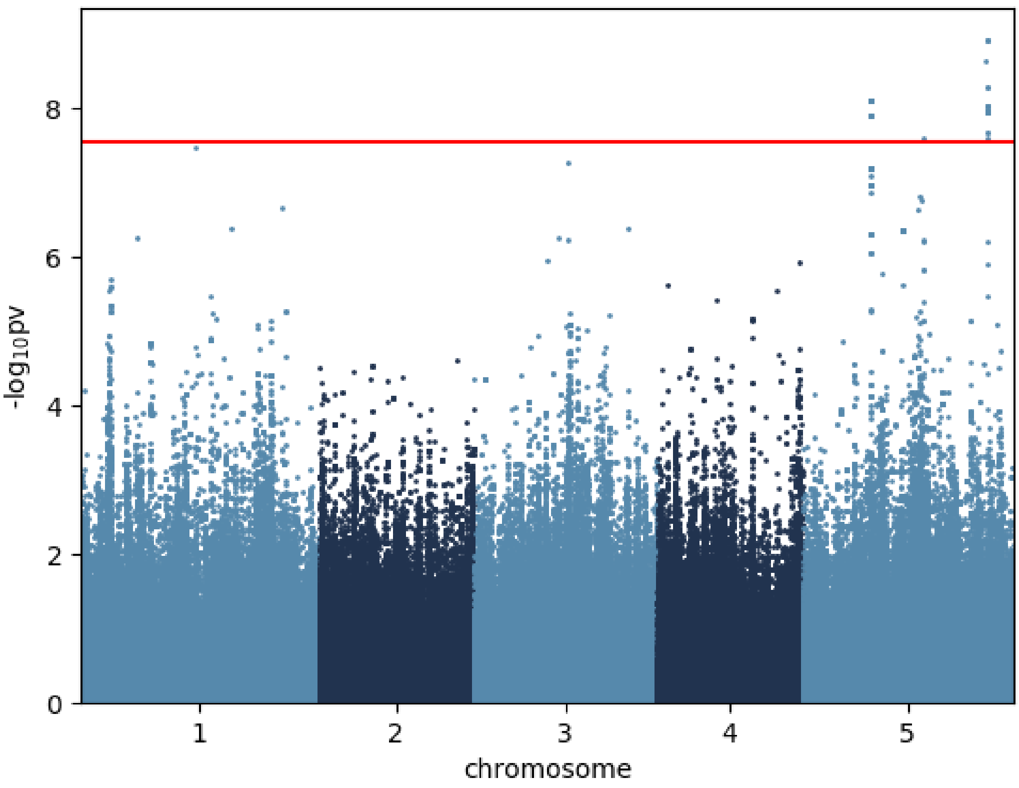


Tyrosine_6 °C

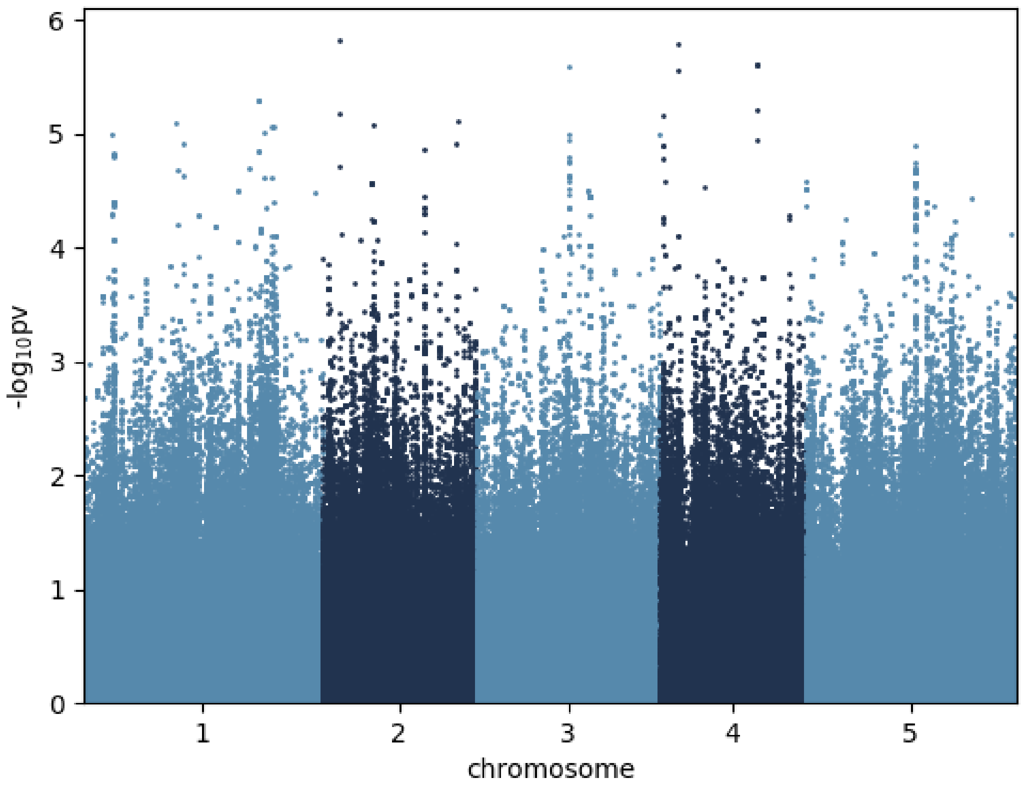


Valine_6 °C
